# The evolution of condition-dependent self-fertilisation

**DOI:** 10.64898/2026.02.15.705963

**Authors:** Thomas Lesaffre, John R. Pannell, Charles Mullon

## Abstract

Self-fertilisation is common in cosexuals, but selfing rates vary among species, among populations and among individuals within populations. Most evolutionary theory seeking to explain this variation assumes genetically determined selfing rates. Here we study the evolution and consequences of condition-dependent selfing, where individuals adjust their selfing rate in response to their deleterious mutation load. We analyse a two-locus population-genetic model in which one locus determines condition and the other is a modifier locus that determines condition-dependent selfing rates, and then extend the analysis to a polygenic background in which condition is determined by many loci. Our results show that selection favours positive condition dependence: high-condition individuals self-fertilise whereas low-condition individuals outcross. The resulting reaction norm generates stable within-population variation in realised selfing rates at evolutionary equilibrium and reduces the mutation load. We further show that it persists under environmental heterogeneity, and that pollen discounting favours a gradual increase in selfing with condition, leading to a continuum of selfing phenotypes. Altogether, our results indicate that condition-dependent selfing can generate substantial within-population variation in selfing rates. It may therefore contribute to mating-system diversity, and in particular to the maintenance of mixed mating.

## 1 Introduction

Most plants and many animals are cosexual, meaning that individual organisms carry both male and female sexual functions. In those with an ability to self-fertilise, the selfing rate varies widely, from predominantly selfing species to strictly outcrossing ones, with many showing mixed mating in which individuals self-fertilise at an intermediate rate (Barrett and Harder, 1996; Goodwillie et al., 2005; Jarne and Auld, 2006; Barrett and Harder, 2017a; Whitehead et al., 2018). This variation occurs not only among but also within species, where different populations often exhibit different selfing rates (Whitehead et al., 2018), and within population, where individuals can differ substantially in their realised selfing rates owing to environmental and morphological differences (Holtsford and Ellstrand, 1992; Herlihy and Eckert, 2007; Eckert et al., 2009; Ferriol et al., 2011; Nora et al., 2016; Brunet and Eckert, 1998; Karron et al., 1997; Chen and Pannell, 2024). Because these patterns determine how genes are transmitted across generations, explaining the evolutionary mechanisms that shape selfing-rate diversity within and between populations has been a longstanding problem in evolutionary biology (Darwin, 1877; Charlesworth and Charlesworth, 1979; Barrett, 2003).

Self-fertilisation comes with major ecological benefits, as it assures reproduction when mates or pollinators are scarce and avoids the costs and risks associated with mate searching (Chap. 1 in Darwin, 1877; Jarne and Charlesworth, 1993; Kalisz et al., 2004; Eckert et al., 2006; Sicard and Lenhard, 2011). It also carries an inherent transmission advantage: a selfing parent acts as both mother and father and so transmits twice as many gene copies per descendant as it would through outcrossing (Fisher, 1941). Against these advantages stands the cost of inbreeding depression, the reduction in fitness of selfed relative to outcrossed offspring caused by the expression of recessive deleterious mutations segregating in populations (the ‘mutation load’, Charlesworth and Charlesworth, 1987, 1999; Charlesworth and Willis, 2009). According to classical theory, the interaction between inbreeding depression and the advantages of selfing leads to a threshold effect in mating-system evolution (Lande and Schemske, 1985). In outcrossing populations, recessive deleterious mutations are held at low frequencies and so are almost always found in heterozygous state. Selfing increases homozygosity and exposes their effects, which reduces the fitness of selfed offspring (Charlesworth and Charlesworth, 1987), but also makes the purging of mutations more effective. This generates a positive feedback for the evolution of selfing, because purging reduces inbreeding depression and therefore strengthens selection for increased selfing. As a result, theory predicts that complete outcrossing should be maintained when inbreeding depression is above a threshold (strong enough to overcome the benefits of selfing), and complete selfing should evolve when it is below, so that mixed mating is unstable (Lande and Schemske, 1985; Charlesworth et al., 1990, 1991; Lande and Porcher, 2015; Abu Awad and Billiard, 2017; Abu Awad and Roze, 2018, 2020).

Much work has since built on this seminal insight to explain the widespread occurrence of mixed mating in cosexual species (about 40% have selfing rates between 0.2 and 0.8 according to surveys in plants and animals, although the frequency of selfing plant species may be overestimated owing to a study bias in their favour; Goodwillie et al., 2005; Jarne and Auld, 2006; Igic and Kohn, 2006; Meyer et al., 2025). Several studies have shown that additional mechanisms relevant to the dynamics of mating can contribute to maintaining mixed mating under certain conditions (Goodwillie et al., 2005), notably limited pollen and seed dispersal (Ronfort and Couvet, 1995), variable pollination conditions (Morgan and Wilson, 2005) and pollen discounting – the reduction in pollen export that can accompany increased selfing (Holsinger et al., 1984; Holsinger, 1991; Harder and Wilson, 1998; Johnston, 1998; Porcher and Lande, 2005). Other studies considered more elaborate biological scenarios for the effects of inbreeding depression, allowing it to vary, e.g., between female and male components of fitness (Rausher and Chang, 1999), as a function of population density (Cheptou and Dieckmann, 2002), or of time and space (Cheptou and Mathias, 2001). These models show that strong differences in inbreeding depression across contexts can also select for mixed mating. However, their conclusions rely on the assumption of fixed inbreeding depression and so ignore purging, a major force in mating-system evolution in the face of which wide within-population variation in the magnitude of inbreeding depression may not be readily maintained (Porcher et al., 2009).

Most theory on mating-system evolution assumes that the selfing rate is under genetic control, so that all individuals in the population have the same selfing rate at evolutionary equilibrium (see Jarne and Charlesworth, 1993; Goodwillie et al., 2005; Clo et al., 2025 for reviews). Yet, individuals may vary substantially in their selfing rates (Ferriol et al., 2011; Nora et al., 2016; Brunet and Eckert, 1998; Karron et al., 1997; Whitehead et al., 2018). This variation could arise as a result of stochastic effects in plant-pollinator interactions (Kalisz et al., 2004; Richards et al., 2009; Barrett and Harder, 2017b). However, it could also arise through a form of plasticity in which individuals adjust their mating behaviour according to environmental or physiological cues (Levin, 2010; Suijkerbuijk et al., 2025). Plastic responses of various kinds are common in plants, including in mating-related traits such as herkogamy or the timing of autonomous selfing, which have been shown to vary with environmental conditions (Vallejo-Marín and Barrett, 2009; Levin, 2010; Jorgensen and Arathi, 2013; Spigler, 2017; Barrett and Harder, 2017b; Suijkerbuijk et al., 2025). One form of plasticity leading to inter-individual differences in selfing that has received attention is delayed selfing, where individuals self-fertilise only when outcrossing fails (Lloyd, 1979, 1992; Goodwillie and Weber, 2018). This plastic response is favoured in uncertain pollination environments, where the benefits of selfing vary between individuals (Tsitrone et al., 2003; Morgan and Wilson, 2005). Much less attention has been paid to the possibility that the costs of selfing may also vary among individuals in a population, simply because individuals differ in the number of deleterious mutations that they carry. In this context, a form of plasticity where individuals adjust their mating behaviour to their individual condition – an individual’s overall vitality given its genetic background and environment – could also be favoured. For example, the degree of herkogamy or the timing of anther dehiscence relative to pistil receptivity within flowers, both of which have been shown to affect individual selfing rates (Holtsford and Ellstrand, 1992; Kalisz et al., 2012; Koski et al., 2018; Opedal, 2018), could be plastically tuned in response to condition.

The notion that mating-related traits are expressed plastically in response to individual condition is well-established in dioecious animals (Rowe and Houle, 1996; Jennions et al., 2001; Fricke et al., 2009; Flintham et al., 2023). It relies on the idea that secondary sexual traits are costly to produce and maintain. Individuals in good condition can afford to express larger ornaments, stronger courtship displays or more competitive behaviours, whereas those in poor condition cannot. This makes such traits reliable indicators of genetic quality, which the other sex (often females) can use when choosing mates to obtain indirect genetic benefits, as their offspring inherit alleles associated with higher condition (Rowe and Houle, 1996; Pomiankowski and Iwasa, 2001; Van Doorn and Weissing, 2006, see also Tonnabel et al., 2021 for a review of how similar processes might unfold in plants). Condition dependence also plays a central role in models of plastic switches between sexual and asexual reproduction. In these models, selection favours modifier alleles that increase the rate of sex in low-fitness genotypes, because recombination then allows the modifier to escape such deleterious genetic backgrounds (Hadany and Otto, 2007; Otto, 2009). This “abandon-ship” mechanism favouring condition-dependent sex could also play a role in the evolution of condition-dependent self-fertilisation, with outcrossing as an escape device. However, the idea that an individual’s selfing rate might be a plastic response to its condition, particularly its genetic condition, appears not to have been considered.

In this paper, we investigate the evolution of condition-dependent selfing using a combination of mathematical modelling and individual-based simulations. As a baseline, we analyse a two-locus model in which one locus affects condition and the other controls genotype-specific selfing rates, to ask whether selection favours individuals in different genetic conditions to self-fertilise at different rates. Because inbreeding depression is thought to arise from deleterious mutations at many loci, we then consider a more realistic polygenic case in which condition depends on mutations at numerous loci, and the selfing rate of each individual is determined by a gene-regulatory network that can evolve to respond to condition (Wagner, 1994; Ezoe and Iwasa, 1997; Deshpande and Fronhofer, 2022, 2025). This approach lets us examine the reaction norm linking condition to selfing that evolves, while placing minimal constraints on its shape.

In the last part of the paper, we examine how two additional factors might modify the evolution and stability of condition-dependent selfing. The first is the effect of the environment on condition. Individual condition not only reflects genetic differences but also environmental variation in factors such as resource availability, competition, and exposure to pests and diseases (Wilson and Nussey, 2010; Forsythe et al., 2021). When condition-dependent selfing relies on condition as a cue to genetic condition, environmentally induced variation can blur the association between condition and genotype at loci affecting condition, which raises the question of whether environmental effects on condition could weaken, modify or perhaps even destabilise condition-dependent differences in selfing rate within populations. The second factor is the effect of pollen discounting on the balance between selfing and outcrossing. Pollen discounting measures the extent to which self-fertilisation reduces an individual’s success as a pollen donor (Harder and Wilson, 1998). This can occur when pollen that could have sired outcrossed progeny is instead deposited on the same plant, either within flowers or via geitonogamous transfer among flowers (de Jong et al., 1993; Barrett, 2002), or when changes in floral morphology that promote selfing, for instance reductions in herkogamy, compromise pollen export (Harder and Barrett, 1995). Pollen discounting imposes an additional mating cost to selfers that scales with their selfing rate. In doing so, it selects against self-fertilisation and can favour mixed mating under some scenarios (Johnston, 1998). In a condition-dependent scenario, pollen discounting may affect how the benefits of selfing vary with condition, and could therefore influence the shape of the reaction norm.

## 2 Condition-dependent self-fertilisation is favoured as an escape strategy

Most of the theory presented below should apply generally to any cosexual plant or animal species with an ability to self-fertilise (Goodwillie et al., 2005; Jarne and Auld, 2006). However, we frame our models explicitly in terms relevant to plants, both for brevity and because it allows us to situate our work more clearly within the field, as most empirical and theoretical research on the evolution of selfing focuses on this group.

### 2.1 Two-locus model

We consider a large population of diploid cosexuals characterised by their genotype at two loci (Fig. 1). The first locus affects condition with two alleles: a wild-type allele ‘A’ and a deleterious allele ‘a’. The condition of individuals with genotype AA, Aa and aa at this locus, which we refer to as the ‘condition locus’ hereafter, is given by *φ*_AA_ = 1, *φ*_Aa_ = 1 − *sh* and *φ*_aa_ = 1 − *s*, respectively, where *s* is the deleterious effect of allele a on condition and *h* is its dominance coefficient. During meiosis, the wild-type allele A mutates into deleterious allele a with probability *µ*_A_ and the deleterious allele a mutates into wildtype allele A with probability *µ*_a_, so that the locus remains polymorphic at a balance between mutation and selection, and individuals in the population vary in their condition. The second locus is a selfing modifier at which alleles are additive and pleiotropically encode three genotype-specific selfing rates, ***α*** = (*α*_AA_, *α*_Aa_, *α*_aa_) ∈ [0, 1]^3^, which express conditionally on genotype at the condition locus (e.g., *α*_AA_ is the selfing rate expressed in a AA homozygote). In this model, organisms are therefore assumed to plastically respond to their condition directly, and condition-dependent selfing occurs whenever genotype-specific selfing rates differ. These rates each evolve via the input of rare and independent mutations of small and unbiased phenotypic effects (“continuum-of-alleles” model; Kimura, 1965).

**Figure 1:**
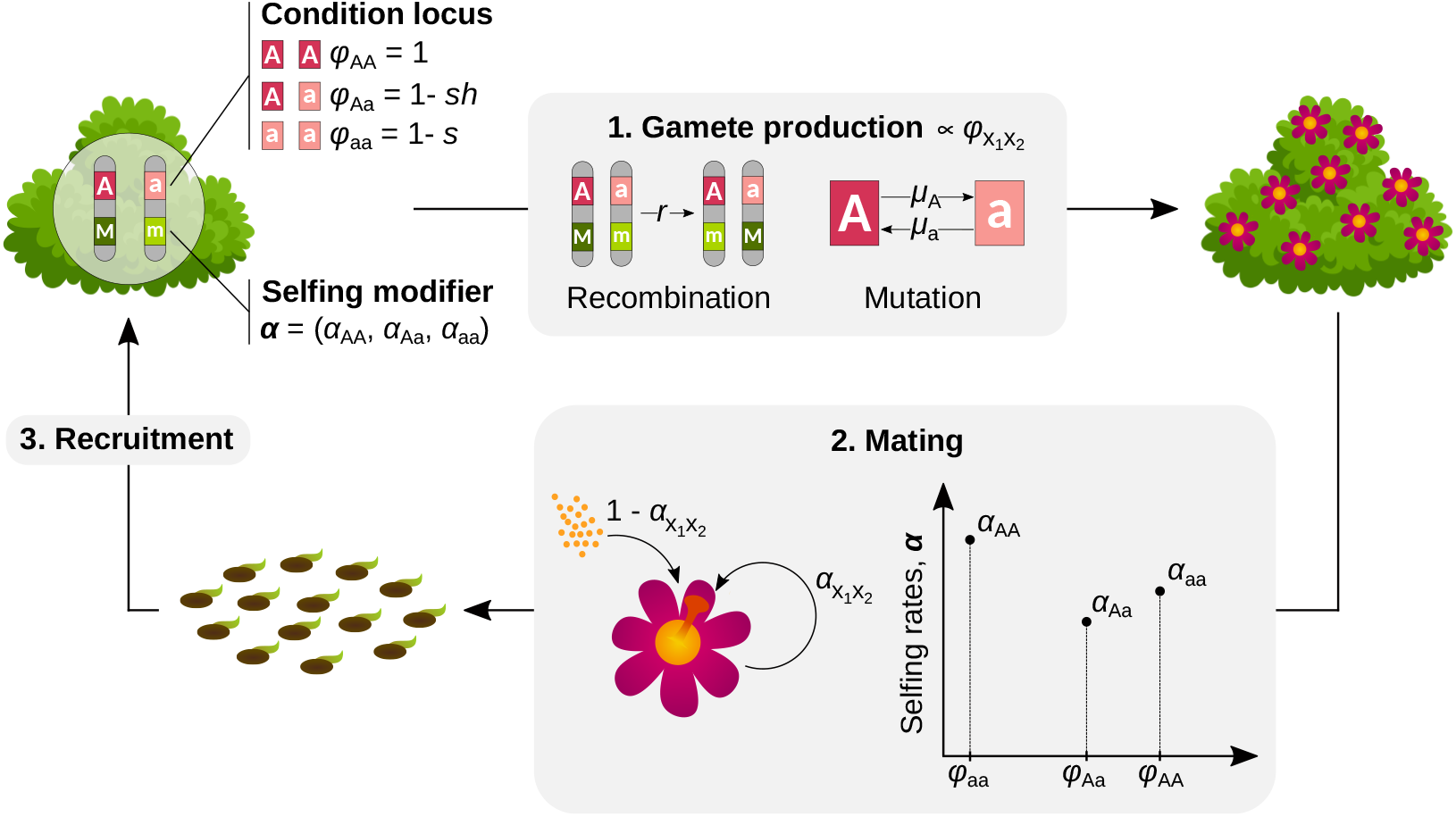
Life cycle and genetic bases of condition and selfing in the two-locus model. Individuals are characterised by their genotype at the condition locus (in red) and the selfing modifier (in green). They first produce gametes in proportion to their condition, at which point recombination takes place between the two loci and mutation occurs at the condition locus. Individuals then self-fertilise at a rate that depends on their genotype at both loci, and mate randomly for the remaining ovules. Following seed production, adults die and the next generation is recruited from the produced seeds. See text for more detail.

Each generation, individuals pass through the following life cycle events. (1) *Gamete production:* They first produce a large number of female and male gametes proportionally to their condition 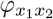, where *x*_1_, *x*_2_ ∈ {A, a} denote the two alleles carried by the individual at the condition locus (we describe the effect of alleles at this locus as affecting the number of female and male gametes produced by individuals in the text for simplicity, but differences in contribution to the gamete pools owing to condition could also stem from differences in individual survival to adulthood or in the quality, rather than quantity of gametes produced; the modelling would be unchanged). The two loci recombine at rate *r* ∈ [0, 1/2] and mutation occurs at the condition locus at a rate *µ*_*x*_, *x* ∈ {A, a}, per allele during gametogenesis. (2) *Mating:* Individuals then self-fertilise a fraction 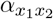 of their ovules, which depends on their genotype at the modifier and on their condition. The remaining fraction of ovules 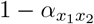 is fertilised via random mating. We assume that there is enough pollen exchanged for all ovules to be fertilised (i.e., no pollen limitation), and that self-fertilising does not reduce an individual’s contribution to the outcross pollen pool (i.e., no pollen discounting – we later relax this assumption, see Fig. 5 in section 3.4 below). (3) *Recruitment:* Once all ovules have been fertilised, seeds mature, adults die and the next generation is sampled from the produced seeds, so that the population remains of large and constant size.

### 2.2 Effect of condition-dependent selfing on the condition locus

We start by fixing the relationship between selfing and condition to investigate how condition dependence influences the equilibrium frequency of the deleterious allele *p*^∗^ and resulting mean condition *φ*^∗^ in the population.

#### Linear condition dependence

In order to obtain explicit mathematical results, let us assume a linear relationship between condition and selfing, with genotype-specific selfing rates given by

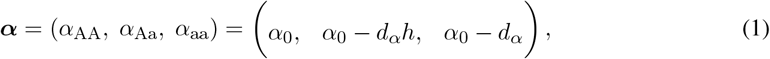

where *α*_0_ ∈ [0, 1] is the selfing rate of wild-type homozygotes, and *d*_*α*_ ∈ [−*α*_0_, 1 − *α*_0_] gives the strength of condition dependence (where the dominance coefficient *h* of the deleterious allele features in the selfing rate of heterozygotes to ensure that the selfing rate varies linearly with condition, see Fig. 2A). Selfing rates are independent of condition when *d*_*α*_ = 0, whereas the population experiences positive condition dependence for the selfing rate when *d*_*α*_ *>* 0 and negative condition dependence when *d*_*α*_ *<* 0 (Fig. 2A). Under this assumption, we show in Appendix A.1.2 that the equilibrium frequency of allele a can be expressed as

**Figure 2:**
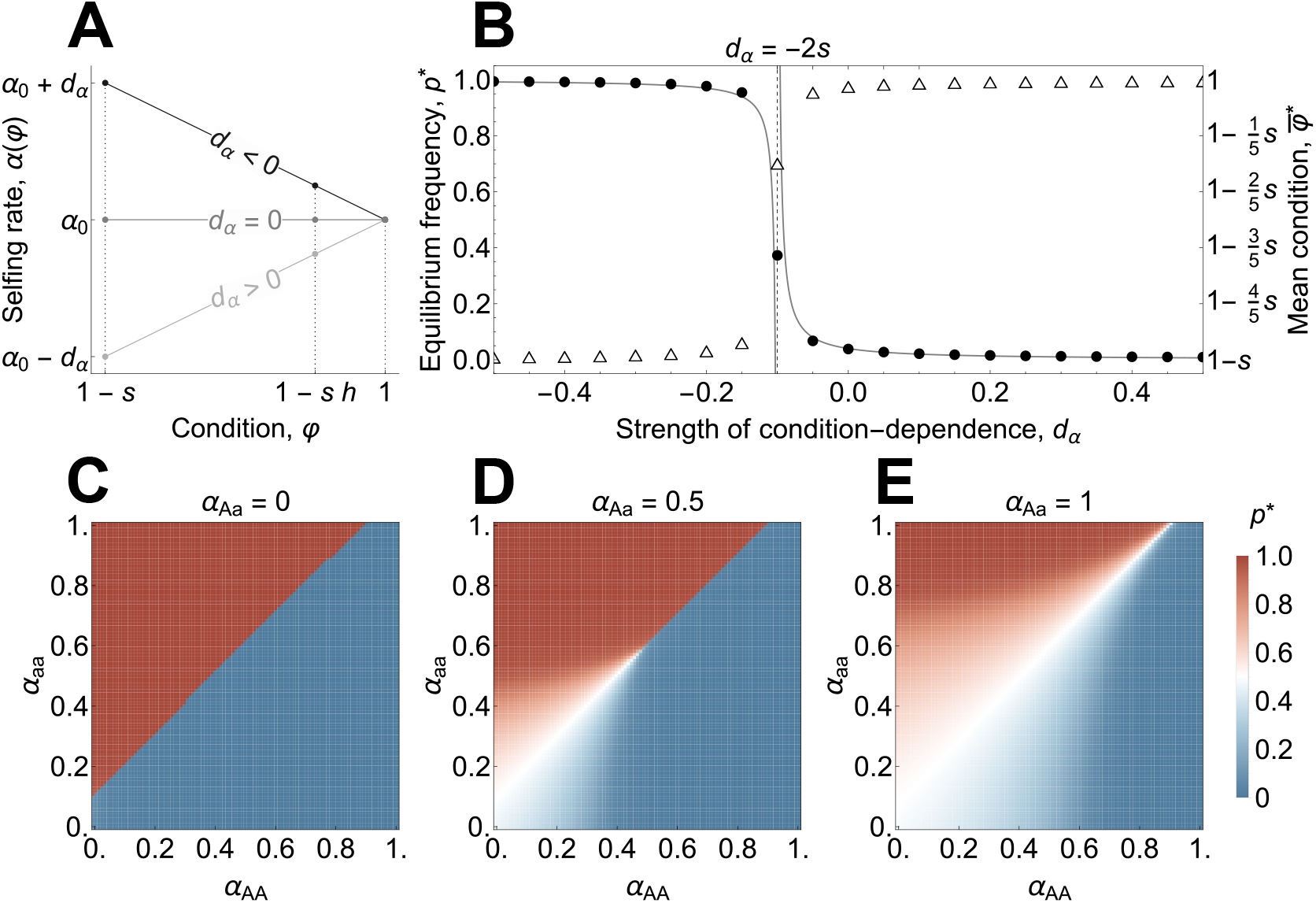
Effect of condition-dependent selfing on evolution at the condition locus. **A** Linear relationship between condition and the selfing rate, as assumed to compute eq. (2). *d*_*α*_ *<* 0 leads to negative condition dependence and *d*_*α*_ *>* 0 leads to positive condition dependence. **B** Equilibrium frequency of the deleterious allele a, *p*^∗^, and mean condition at this equilibrium, 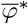, as a function of the strength of condition dependence, *d*_*α*_. Dots and triangles respectively indicate the equilibrium frequency and mean condition computed numerically without approximation. Lines indicate the analytical approximations for *p*^∗^ given in eqs. (2). Parameters specific to this plot: *α*_0_ = 0.5. **C-D-E** Equilibrium frequency *p*^∗^ as a function of the selfing rates of wild-type homozygotes (x-axis) and deleterious homozygotes (y-axis) at the condition locus in the general case, for three selfing rates of heterozygotes (*α*_Aa_ = 0, 0.5, 1 from left to right). Parameters used in all plots: *s* = 0.05, *h* = 0.25, *µ*_A_ = *µ*_a_ = 10^−3^.

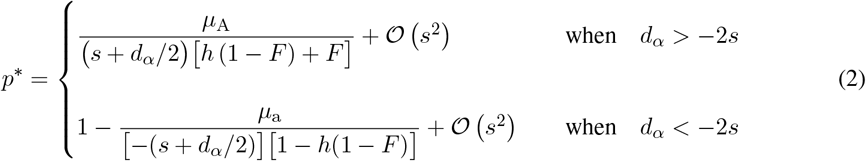

to first order in *s* (i.e., under weak variation in fitness), where

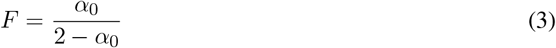

is the expected inbreeding coefficient at a neutral diallelic locus under partial selfing and without condition dependence (see, e.g., pp. 92–94 in Crow and Kimura, 1970). This first-order approximation closely matches the equilibrium frequency obtained when solving exact recursions numerically (Fig. 2B; Appendix A.1.2 for details).

The equilibrium frequency given in eq. (2) reflects a balance between mutation at the numerator and selection at the denominator, reducing to the classical expression for a deleterious allele at mutation-selection equilibrium in a partially selfing population in the absence of condition dependence (*d*_*α*_ = 0; eq. 6b in Lande and Schemske, 1985). Condition dependence appears at the denominator, where it modulates the intensity of selection against allele a. Positive condition dependence (*d*_*α*_ *>* 0) decreases the equilibrium frequency of allele a relative to the condition independent case, whereas negative condition dependence (*d*_*α*_ *<* 0) increases its frequency. Strongly negative condition dependence (*d*_*α*_ *<* −2*s*, second line in eq. 2) can even bring the deleterious allele close to fixation (selection on allele a becomes positive as *s* + *d*_*α*_/2 becomes negative), where it is then maintained at a balance between selection and mutation reintroducing wild-type alleles into the population (Fig. 2B). Here, condition dependence affects selection at the condition locus because it leads alleles A and a to be associated with different average selfing rates: the allele associated with the higher rate enjoys a transmission advantage over the other, as selfing parents contribute more gene copies per offspring on average (Fisher, 1941), which causes this allele to increase in frequency.

The effect of condition dependence on the equilibrium frequency of the deleterious allele is reflected in the mean condition of the population, which increases as this allele becomes less frequent under positive condition dependence, and conversely decreases under negative condition dependence (white triangles in Fig. 2B).

#### Arbitrary relationship between condition and selfing

We next investigate how the mechanism identified in the linear case applies more generally, by computing the equilibrium frequency of allele a for arbitrary genotype-specific selfing rates using a numerical approach. Consistent with our analytical results, we find that deleterious allele a can rise to a high frequency once the selfing rate of deleterious homozygotes (*α*_aa_) is sufficiently high relative to that of wild-type homozygotes (*α*_AA_), owing to the transmission advantage that it then enjoys (Figs. 2C-E). In addition, we find that whereas polymorphism was always maintained at a balance between mutation and selection in the linear case (and so would collapse in the absence of recurrent mutation), both alleles can persist as a protected polymorphism without recurrent mutation when heterozygotes show a higher selfing rate than both homozygotes. This is because the allele that is rarer in the population is more likely to be found in heterozygous state, and so has a higher average selfing rate than the frequent allele when heterozygotes self-fertilise more than homozygotes, leading to a form of overdominance and the maintenance of polymorphism (we show this analytically in Appendix A.1.3).

### 2.3 Evolution of condition-dependent selfing

We now consider the evolution of condition-dependent selfing. To characterise selection on genotype-specific selfing rates ***α*** = (*α*_AA_, *α*_Aa_, *α*_aa_), we employ an invasion analysis approach to compute the selection gradients on each of these rates, *S*_AA_ (***α***), *S*_Aa_ (***α***), and *S*_aa_ (***α***), which we collect in vector

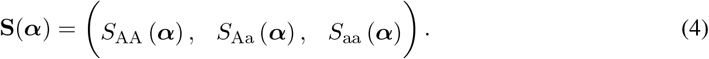

Each of these gradients captures the strength and direction of selection on the corresponding genotype-specific selfing rate in a population expressing ***α*** (Appendix A.2.1, Leimar, 2009). A positive (respectively negative) selection gradient indicates that selection favours an increase (respectively a decrease) in trait value, and the absolute value of the gradient indicates the strength of selection in this direction. Thus, condition dependence is favoured here whenever the sign of the gradient differs between genotype-specific selfing rates. We provide a detailed account of our analyses in Appendix A.2 and summarise our key results below.

Computing the selection gradients analytically for any ***α*** is difficult. We therefore start by investigating special cases where analytical expressions can be obtained, beginning with the case where the population is completely outcrossing (i.e., ***α*** = **0**). We show in Appendix A.2.2 that the selection gradients are then given by

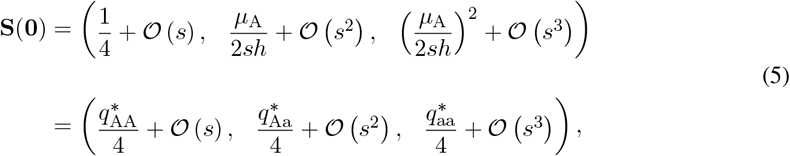

where 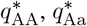 and 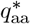 are the frequencies of genotypes AA, Aa and aa in the outcrossing population, respectively. The three gradients in eq. (5) are proportional to these frequencies because the strength of selection on a trait is proportional to the frequency with which it is expressed. Further, they are all positive, meaning that selection favours self-fertilisation irrespective of condition in a fully outcrossing population. This is because, when the rest of the population is fully outcrossing, alleles coding for increased self-fertilisation benefit from a substantial transmission advantage regardless of condition.

We next compute selection gradients under complete selfing, that is, in a population fixed for ***α*** = **1**_3_ = (1, 1, 1). We show in Appendix A.2.3 that these are given by

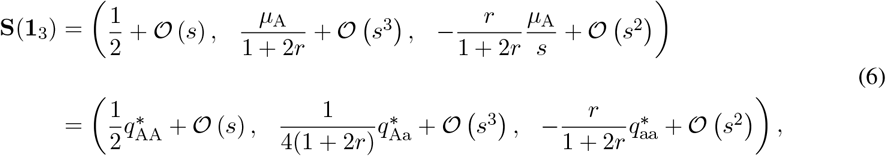

which indicates that in a completely selfing population, wild-type homozygotes and heterozygotes are selected to remain fully selfing (*S*_AA_ (**1**_3_) *>* 0 and *S*_Aa_ (**1**_3_) *>* 0), whereas deleterious homozygotes are selected to become more outcrossing (*S*_aa_ (**1**_3_) *<* 0).

This can be understood as follows: in a completely selfing population, heterozygous individuals at the condition locus are rare and homozygotes only produce offspring of the same genotype (unless mutation occurs, which is negligibly rare because *µ*_*x*_ ≪ *s*). As a result, wild-type and deleterious homozygotes effectively form distinct lineages that do not cross, but compete for breeding spots. In this setting, deleterious homozygote lineages leave no descendant in the long-term because they are always outcompeted by the wild-type, so that their class-reproductive value tends to zero (see Appendix A.2.3 for details). Now, consider the fate of a mutant allele arising at the selfing modifier in a deleterious homozygote individual. If such an allele encodes complete selfing in this genetic context, it is guaranteed to leave no long-term descendant because it is trapped in a lineage that is doomed to extinction. In contrast, if this allele encodes some outcrossing, some of the ovules produced by its bearer will be outcrossed by pollen carrying the wild-type allele at the condition locus, which will give the mutant allele an opportunity to recombine onto a wild-type haplotype. Thus, from the point-of-view of alleles at the selfing modifier, outcrossing constitutes a strategy of escape from certain doom and is favoured as such. In line with this interpretation, the strength of selection on *α*_aa_ is zero in the absence of recombination (*r* = 0) because there is then no way to escape the deleterious background through outcrossing, and increases with *r* (eq. 6).

Finally, we analysed the evolution of genotype-specific selfing rates in the general case using a numerical approach described in Appendices A.2.4 and A.2.5. This analysis reveals that while selection always favours an increase in the selfing rate of wild-type homozygotes and heterozygotes, the genetic escape mechanism described above selects for ever lower selfing rates in deleterious homozygotes once other genotypes are highly selfing, so that eventually the population converges to complete outcrossing in deleterious homozygotes and complete selfing in other genotypes, i.e.,

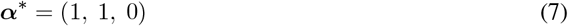

at evolutionary equilibrium, when recombination is sufficiently frequent (*r* ≫ 0; Supp. Fig. S1). Intuitively, once other genotypes are fully selfing, it is always preferable for deleterious homozygotes to outcross more, as any selfing lowers the chances of modifier alleles to escape this poor background.

When the recombination rate is very low (*r* ≈ 0), however, complete selfing evolves for all genotypes. This is because the advantage of outcrossing as a deleterious homozygote becomes weak in this case, as it provides poor chances of recombining onto a wild-type haplotype (i.e., to escape), so that the production of wild-type offspring through mutation and selfing becomes a better escape route. Accordingly, this effect vanishes and the condition-dependent strategy in eq. (7) is always favoured when the deleterious allele cannot mutate back into the wild-type (*µ*_a_ = 0; Fig. S2 in Appendix A.2.5).

### 2.4 The effect of a fixed inbreeding depression component

The analyses presented so far assume that self-fertilisation does not generate any inbreeding depression outside of that caused by alleles segregating at the condition locus. As a result, individuals carrying the wild-type allele benefit from the transmission advantage of selfing with little to no cost, so that complete selfing is especially favoured for these genetic backgrounds. Before turning to simulations to investigate a more realistic scenario where deleterious alleles at many loci may generate substantial inbreeding depression, we extend our two-locus model to investigate how a fixed inbreeding depression component would influence the evolution of condition dependence. Specifically, we assume that outcrossed ovules mature into viable seeds with probability one, whereas self-fertilised ovules now only do so with probability 1 − *δ*, where *δ* ∈ [0, 1] measures the strength of inbreeding depression.

We analysed this extended model numerically using the approach as described in Appendix A.2.5. These analyses reveal that inbreeding depression has no effect on the evolutionary equilibrium (eq. 7) when *δ <* 1/2, unless it is very close to a half (*δ* ~ 1/2), in which case we find that selection now favours heterozygotes to evolve complete outcrossing (***α***^∗^ = (1, 0, 0); Fig. S3). This is because the transmission advantage of selfing is substantially reduced in this case (owing to high selfed seed mortality), so that complete selfing no longer maximises the rate at which modifiers alleles are transmitted on wild-type background in heterozygotes (whereas it still does in wild-type homozygotes). Once inbreeding depression exceeds a half (*δ >* 1/2), complete outcrossing is favoured for all genotypes irrespective of condition, as expected under classical theory (Lande and Schemske, 1985).

## 3 Polygenic condition and the evolution of condition-dependent selfing

Our two-locus model suggests that selection broadly favours positive condition dependence for the selfing rate. We now consider the likely more realistic case where condition is the result of expression of alleles at many loci. Such a polygenic case is relevant because numerous deleterious mutations typically segregate simultaneously in natural populations and contribute to inbreeding depression, the principal force opposing selfing evolution, which is largely absent from the two-locus model (Charlesworth et al., 1990; Charlesworth and Willis, 2009; Agrawal and Whitlock, 2012; Roze, 2015). Investigating the evolution of condition-dependent selfing in this setting becomes more challenging, as individuals may carry many different multilocus genotypes at condition loci, so that we can no longer rely on a modifier locus encoding genotype-specific selfing rates. Instead, we must now consider the evolution of the plastic response of selfing to condition as a function that takes individual condition as input and returns a mating strategy. To study this, we turn to individual-based simulations.

### 3.1 The model

We simulate a population of *N* diploid cosexuals that follow the same life cycle as in the two-locus model: each individual *i* ∈ {1, …, *N*} first produces male and female gametes proportionally to its condition *φ*_*i*_ and then self-fertilises a fraction *α*_*i*_ of its ovules, while the remaining fraction is fertilised via random mating. Following mating, seeds mature, adults die and the next generation is recruited from the produced seeds.

Individual condition is affected by *L* ⩾ 1 unlinked loci. Each locus *ℓ* ∈ {1, …, *L*} has two alleles, a wild-type allele A_*ℓ*_ and a deleterious allele a_*ℓ*_ which mutate during meiosis at a rate *µ*_A_ = *µ*_a_ = *µ* (i.e., mutation to and from the deleterious allele occur at the same per-allele rate). Allele a_*ℓ*_ decreases individual condition by a proportion *s* when homozygous, and expresses proportionally to its dominance coefficient *h* when heterozygous. Condition loci are assumed to act multiplicatively (Charlesworth, 1990; Roze, 2015), so that the condition *φ*_*i*_ of an individual *i* homozygous for the deleterious allele at 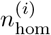 loci and heterozygous at 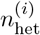 loci is given by

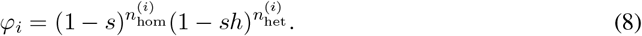

To model the evolution of condition-dependent self-fertilisation, that is of the function

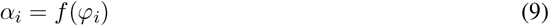

that takes the condition *φ*_*i*_ of individual *i* as input and returns its selfing strategy *α*_*i*_, one approach would be to assume that *f* (*φ*_*i*_) has a specific shape (e.g., a linear function of condition, Iwasa and Pomiankowski, 1994) and consider the evolution of its parameters (e.g., its slope and intercept in the linear case). This approach simplifies the analysis, but introduces constraints that might interfere with the evolution of condition dependence. Instead, we assume that the selfing rate expressed by an individual is determined by a gene regulatory network that may evolve to respond to condition (Fig. 3A). Gene regulatory networks have been used in other contexts to model the evolution of a plastic responses to abiotic or social cues (e.g., Ezoe and Iwasa, 1997; Deshpande and Fronhofer, 2022, 2025). They have been shown to be able to produce a wide range of functions from a limited number of genes, therefore allowing us to study the evolution of condition-dependent selfing with fewer constraints.

**Figure 3:**
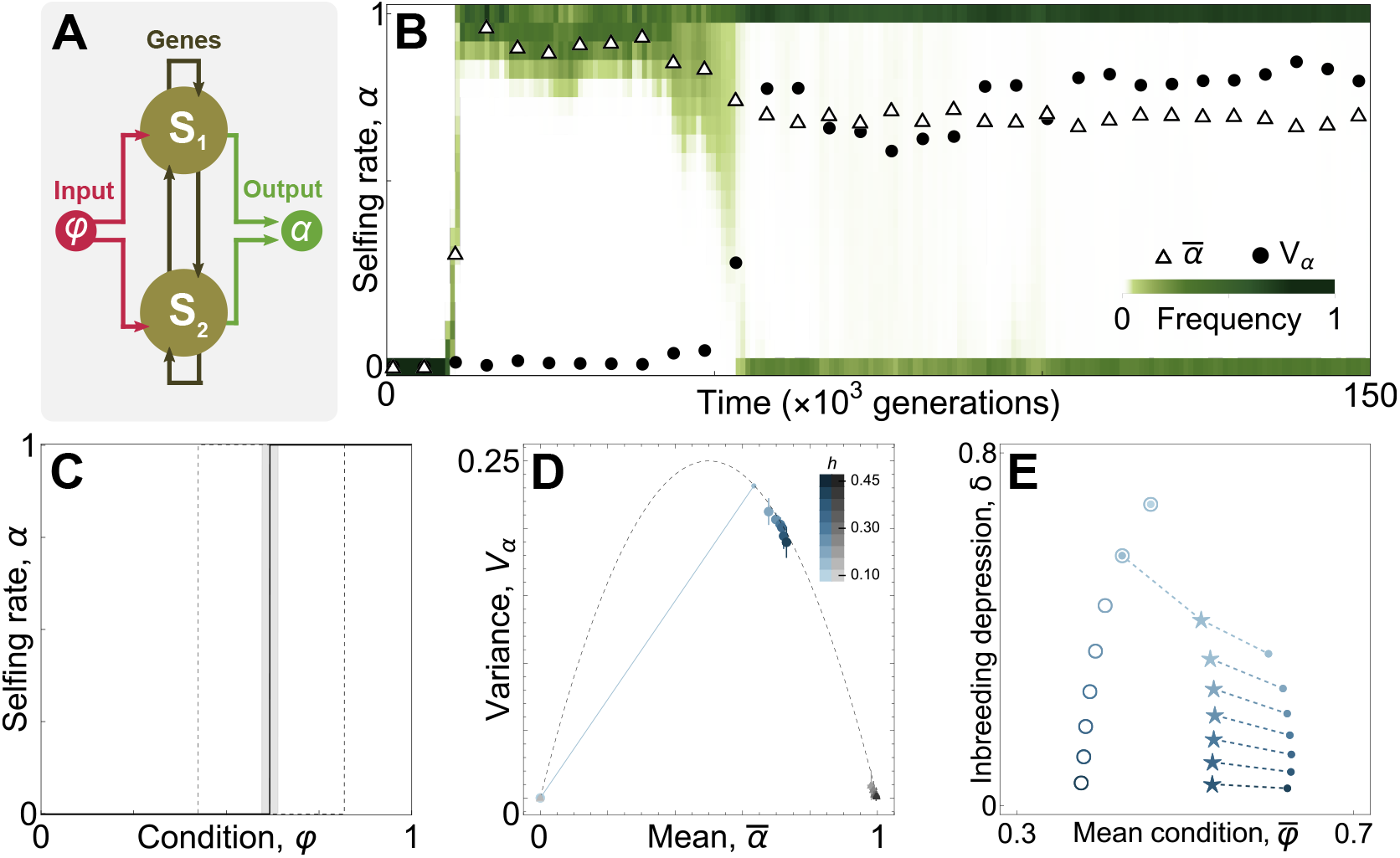
Evolution of condition-dependent selfing and its effect on the mutation load. **A** A gene regulatory network with *n*_g_ = 2 loci. Gene expression is affected by condition in proportion to input weights and by regulatory interactions among loci, which are given by the regulatory matrix. Once loci reach a steady state expression level, their equilibrium expression level is combined proportionally to output weights to obtain the individual’s mating strategy. See Appendix B.1 for details. **B** Distribution of selfing rates in the population as a function of time in a sample simulation with *L* = 10^3^ loci affecting condition. Shades of green indicate the proportion of individuals with a selfing rate in the corresponding interval at a given time step. The darker the colour, the more individuals. Triangles indicate the mean selfing rate and black points indicate variance scaled by the highest possible variance (variance of a binomial distribution with success probability 1/2, which corresponds to a case where exactly half of the population has a selfing rate *α* = 0 and the other half has *α* = 1). The population rapidly evolves complete selfing once mutation is triggered at network loci, and then experiences the gradual establishment of condition dependence. Parameters specific to this simulation: *h* = 0.25. **C** Distribution of evolved condition-dependent selfing strategies for a hundred individuals every 50 generations over the last 10^5^ generations. The solid black line indicates the median, grey shading gives the first and third quartiles and dashed lines indicate the minimal and maximal values, respectively, for each condition value. All individuals express a step-wise relationship between condition and selfing, varying only in the position of their step. **D** Mean and variance in the selfing rate averaged over the last 10^5^ generations and between replicates. Blue points indicate runs of the complete model and grey triangles indicate control runs, darker colours indicate higher dominance coefficients (see legend on top right corner). The dashed line indicates the maximum achievable variance, which is reached when the population evolves condition-dependent selfing. The solid blue line connects replicates of the complete model for *h* = 0.15, some of which evolved condition-dependent selfing (4 out of ten), while the rest remained fully outcrossing. **E** Mean condition and inbreeding depression at evolutionary equilibrium as a function of the dominance coefficient of deleterious mutations. Open circles indicate where initial populations stood under complete outcrossing. Full circles indicate simulations where condition dependence evolved, and stars show the outcome of simulations with the same mean selfing rate, but no condition dependence. Parameters used in all simulations: *N* = 5 × 10^3^, *L* = 10^3^, *s* = 0.05, *µ* = 5 × 10^−4^, *µ*_g_ = 5 × 10^−3^, *σ*_g_ = 5 × 10^−2^.

The architecture of the network is inspired from the Wagner model (Wagner, 1994) and closely resembles the one used by previous authors (Ezoe and Iwasa, 1997; Deshpande and Fronhofer, 2022, 2025). A detailed explanation of the network model can be found in Appendix B.1. Briefly, the network involves a fixed number of loci, each of which contains a protein-coding gene that is expressed during individual development from seed to adult, and contributes to determining its selfing rate at the adult stage. The level of expression of these genes throughout the development of an individual is influenced by individual condition and by loci involved in the network (including by themselves), which may up- or down-regulate gene expression through cis-regulatory interactions. The selfing rate expressed by the individual is obtained as a weighted sum of the steady-state expression levels reached by network loci at the end of development (Fig. 3A). The attributes of network loci that control their sensitivity to condition and regulatory interactions as well as their final contribution to phenotype evolve under a ‘continuum-of-alleles’ model (Kimura, 1965). Alleles at each network locus are additive, so that the attributes of a given locus in a diploid individual are given by the average of the two alleles, and network loci are assumed to recombine freely during meiosis. Further details on the model can be found in Appendix B.1.

### 3.2 Condition-dependent selfing evolves from gene-regulatory interactions and reduces the mutation load

We ran simulations for dominance coefficients ranging from *h* = 0.10 to *h* = 0.45 while keeping all other parameters fixed (see figure caption for parameter values, and Appendix B.1 for details on the simulation procedure). For each parameter set, we ran ten replicates of the model and additionally ran ten “control” runs where the network received a random value sampled from a uniform distribution as input instead of condition. These controls allowed us to check that the relationship between condition and selfing that evolves in our model is due to selection for condition dependence and not an artefact of network structure. For each simulation, the population was initialised as fixed for wild-type alleles at all condition loci and monomorphic for network loci attributes sampled in a normal distribution with mean zero and standard deviation 2*σ*_g_, except for output weights which were fixed at zero to ensure that the population was initially fully outcrossing. Mutation was switched off at network loci for the first 5 × 10^3^ generations to allow the population to reach mutation-selection balance at condition loci. Mutation was then allowed at network loci, and simulations were left to run for a total of 5 × 10^5^ generations.

When inbreeding depression was weak enough to allow the initial emergence of self-fertilisation from initially complete outcrossing, simulations of our complete model then always showed the gradual emergence of condition dependence (Fig. 3B) and the establishment of a step-wise relationship between condition and selfing, with individuals above a condition threshold being purely selfing and lower condition individuals being purely outcrossing (Fig. 3C). In contrast, control runs remained close to complete outcrossing for high levels of inbreeding depression (*δ >* 1/2, Lande and Schemske, 1985) and evolved near complete selfing otherwise, with little variance in selfing rate among individuals in both cases (grey triangles in Fig. 3D), as expected in the absence of information on individual condition. The step-wise relationship that evolves in our simulations is consistent with the notion that positive condition dependence evolves as an escape strategy from deleterious backgrounds. Indeed, from the point of view of a selfing modifier, the cost-benefit balance of selfing (staying) versus outcrossing (escaping) in a particular genetic background is independent of the selfing rate expressed by its bearer. Either the load carried by the bearer is sufficiently low for selfing to be beneficial, or the load is high enough to make outcrossing preferable.

We also quantified the effect of condition dependence on the mutation load maintained at mutation-selection balance. To do this, we simulated populations with a fixed selfing rate that we took to be the average selfing rate observed over the last 5 × 10^4^ generations in simulations where condition-dependent selfing evolved. We then compared the equilibrium mean condition and inbreeding depression attained in these fixed selfing simulations with the corresponding simulation runs where condition dependence had evolved (Fig. 3E). We found that positive condition dependence increases mean condition substantially and decreases inbreeding depression relative to the fixed selfing case. This is because positive condition dependence grants a transmission advantage to wild-type alleles, as they tend to be found in individuals in better condition who self-fertilise at a higher rate, which leads to more efficient purging.

Overall, simulation results indicate that the mechanisms described in our two-locus model also apply when condition is affected by many loci. Positive condition dependence for the selfing rate is favoured by natural selection, as it offers modifiers an escape strategy from more deleterious backgrounds, and reduces the mutation load in the population. Under the scenario considered here, selection favours an extreme form of condition dependence whereby low-condition individuals are fully outcrossing and high-condition ones are fully self-fertilising. We next extend our simulation model to investigate how two additional ecological mechanisms, namely environmental fluctuations (Section 3.3) and pollen discounting (Section 3.4), might influence the evolution of condition-dependent selfing.

### 3.3 Environmental fluctuations destabilise condition dependence

So far, variation in condition between individuals was only determined by their genotype at condition loci. However, variation in condition can also result from environmental differences between individuals (Wilson and Nussey, 2010; Forsythe et al., 2021), for example if some individuals germinate on better quality soil or in less disturbed areas than others. Environmental effects might be especially relevant to the evolution of condition-dependent selfing, because they may obscure the relationship between individual condition and genotype at condition loci, which could in turn affect the evolution of condition dependence.

To study these effects, we simulate a population that follows the same life cycle as before (Section 3.1), except that it now evolves in a heterogeneous environment composed of many patches that host one adult plant each. Seeds produced by these plants are all dispersed away from their natal patch. Each patch is characterised by an environmental variable, *ϵ*_*i*_, which changes every generation and follows a normal distribution, i.e.,

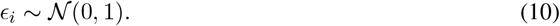

This variable could represent any relevant ecological factor, such as exposure to sunlight or water availability. A patch with *ϵ*_*i*_ = 0 is optimal for plant development, and any deviation from this optimum reduces individual condition. More precisely, the condition of individual *i*, carrying 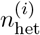 and 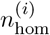 heterozygous and homozygous deleterious mutations and developing in an environment *ϵ*_*i*_ is given by

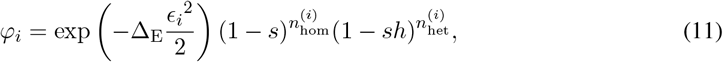

where the first term captures the effect of the environment on condition, and the second and third term give the effect of mutations in homozygous and heterozygous state on condition, as before. The intensity of environmental effects increases with parameter Δ_E_ ⩾ 0, and a stable environment is recovered when Δ_E_ = 0.

We ran simulations where we varied the intensity of environmental effects (Δ_E_ = {0.05, 0.25, 0.5, 1}), keeping all other parameters fixed for values that allowed the emergence of condition dependence in a stable environment. We find that environmental fluctuations have little impact on the evolution of condition-dependent selfing when their effect on condition is moderate (Δ_E_ = 0.05 and Δ_E_ = 0.25): a step-wise relationship between condition and selfing eventually evolves and is stable once established in most replicates, resulting in a high variance in selfing rate (Fig. 4A). Under strong environmental effects (Δ_E_ ⩾ 0.5), meanwhile, evolutionary trajectories vary widely between replicates. Some rapidly evolve a step-wise relationship, whereas others take much longer to do so, sometimes remaining at a low level of condition dependence for the entire simulation, with a low variance in selfing rate. These differences among replicates are likely due to the fact that environmental fluctuations make condition a poorer indicator of individuals’ genetic background and therefore weaken selection for condition dependence: because network loci are initialised with random attributes at the beginning of each simulation, some populations have an easier time evolving condition dependence than others from their initial state, and while this is the case in a stable environment as well (Δ_E_ = 0, see Fig. S4), differences between populations are likely exacerbated under weaker selection.

**Figure 4:**
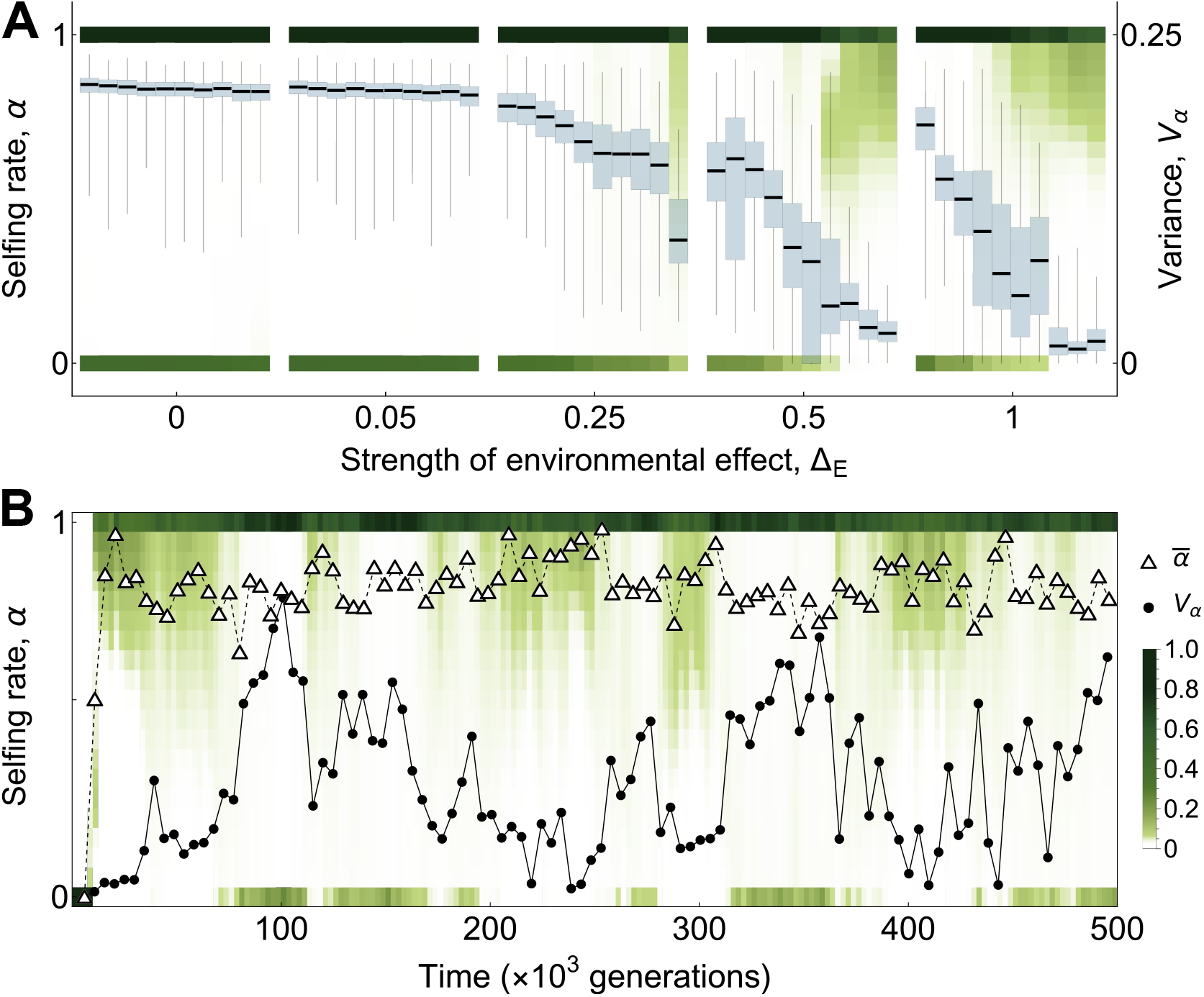
Effect of environmental fluctuations on the evolution of condition-dependent selfing. **A** Distribution of selfing rates in the population in the last 10^5^ generations of the simulation for ten replicates and different strengths of environmental effects, increasing from left to right. Shades of green indicate the proportion of individuals expressing a selfing rate in the corresponding interval. The darker the colour, the more individuals in that interval (the colour scale is the same as below). Blue box plots show the distribution of the variance in selfing rate in the population over the last 10^5^ generations. The distribution of selfing rates is bimodal with complete selfers and complete outcrossers, resulting in high variance when environmental effects are weak. Stronger environmental effects lead to more diverse distributions because they hinder the emergence of condition dependence and lead to its recurrent collapse and re-emergence once evolved. **B** Distribution of selfing rates in the population as a function of time in a sample simulation under strong environmental fluctuations (Δ_E_ = 1). Shades of green indicate the proportion of individuals with a selfing rate in the corresponding interval at a given time step. The darker the colour, the more individuals. Triangles indicate the mean selfing rate and black points indicate variance. The population initially evolves condition dependence, and then repeatedly loses and re-evolves it. Parameters used in all simulations: *N* = 5 × 10^3^, *L* = 10^3^, *s* = 0.05, *h* = 0.25, *µ* = 5 × 10^−4^, *µ*_g_ = 5 × 10^−3^, *σ*_g_ = 5 × 10^−2^.

We also find that strong environmental effects can affect the stability of condition-dependent selfing once it has evolved, with some populations repeatedly losing and later re-evolving it (Fig. 4B; see Supp. Figs. S4-S8 in Appendix B.3 for more replicates). This repeated loss of condition dependence may be due to the fact that strong environmental effects often cause individuals with a good genetic background to be in poor condition, which leads them to outcross despite selfing being their optimal strategy. In this context, an allele making its bearer fully selfing irrespective of condition could increase in frequency if it finds itself in a good genetic background, and cause the collapse of condition dependence.

### 3.4 Pollen discounting leads to a continuum of mating strategies

Traits that promote selfing often interfere with pollen export, giving rise to a trade-off between selfing and male siring success through outcrossing known as pollen discounting (Harder and Wilson, 1998), which can favour intermediate selfing rates in the absence of condition dependence (Johnston, 1998). As a final extension, we study the effect of pollen discounting on the evolution of condition-dependent selfing. We consider a population evolving in a stable, homogeneous environment (i.e., no environmental fluctuations). Individuals follow the same life cycle as before (Section 3.1), except that the amount of pollen exported by individual *i, β*_*i*_, and so its contribution to the outcrossing pollen pool, now decreases with its selfing rate. Specifically, it is given by

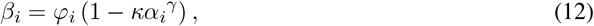

where *κ* ∈ [0, 1] measures the intensity of pollen discounting, and *γ >* 0 is a parameter controlling the shape of the trade-off between selfing and pollen export (see Appendix B.2.2 for details).

Previous work demonstrates that linear or convex trade-off curves (*γ* ⩽ 1) do not allow the maintenance of intermediate selfing rates (Johnston, 1998; Abu Awad and Roze, 2020), and so should not lead to qualitatively new results in our model. We therefore focused on concave relationships (*γ >* 1). We varied the intensity (*κ*) and concavity (*γ*) of the trade-off between selfing and pollen export, while keeping all other parameters fixed. Our simulations show that weak pollen discounting (small *κ*) leads to a step-wise relationship – similar to the baseline model – irrespective of the shape of the trade-off curve, presumably because pollen discounting only weakly affects the costs and benefits of selfing in this case. Stronger pollen discounting (large *κ*), in contrast, favours a more gradual increase in selfing rate with individual condition, leading to a continuum of selfing strategies (Fig. 5A,B). This is because the cost of selfing in terms of siring success increases sharply with the selfing rate under strong pollen discounting, causing the benefits of self-fertilisation to be overcome by the loss of outcrossing opportunities past a critical selfing rate that depends on condition. These results illustrate that including additional mechanisms relevant to the dynamics of mating can alter the relationship that evolves between condition and selfing, resulting in a diverse array of condition-dependent, mixed mating strategies.

**Figure 5:**
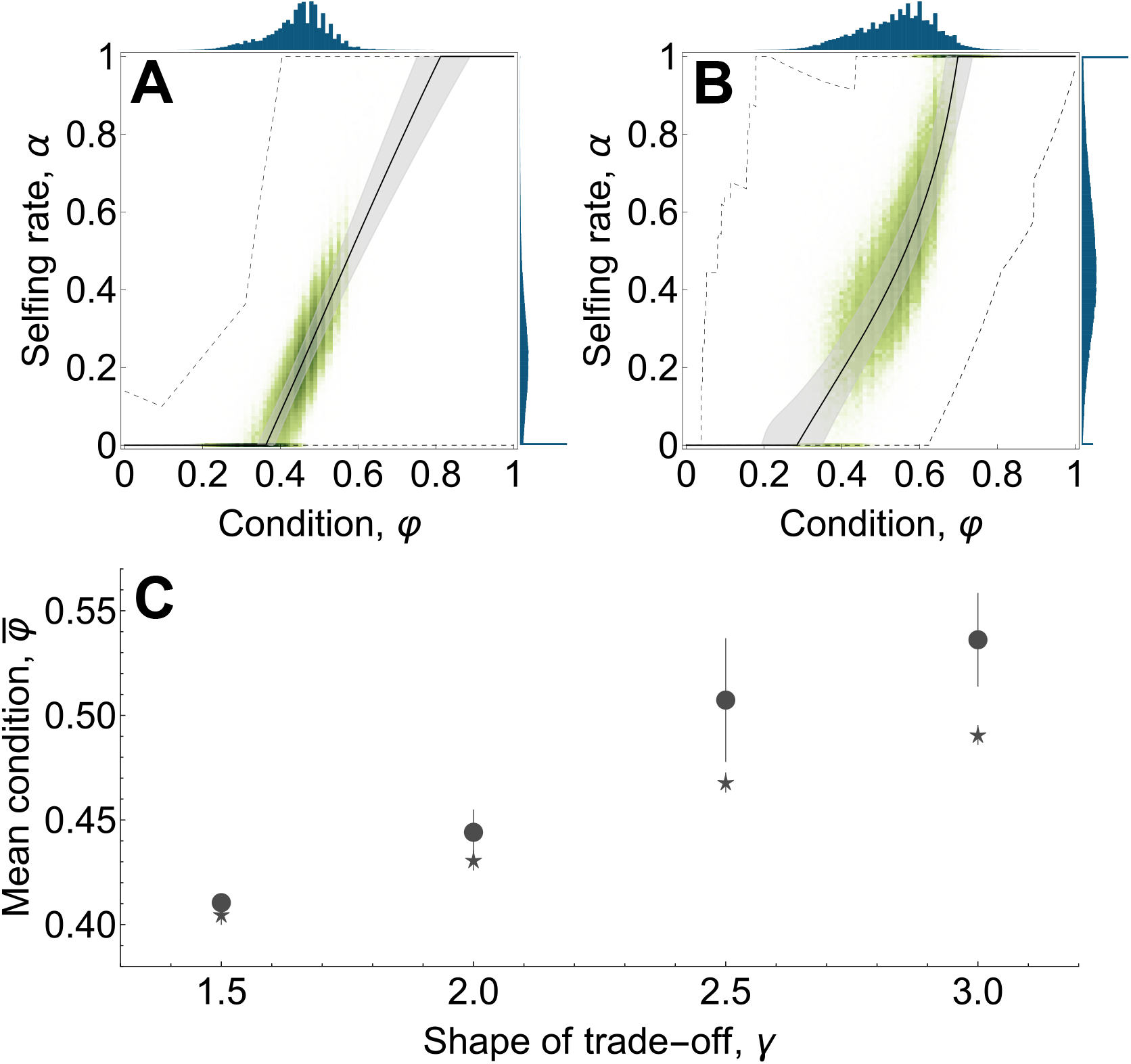
Effect of pollen discounting on the evolution of condition-dependent selfing and the mutation load. **A-B** Evolved condition-dependent selfing strategies in sample simulations under strong pollen discounting (*κ* = 1) for *γ* = 2 in panel **A** and *γ* = 3 in panel **B**. Shades of green represent the selfing rate and condition of a hundred individuals every 50 generations over the last 10^5^ of a sample simulation. The lines give the quantiles of the distribution of selfing rates encoded by the gene regulatory networks carried by these same individuals as a function of condition. The solid line gives the median and the grey shading indicates the first and third quartiles. The dashed lines give the maximum and minimum selfing rates observed in the simulations. Note that the relationship between condition and selfing is only subject to selection over the range of condition values that actually occur, so that there are intervals on both ends of the relationship that behave as effectively neutral. The blue histograms give the distribution of condition (top) and selfing rate (right) in the population over the last 10^5^ generations. **C** Effect of condition dependence on mean condition under strong pollen discounting. Points indicate mean condition over the last 10^5^ generations of simulations where condition dependence evolved, and stars indicate mean condition in a simulation with a fixed selfing rate, taken to be the average rate observed in the corresponding simulation where condition dependence evolved. Parameters used in all simulations: *N* = 5 × 10^3^, *L* = 10^3^, *s* = 0.05, *h* = 0.25, *µ* = 5 × 10^−4^, *µ*_g_ = 5 × 10^−3^, *σ*_g_ = 5 × 10^−2^.

We also quantified the effect of condition dependence on the load. We compared mean condition in the population with mean condition in a population where all individuals have the same fixed selfing rate, which we took to be the mean selfing rate observed in simulations where condition dependence evolved. We find that condition dependence again reduces the load relative to the condition independent case, though the effect is less pronounced for more linear trade-off curves (Fig. 5C). This is because more linear curves cause pollen discounting to increase more rapidly with selfing, which leads to lower selfing rates in highest-condition individuals. As a result, the difference in selfing rate between individuals with different genetic backgrounds is reduced, which decreases the transmission advantage enjoyed by wild-type alleles and so makes purging owing to condition dependence less efficient.

## 4 Discussion

Most theory on the evolution of self-fertilisation considers genetically determined, non-plastic mating strategies (Goodwillie et al., 2005; Clo et al., 2025). In this paper, we investigated how condition-dependent selfing might evolve, whereby individuals plastically adjust their mating strategy according to their biological condition. We found that positive condition dependence, where higher condition individuals self-fertilise at a higher rate, evolves whenever the mutation load is weak enough to allow the initial emergence of self-fertilisation from complete outcrossing. This positive relationship between selfing and condition is favoured as an escape strategy from deleterious genetic backgrounds, similar to the “abandon ship” mechanism, which has been shown to select for individuals in poorer condition to engage more in sexual reproduction (Hadany and Otto, 2007; Otto, 2009), recombine more often (Agrawal et al., 2005), and undergo mutation at a higher rate during meiosis (Ram et al., 2018). The evolution of condition-dependent self-fertilisation may therefore represent a novel manifestation of a more general mechanism, with specific biological implications that we discuss below.

We showed that selection favours a positive relationship between condition and the selfing rate, which we modelled as a single trait. In reality, however, the rate at which a plant self-fertilises is likely determined by the interaction of many aspects of its phenotype including, for example, the distance between anthers and pistils (Opedal, 2018), the strength of self-incompatibility reactions in the style (Levin, 1996), the plant’s allocation to pollen production (Damgaard and Abbott, 1995; Chen and Pannell, 2024, 2025) or the overlap in the timing of sexual maturity of its female and male function (Kalisz et al., 2012; Barrett and Harder, 2017a). Our results imply that all these traits, rather than the selfing rate directly, should be under selection to evolve condition-dependent expression through their effect on selfing. The degree of condition dependence favoured for a given trait may however depend on several factors. A key factor that we highlighted is how much pollen discounting is caused by the trait when promoting selfing, as we showed that the shape and intensity of the relationship between selfing and pollen export is critical to the condition-dependent strategy favoured by selection. Traits that affect selfing are also likely to be under selection from other ecological and physiological sources, such as pollinator preferences or functional constraints (e.g., on floral morphology, Barrett, 2002), as well as life-history trade-offs, which could interact with selection for condition dependence in ways that have not been explored. In the future, understanding how this interaction between sources of selection shapes the evolution of traits affecting the selfing rate may provide useful insight into the diversity and phenotypic underpinnings of mating strategies in cosexual species, and in particular into the mechanisms promoting the evolution of mixed mating, which remain elusive (Goodwillie et al., 2005; Clo et al., 2025).

Among the many traits that affect selfing, plant size has received special attention. At an intraspecific level, larger plants can show elevated selfing rates, a pattern usually viewed as a consequence of increased pollen transfers among flowers of the same individual as the number of simultaneously displayed flowers increases (geitonogamy; de Jong et al., 1993; Karron et al., 2004; Williams, 2007; Christopher et al., 2021), and interpreted as a cost to these plants in terms of lost outcrossing opportunities (de Jong et al., 1993; Harder and Barrett, 1995; Barrett, 2003). Our results suggest that this increased selfing in larger plants may also be favoured by selection through two pathways. First, the size of a plant depends on its ability to extract resources from the environment and allocate them to growth, and may therefore often depend on its condition. If higher-condition plants tend to be larger, then condition-dependent selfing would cause larger individuals to self-fertilise more frequently even if size *per se* did not affect selfing. Second, since plant size is indeed thought to have an effect on its selfing rate, allocation to growth could be selected to become condition-dependent owing to this effect. Through this effect on growth, selection for condition-dependent selfing could therefore influence the evolution of traits that do not directly relate to mating as plant size plays a central role in many aspects of a plant’s ecology, including its ability to competitively acquire resources or its attractiveness to animal pollinators, dispersers and predators.

There is empirical evidence that environmental stress can affect individual selfing rates through plastic changes in mating-related traits (Levin, 2010; Suijkerbuijk et al., 2025). For example, Holtsford and Ellstrand (1992) found that herkogamy and dichogamy are substantially reduced in *Clarkia tembloriensis* individuals grown in hot and dry summer conditions relative to more clement spring weather, resulting in higher selfing rates under stress. This pattern of increased selfing in more challenging conditions was observed in other species as well (e.g., Motten and Stone, 2000; Camargo et al., 2017; see Levin (2010) for a review). Environmental stress has also been associated with changes in the ratio of closed (obligately selfing) to open (potentially outcrossing) flowers in cleistogamous species, with higher proportions of closed flowers produced under stress in a majority of cases (e.g., Cheplick, 2007, see Oakley et al., 2007 and Culley and Klooster, 2007 for reviews). Although limited to a few species, this empirical evidence suggests that plants in poor condition owing to environmental stress might often exhibit higher rather than lower selfing rates. Such negative condition dependence for the selfing rate could be explained by pollen limitation, which we did not consider here. Under inbreeding depression high enough for outcrossing to be favoured in all individuals when pollen is not limiting, theory shows that non-plastic mixed mating can be maintained in uncertain pollination environments, as it then provides reproductive assurance (Morgan and Wilson, 2005). If low condition individuals tend to be more pollen-limited, the benefits of selfing in terms of reproductive assurance would then become condition-dependent, and could therefore favour negative condition dependence for the selfing rate.

Irrespective of the particular trait, the evolution of condition dependence relies on the ability of organisms to acquire information about their genetic background. Here, we assumed that organisms are able to sense their condition directly by letting the selfing rate be determined by a gene regulatory network receiving individual condition as input. When individual condition is purely genetically determined, it acts as a reliable indicator of individual genotype and so condition dependence evolves readily from gene-regulatory interactions. Environmental effects on condition hinder the evolution of condition dependence, because they obscure the relationship between condition and genotype, which makes it a less reliable indicator and so weakens selection for condition dependence. However, no amount of environmental fluctuations could entirely prevent condition-dependent selfing from evolving in our simulations, which suggests that the evolution of condition dependence may occur as long as individual condition has some genetic basis and organisms are able to sense their condition. In addition to environmental effects, the evolution of condition dependence could also be hindered if organisms do not sense their condition directly and instead rely on phenotypic cues that correlate imperfectly with condition and might be costly to detect (DeWitt et al., 1998; Van Kleunen and Fischer, 2005). Reliance on such cues could also set up an evolutionary feedback between the mating system and trait(s) used as cue, which could for example include individual growth rate or any other measure of metabolic activity.

Like any other form of phenotypic plasticity, the evolution of condition-dependent selfing might also be limited by constraints on the shape of the plastic response that organisms are able to produce in traits affecting the selfing rate, even when individuals can readily acquire information about their genetic background (DeWitt et al., 1998; Van Kleunen and Fischer, 2005). Such limits might stem from a lack of genetic variation required for the plastic response to reach its optimal shape (Gomulkiewicz and Kirkpatrick, 1992; Lande, 2009). Genetic constraints of this kind are hardwired into models that impose the shape of the plastic response. For example, studies that consider a linear response curve (e.g., Iwasa and Pomiankowski, 1994; Lande, 2009; Flintham et al., 2023) assume that the parameters of the response (its slope and intercept) can evolve but its shape cannot, even though a linear response may not be what would be favoured by selection. Gene regulatory networks are also subject to genetic constraints of this type, as a single mutation in the network often results in only slight changes to the general shape of the resulting plastic response. However, these constraints mainly operate on the short term, as they can typically be resolved in the long-term through recurrent mutation and changes to the genetic architecture of traits in response to selection.

Constraints on the shape of plastic responses might also stem from physiological costs and limits associated with plastic trait expression (DeWitt et al., 1998; Van Kleunen and Fischer, 2005). For example, the optimal plastic response in a given trait might only be achievable through correlated changes in other traits that displace them from their optima, leading to a fitness cost. Such physiological constraints are absent from our gene regulatory network model, which lets individuals with slight differences in condition express radically different mating strategies. In the case of self-fertilisation, however, large differences in individual selfing rates are readily produced through small changes to various traits pertaining to plant phenology (e.g., the extent to which maturity through male and female function overlap), floral morphology (e.g., herkogamy or floral pollen content, Opedal, 2018; Chen and Pannell, 2024) or mate choice mechanisms (e.g., incomplete self-incompatibility reactions or selective seed abortion; Levin, 2010; Brandvain et al., 2024). Plasticity in any combinations of those traits could contribute to producing the condition-dependent strategy favoured by selection, so that physiological constraints on one trait could be circumvented through changes in others.

Our findings point to a need for empirical studies relating plant traits to condition and mating strategy at the individual level. Such data is currently lacking, in part because selfing rates are often measured at the level of the population and so can seldom be related to individual phenotype (Whitehead et al., 2018). One exception to this comes from studies of herkogamy (the spatial separation of female and male organs), which often show that closer proximity between the two sexual functions leads to higher individual selfing rates (reviewed in Opedal, 2018). One possible way of looking for condition-dependent selfing empirically could therefore be to relate the degree of herkogamy to condition at the individual level in species where it has been shown to affect the selfing rate. According to our model, lower herkogamy and accordingly higher selfing rates would be expected in higher condition individuals. Condition dependence for the selfing rate could also be revealed by an experimental design where plants with diverse genetic backgrounds are grown under controlled conditions in order to minimise environmental differences between individuals. One would quantify their relative condition through phenotypic proxies (e.g., total number of flowers, or plant biomass, which has been shown to positively correlate with fecundity in many studies; see Younginger et al., 2017 for a review) and leave them to exchange pollen. Individual selfing rates could then be estimated using the resulting seeds’ genotypes, and related to proxies of condition. A positive relationship between these proxies and the selfing rate would be expected. One could also indirectly quantify the amount of deleterious variation carried by a plant by measuring the performance of self-fertilised and outcrossed seeds produced via controlled crosses on said plant (or one with the same genotype), in order to quantify inbreeding depression within its progeny. In this setting, one would expect a negative relationship between the selfing rate of the parent and the magnitude of inbreeding depression measured among its offspring. Nevertheless, the ability of the empirical tests proposed above to detect condition dependence might be limited by the confounding effect of purging. This is because purging causes modifiers encoding higher selfing rates to more often occur on fitter genetic backgrounds (Uyenoyama and Waller, 1991; Abu Awad and Roze, 2020; Xu, 2024), which may lead to a positive relationship between condition and selfing rate even in the absence of a plastic response (this effect is however expected to be weak unless large effect modifiers and/or strongly deleterious mutations are considered, Abu Awad and Roze, 2020; Xu, 2024). An alternative approach that would not be affected by purging could be to compare the selfing rates of selfed and outcrossed offspring derived from the same maternal plant, or to grow genetically identical plants in environments of varying qualities, and examine how their selfing rates vary in response to environmentally induced variation in their condition.

The genetic escape mechanism that we highlight should also select for condition dependence in traits that affect inbreeding in general – of which selfing is but an extreme case. Beyond cosexual organisms, our results might therefore have implications for the evolution of condition dependence in organisms with separate sexes as well. In dioecious animals, this mechanism could for instance favour the evolution of condition-dependent dispersal – whereby lower condition individuals would tend to be more dispersive – or condition-dependent preference for mating with kin, in which case high condition individuals would be more prone to mate with individuals related to them. Behavioural preference and avoidance of inbreeding are both well-documented in animals (Pike et al., 2021), but the possibility that these preferences might vary between individuals as a function of their condition has not been considered, be it in theory or empirically.

In conclusion, our analyses demonstrate that selection broadly favours condition-dependent inbreeding, leading to a diverse array of mating strategies coexisting in populations. The consequences of this previously undescribed source of selection may be seen on many traits in partially selfing cosexuals as well as in dioecious organisms, though the emergence of condition dependence may be constrained by morphological and physiological trade-offs, and the strength of selection for condition dependence would be diminished by strong environmental effects or reliance on imperfect phenotypic cues. The question of how often condition-dependent inbreeding actually occurs in natural populations, and if so, through which phenotypic basis, is open for empirical investigation.

## Acknowledgements

The authors would like to thank the Swiss National Science Foundation for funding (SNF grants 310030_215135 to JRP and PCEFP3181243 to CM).

## Data availability

The code used for individual-based simulations, numerical analyses and to check analytical results is available at `10.5281/zenodo.20510607`.

## Authors contributions

TL, JRP and CM conceived the study. TL carried out the analytical work and simulations, with input from CM. TL wrote the first draft. All authors contributed to revisions and approved the final manuscript. CM and JRP supervised the work and acquired the funding.

