## Appendix for "The evolution of condition-dependent self-fertilisation"

**B Multilocus simulations** **40**

**B.1 The baseline program** **40**

**B.2 Extensions** **44**

B.2.1 Environmental fluctuations . . . . . . . . . . . . . . . . . . . . . . . . . . . . . . . . . . . 45

B.2.2 Pollen discounting . . . . . . . . . . . . . . . . . . . . . . . . . . . . . . . . . . . . . . . . . 45

**B.3 Supplementary figures for environmental fluctuations** **46**

#### Appendix A

### Analysis of the two-locus model

In this Appendix, we analyse the two-locus model described in Section 2 of the main text. We first study the effect of a fixed relationship between condition and selfing on the condition locus (Section A.1), and then consider the evolution of condition-dependent selfing (Section A.2).

#### A.1 Effect of condition-dependent selfing on the condition locus

##### A.1.1 General population genetic recursions

We consider a population fixed for an arbitrary selfing strategy $\boldsymbol{\alpha} = (\alpha_{\mathrm{AA}}, \alpha_{\mathrm{Aa}}, \alpha_{\mathrm{aa}})$. In this population, we denote the frequency of genotype $x_1x_2$ (with $x_1,\ x_2 \in \{\mathrm{A}, \mathrm{a}\}$) at some arbitrary time $t$ as $q_{x_1x_2}(t)$, and by $M_x(\boldsymbol{\alpha}, t)$ the frequency of allele $x \in \{\mathrm{A}, \mathrm{a}\}$ in the outcross pollen pool. Accounting for mutation among the gametes produced by each genotype, frequencies in the outcross pollen pool are given by

$$\begin{aligned} M_{\mathrm{A}}(\boldsymbol{\alpha}, t) &= (1-\mu_{\mathrm{A}})\, q_{\mathrm{AA}}(t)\frac{\varphi_{\mathrm{AA}}}{\overline{\varphi}(t)} + q_{\mathrm{Aa}}(t)\frac{1-\mu_{\mathrm{A}}+\mu_{\mathrm{a}}}{2}\frac{\varphi_{\mathrm{Aa}}}{\overline{\varphi}(t)} + \mu_{\mathrm{a}}\, q_{\mathrm{aa}}(t)\frac{\varphi_{\mathrm{aa}}}{\overline{\varphi}(t)}, \\ M_{\mathrm{a}}(\boldsymbol{\alpha}, t) &= \mu_{\mathrm{A}}\, q_{\mathrm{AA}}(t)\frac{\varphi_{\mathrm{AA}}}{\overline{\varphi}(t)} + q_{\mathrm{Aa}}(t)\frac{1-\mu_{\mathrm{a}}+\mu_{\mathrm{A}}}{2}\frac{\varphi_{\mathrm{Aa}}}{\overline{\varphi}(t)} + (1-\mu_{\mathrm{a}})\, q_{\mathrm{aa}}(t)\frac{\varphi_{\mathrm{aa}}}{\overline{\varphi}(t)}, \end{aligned} \tag{A1}$$

for allele $\mathrm{A}$ and $\mathrm{a}$, respectively, where

$$\overline{\varphi}(t) = \varphi_{\mathrm{AA}}q_{\mathrm{AA}}(t) + \varphi_{\mathrm{Aa}}q_{\mathrm{Aa}}(t) + \varphi_{\mathrm{aa}}q_{\mathrm{aa}}(t) \tag{A2}$$

is the mean condition in the population at time $t$. Using these notations, genotypic frequencies in the next generation are given by

$$\begin{aligned}q_{\mathrm{AA}}(t+1) = q_{\mathrm{AA}}(t)\frac{\varphi_{\mathrm{AA}}}{\overline{\varphi}(t)}(1-\mu_{\mathrm{A}})\left[\alpha_{\mathrm{AA}}(1-\mu_{\mathrm{A}}) + (1-\alpha_{\mathrm{AA}})M_{\mathrm{A}}(\boldsymbol{\alpha},t)\right] \\ + q_{\mathrm{Aa}}(t)\frac{\varphi_{\mathrm{Aa}}}{\overline{\varphi}(t)}\frac{1-\mu_{\mathrm{A}}+\mu_{\mathrm{a}}}{2}\left[\alpha_{\mathrm{Aa}}\frac{1-\mu_{\mathrm{A}}+\mu_{\mathrm{a}}}{2} + (1-\alpha_{\mathrm{Aa}})M_{\mathrm{A}}(\boldsymbol{\alpha},t)\right] \\ + q_{\mathrm{aa}}(t)\mu_{\mathrm{a}}\frac{\varphi_{\mathrm{aa}}}{\overline{\varphi}(t)}\left[\alpha_{\mathrm{aa}}\mu_{\mathrm{a}} + (1-\alpha_{\mathrm{aa}})M_{\mathrm{A}}(\boldsymbol{\alpha},t)\right],\end{aligned} \tag{A3a}$$

$$\begin{aligned}q_{\mathrm{Aa}}(t+1) = q_{\mathrm{AA}}(t)\frac{\varphi_{\mathrm{AA}}}{\overline{\varphi}(t)}\left\{2\mu_{\mathrm{A}}(1-\mu_{\mathrm{A}})\alpha_{\mathrm{AA}} + (1-\alpha_{\mathrm{AA}})\big[(1-\mu_{\mathrm{A}})M_{\mathrm{a}}(\boldsymbol{\alpha},t) + \mu_{\mathrm{A}}M_{\mathrm{A}}(\boldsymbol{\alpha},t)\big]\right\} \\ + 2q_{\mathrm{Aa}}(t)\frac{\varphi_{\mathrm{Aa}}}{\overline{\varphi}(t)}\frac{1-\mu_{\mathrm{A}}+\mu_{\mathrm{a}}}{2}\frac{1-\mu_{\mathrm{a}}+\mu_{\mathrm{A}}}{2} \\ + q_{\mathrm{aa}}(t)\frac{\varphi_{\mathrm{aa}}}{\overline{\varphi}(t)}\left\{2\mu_{\mathrm{a}}(1-\mu_{\mathrm{a}})\alpha_{\mathrm{aa}} + (1-\alpha_{\mathrm{aa}})\big[(1-\mu_{\mathrm{a}})M_{\mathrm{A}}(\boldsymbol{\alpha},t) + \mu_{\mathrm{a}}M_{\mathrm{a}}(\boldsymbol{\alpha},t)\big]\right\},\end{aligned} \tag{A3b}$$

and

$$\begin{aligned}q_{\mathrm{aa}}(t+1) = q_{\mathrm{AA}}(t)\frac{\varphi_{\mathrm{AA}}}{\overline{\varphi}(t)}\mu_{\mathrm{A}}\left[\alpha_{\mathrm{AA}}\mu_{\mathrm{A}} + (1-\alpha_{\mathrm{AA}})M_{\mathrm{a}}(\boldsymbol{\alpha},t)\right] \\ + q_{\mathrm{Aa}}(t)\frac{\varphi_{\mathrm{Aa}}}{\overline{\varphi}(t)}\frac{1-\mu_{\mathrm{a}}+\mu_{\mathrm{A}}}{2}\left[\alpha_{\mathrm{Aa}}\frac{1-\mu_{\mathrm{a}}+\mu_{\mathrm{A}}}{2} + (1-\alpha_{\mathrm{Aa}})M_{\mathrm{a}}(\boldsymbol{\alpha},t)\right] \\ + q_{\mathrm{aa}}(t)\frac{\varphi_{\mathrm{aa}}}{\overline{\varphi}(t)}(1-\mu_{\mathrm{a}})\left[\alpha_{\mathrm{aa}}(1-\mu_{\mathrm{a}}) + (1-\alpha_{\mathrm{aa}})M_{\mathrm{a}}(\boldsymbol{\alpha},t)\right].\end{aligned} \tag{A3c}$$

In each of the three equations in (A3), the three lines give the frequency of successful offspring of the considered genotype produced by AA, Aa and aa mothers, respectively.

To understand how these were obtained, let us focus on eq. (A3c), which gives the frequency of wild-type homozygotes aa in the next time step. Wild-type homozygote mothers AA contribute a proportion $q_{\mathrm{AA}}(t)\times\varphi_{\mathrm{AA}}/\overline{\varphi}(t)$ of the ovules produced by the population, a fraction $\mu_{\mathrm{A}}$ of which carry the deleterious allele a. A fraction $\alpha_{\mathrm{AA}}$ of these ovules is then self-fertilised and give rise to aa seeds if they get fertilised by a-carrying pollen, which occurs with probability $\mu_{\mathrm{A}}$. The remaining fraction $1-\alpha_{\mathrm{AA}}$ is fertilised through outcrossing, in which case they are fertilised by a-carrying pollen with probability $M_{\mathrm{a}}(\boldsymbol{\alpha},t)$. Heterozygous mothers Aa contribute a proportion $q_{\mathrm{Aa}}(t)\times\varphi_{\mathrm{Aa}}/\overline{\varphi}(t)$ of ovules, a fraction $(1-\mu_{\mathrm{a}}+\mu_{\mathrm{A}})/2$ of which carry the deleterious allele a. These are self-fertilised in proportion $\alpha_{\mathrm{Aa}}$, in which case they receive a-carrying pollen and give rise to a aa seed with probability $(1-\mu_{\rm a}+\mu_{\rm A})/2$. Otherwise, they are fertilised by outcrossing, and receive a-carrying pollen with probability $M_{\rm a}(\boldsymbol{\alpha},t)$. Finally, deleterious homozygote mothers aa contribute a proportion $q_{\rm aa}(t)\times\varphi_{\rm aa}/\overline{\varphi}(t)$ of ovules, a fraction $1-\mu_{\rm a}$ of which carry the deleterious allele a. These get self-fertilised at rate $\alpha_{\rm aa}$ and receive a-carrying pollen with probability $1-\mu_{\rm a}$, otherwise, they get cross-fertilised and receive a-carrying pollen with probability $M_{\rm a}(\boldsymbol{\alpha},t)$.

The equilibrium genotypic frequencies can be obtained from eq. (A3) by finding

$$q^*_{\rm AA},\ q^*_{\rm Aa} \text{ and } q^*_{\rm aa} \quad \text{such that} \quad q_{x_1x_2}(t)=q_{x_1x_2}(t+1)=q^*_{x_1x_2} \text{ for all } x_1,\ x_2\in\{\rm A,a\}. \tag{A4}$$

**A more practical description.** The description of population genetic dynamics in terms of genotypic frequencies given above is the most natural to derive from a particular biological scenario, but is difficult to work with in practice, especially when incorporating assumptions on the relative magnitude of parameters, as we will come to below. A more practical description, which we will use from now on, can be obtained from expressing genotypic frequencies as

$$\begin{aligned} q_{\rm AA}(t) &= (1-p(t))^2 + C(t) \\ q_{\rm Aa}(t) &= 2\left[p(t)(1-p(t)) - C(t)\right] \\ q_{\rm aa}(t) &= p(t)^2 + C(t), \end{aligned} \tag{A5}$$

where

$$p(t) = q_{\rm aa}(t) + \frac{q_{\rm Aa}(t)}{2} \quad \text{and} \quad C(t) = q_{\rm aa}(t) - p(t)^2$$

denote the frequency of allele a and the excess in homozygotes relative to the Hardy-Weinberg expectation at the condition locus at time $t$, respectively, and

$$\Delta p = p(t+1) - p(t) \quad \text{and} \quad \Delta C = C(t+1) - C(t) \tag{A6}$$

denote the change in these quantities over one generation, which is be readily obtained by rearranging eq. (A3). With this description, an equilibrium is defined as

$$p^*, C^* \quad \text{such that} \quad \Delta p = \Delta C = 0. \tag{A7}$$

**Weak variation in condition.** Throughout our analyses, we will assume that the effect of the deleterious allele $\mathrm{a}$ on condition is weak, $s \sim \mathcal{O}(\epsilon)$, and mutation at the condition locus is rare in both directions, $\mu_x \sim \mathcal{O}(\epsilon^2)$, so that $\mu_x \ll s$. To incorporate this assumption, our approach will rely in Taylor-expanding the change in allelic frequency and excess homozygosity at the condition locus around $\epsilon = 0$. In general, this will take the form

$$\mathcal{K} = \mathcal{K}_{(0)} + \epsilon\, \mathcal{K}_{(1)} + \frac{\epsilon^2}{2}\mathcal{K}_{(1)} + ... + \frac{\epsilon^n}{n!}\mathcal{K}_{(n)} + \mathcal{O}\left(\epsilon^{n+1}\right), \tag{A8a}$$

where

$$\mathcal{K}_{(n)} = \left.\frac{\partial^n \mathcal{K}}{\partial \epsilon^n}\right|_{\epsilon=0} \tag{A8b}$$

is the $n^{\text{th}}$ order perturbation of $\mathcal{K}$ with respect to $\epsilon$, and $\mathcal{K}$ can be any differentiable quantity (e.g., $\Delta p$ and $\Delta C$ here).

##### A.1.2 Linear relationship

The explicit solution to eq. (A7) cannot be obtained for an arbitrary selfing strategy $\boldsymbol{\alpha}$, as differences in selfing rates between genotypes greatly complicate the dynamics. To gain analytical traction, we assume that genotype-dependent selfing rates $\boldsymbol{\alpha} = (\alpha_{\mathrm{AA}}, \alpha_{\mathrm{Aa}}, \alpha_{\mathrm{aa}})$ follow a linear relationship. Specifically, we assume that they are given by

$$\boldsymbol{\alpha} = (\alpha_{\mathrm{AA}}, \alpha_{\mathrm{Aa}}, \alpha_{\mathrm{aa}}) = \Big(\alpha_0, \alpha_0 - h d_\alpha,\ \alpha_0 - d_\alpha\Big), \tag{A9}$$

where $d_\alpha$ determines the slope of the line on which $\alpha_{\mathrm{AA}}$, $\alpha_{\mathrm{Aa}}$ and $\alpha_{\mathrm{aa}}$ lie. The selfing rate increases with condition when $d_\alpha > 0$ (positive condition dependence) and decreases with condition occurs when

$d_\alpha < 0$ (negative condition dependence). We assume that condition dependence is weak, $d_\alpha \sim \mathcal{O}(\epsilon)$, in order to obtain analytical results, but numerical analyses show that the obtained approximation predicts exact solutions under a broad parameter range (see Fig. 2B in the main text).

Our aim is to compute the equilibrium allelic frequency $p^*$ and excess homozygosity $C^*$ at the condition locus to first order in $\epsilon$. To do so, we expand the allelic frequency $p(t)$ and excess in homozygotes $C(t)$ using eq. (A8) as

$$p(t) = p_{(0)} + \epsilon\, p_{(1)} + \frac{\epsilon^2}{2} p_{(2)} + ... + \frac{\epsilon^n}{n!} p_{(n)} + \mathcal{O}\left(\epsilon^{n+1}\right) \tag{A10a}$$

and

$$C(t) = C_{(0)} + \epsilon\, C_{(1)} + \frac{\epsilon^2}{2} C_{(2)} + ... + \frac{\epsilon^n}{n!} C_{(n)} + \mathcal{O}\left(\epsilon^{n+1}\right), \tag{A10b}$$

where we dropped time $t$ from perturbation terms for brevity. Similarly, the change in these variables over one generation can be written as sums of perturbations to the $n^{\text{th}}$ order in $\epsilon$, i.e.,

$$\Delta p = \Delta p_{(0)} + \epsilon\, \Delta p_{(1)} + \frac{\epsilon^2}{2} \Delta p_{(2)} + ... + \frac{\epsilon^n}{n!} \Delta p_{(n)} + \mathcal{O}\left(\epsilon^{n+1}\right) \tag{A10c}$$

and

$$\Delta C = \Delta C_{(0)} + \epsilon\, \Delta C_{(1)} + \frac{\epsilon^2}{2} \Delta C_{(2)} + ... + \frac{\epsilon^n}{n!} \Delta C_{(n)} + \mathcal{O}\left(\epsilon^{n+1}\right). \tag{A10d}$$

It follows from eq. (A7) that for a population to be at an equilibrium, we must have

$$\Delta p_{(n)} = \Delta C_{(n)} = 0 \tag{A11}$$

to all orders $n \in \{0, 1, ..., n\}$ in $\epsilon$. Below, we solve eq. (A11) to increasingly high orders to compute perturbations terms in eq. (A10).

Injecting eq. (A10) into eq. (A6) and expanding it to order zero in $\epsilon$, we find that the change in frequency and excess homozygosity at the condition locus are

$$\Delta p_{(0)} = 0 \quad \text{and} \quad \Delta C_{(0)} = \frac{1}{2}\Big[\alpha_0 p_{(0)}(1 - p_{(0)}) - C_{(0)}(2 - \alpha_0)\Big] \tag{A12}$$

to order zero. The change in allelic frequency is zero because all the processes that affect it, namely mutation, fecundity selection and condition dependence are neglected to this order, and so allelic frequencies will not change from one generation to the next in the absence of drift. The change in mean homozygosity to order zero, meanwhile, yields

$$\Delta C_{(0)} = 0 \quad \Rightarrow \quad C^*_{(0)} = F p_{(0)}(1 - p_{(0)}) \tag{A13}$$

at equilibrium, with

$$F = \frac{\alpha_0}{2 - \alpha_0}. \tag{A14}$$

Eq. (A13) corresponds to the expected excess homozygosity at a diallelic neutral locus under partial selfing in the absence of condition dependence (which is neglected to order zero).

The first-order perturbation of the change in frequency at the condition locus is given by

$$\Delta p_{(1)} = -\left(s + \frac{d_\alpha}{2}\right)\left[F + (1 - F)\Big(p_{(0)} + h\left(1 - 2p_{(0)}\right)\Big)\right] p_{(0)}\left(1 - p_{(0)}\right). \tag{A15}$$

The term in square brackets is strictly positive for $0 \leqslant h \leqslant 1$, so that there is two possible equilibria that satisfy eq. (A11), either $p^*_{(0)} = 0$ or $p^*_{(0)} = 1$. Which of the two is attained is determined by the relative strength of selection against the deleterious allele and condition dependence for the selfing rate. Under positive condition dependence ($d_\alpha > 0$), eq. (A15) is always negative, leading to $p^*_{(0)} = 0$. This indicates that the deleterious allele always remains rare in this situation, as higher order terms will only produce small deviations close to $p^*_{(0)} = 0$. Under negative condition dependence ($d_\alpha < 0$), on the other hand, eq. (A15) can become positive and lead to $p^*_{(0)} = 1$, indicating that the deleterious allele attains a frequency close to one at equilibrium. Specifically, we have

$$p^*_{(0)} = \begin{cases} 1 & \text{if} \quad d_\alpha < -2s \\ 0 & \text{otherwise.} \end{cases} \tag{A16}$$

This result can be understood as follows. Fecundity selection always disfavours the deleterious allele a, but condition dependence generates an association between selfing rate and genotype at the condition locus, so that alleles at this locus experience different selfing rates and so benefit differentially from the transmission advantage of self-fertilisation (Fisher, 1941). Under positive condition dependence ($d_\alpha > 0$), the wild-type allele $\mathrm{A}$ already favoured by fecundity selection enjoys an additional transmission advantage, which further acts against allele $\mathrm{a}$ (eq. A15 becomes more negative). Conversely, negative condition dependence ($d_\alpha < 0$) counteracts fecundity selection by granting a transmission advantage to the deleterious allele $\mathrm{a}$ and offsets it completely when $d_\alpha < -2s$, leading to the near fixation of allele $\mathrm{a}$. We calculate subsequent terms separately for the $p^*_{(0)} = 0$ and $p^*_{(0)} = 1$ cases.

When $p^*_{(0)} = 0$, the first-order perturbation of the change in homozygosity, $\Delta C_{(1)}$, is given by

$$\Delta C_{(1)} = \frac{1}{1+F}\Big(Fp_{(1)} - C_{(1)}\Big), \tag{A17}$$

so that at equilibrium

$$\Delta C_{(1)} = 0 \quad \Rightarrow \quad C^*_{(1)} = Fp^*_{(1)}, \tag{A18}$$

which again corresponds to the expected excess in homozygotes at a diallelic locus under neutrality (with terms in ${p^*_{(1)}}^2$ neglected because they are of order $\epsilon^2$). When $p^*_{(0)} = 1$, on the other hand, we have

$$\Delta C_{(1)} = -\frac{1}{1+F}\Big(Fp_{(1)} + C_{(1)}\Big), \tag{A19}$$

which gives

$$C^*_{(1)} = -Fp^*_{(1)}. \tag{A20}$$

The minus sign in eq. (A20) compensates for the fact that $p_{(0)} = 1$ in this case, so that the first-order perturbation of allelic frequency $p^*_{(1)}$ must be negative or null (see below).

The second-order perturbation of the change in frequency $\Delta p_{(2)}/2$ with $p^*_{(0)} = 0$ ($d_\alpha > -2s$) is given by

$$\frac{\Delta p_{(2)}}{2} = \mu_{\mathrm{A}} - \left(s + \frac{d_\alpha}{2}\right)\Big[h\,(1-F) + F\Big]p_{(1)}, \tag{A21}$$

leading to

$$p^* = p^*_{(1)} = \frac{\mu_\mathrm{A}}{\left(s + \dfrac{d_\alpha}{2}\right)\Big[h\,(1-F) + F\Big]} + \mathcal{O}\left(\epsilon^2\right) \quad \text{and} \quad C^* = Fp^* + \mathcal{O}\left(\epsilon^2\right) \tag{A22}$$

at equilibrium. With $p^*_{(0)} = 1$ ($d_\alpha < -2s$), we find

$$\frac{\Delta p_{(2)}}{2} = -\mu_\mathrm{a} + \left(s + \frac{d_\alpha}{2}\right)\Big[1 - h(1-F)\Big]p_{(1)}, \tag{A23}$$

yielding

$$\frac{\Delta p_{(2)}}{2} = 0 \quad \Rightarrow \quad p^*_{(1)} = \frac{\mu_\mathrm{a}}{\left(s + \dfrac{d_\alpha}{2}\right)\Big[1 - h(1-F)\Big]}, \tag{A24}$$

which is negative because $d_\alpha < -2s$, and so

$$p^* = 1 + \frac{\mu_\mathrm{a}}{\left(s + \dfrac{d_\alpha}{2}\right)\Big[1 - h(1-F)\Big]} + \mathcal{O}\left(\epsilon^2\right) \quad \text{and} \quad C^* = F(1-p^*) + \mathcal{O}\left(\epsilon^2\right). \tag{A25}$$

Eqs. (A22) and (A25) correspond to eq. (2) in the main text.

##### A.1.3 Heterozygote advantage

Polymorphism is always maintained at a balance between mutation and selection in the linear case above, and so would collapse in the absence of recurrent mutation, but we find in numerical analyses that the two alleles at the condition locus can be maintained at an intermediate frequency in the absence of mutation for arbitrary selfing strategies where heterozygotes self-fertilise at a higher rate than homozygotes, i.e., when

$$\alpha_\mathrm{Aa} > \alpha_\mathrm{AA},\ \alpha_\mathrm{aa}. \tag{A26}$$

This is because such strategies generate a form of overdominance at the condition locus that leads to a protected polymorphism. To show this analytically, we now assume that genotype-dependent selfing rates

$\boldsymbol{\alpha} = (\alpha_{\mathrm{AA}}, \alpha_{\mathrm{Aa}}, \alpha_{\mathrm{aa}})$ are given by

$$\boldsymbol{\alpha} = \Big(\alpha_0, \quad \alpha_0 + d_\alpha, \quad \alpha_0\Big), \tag{A27}$$

so that homozygotes share the same selfing rate $\alpha_0$ and heterozygotes may deviate from this rate by an amount $d_\alpha$. Heterozygotes have a higher selfing rate than homozygotes when $d_\alpha > 0$ and a lower selfing rate when $d_\alpha < 0$. We make no assumption on the magnitude of $d_\alpha$ here.

To order zero in $\epsilon$, we find that the change in frequency and excess homozygosity at the condition locus are given by

$$\Delta p_{(0)} = -\frac{d_\alpha}{2}\left(1 - 2p_{(0)}\right)\left(C_{(0)} - p_{(0)}q_{(0)}\right), \tag{A28a}$$

and

$$\Delta C_{(0)} = \frac{\alpha_0}{2} p_{(0)}q_{(0)} - \left(1 - \frac{\alpha}{2}\right) C_{(0)} + \left(\frac{1 - 2p_{(0)}}{2}\right)^2 \left\{1 - \left[1 + d_\alpha\left(C_{(0)} - p_{(0)}q_{(0)}\right)\right]^2\right\} \tag{A28b}$$

where $q_{(0)} = 1 - p_{(0)}$. Solving for equilibrium (eq. A11), we find that this system has three equilibria, either one of the alleles goes to fixation, i.e.,

$$p^*_{(0)} = 0 \quad \text{or} \quad p^*_{(0)} = 1, \tag{A29}$$

in which case $C^*_{(0)} = 0$; or alleles are equally frequent,

$$p^*_{(0)} = \frac{1}{2} \quad \text{and} \quad C^*_{(0)} = \frac{F}{4} \tag{A30}$$

with $F = \alpha_0/(2 - \alpha_0)$. To determine the stability of these equilibria, we compute the Jacobian matrix $\mathbf{J}^*_{(0)}$,

$$\mathbf{J}^*_{(0)} = \begin{pmatrix} \left.\dfrac{\partial p'_{(0)}}{\partial p_{(0)}}\right|_* & \left.\dfrac{\partial p'_{(0)}}{\partial C_{(0)}}\right|_* \\ \left.\dfrac{\partial C'_{(0)}}{\partial p_{(0)}}\right|_* & \left.\dfrac{\partial C'_{(0)}}{\partial C_{(0)}}\right|_* \end{pmatrix}, \tag{A31}$$

where $p'_{(0)} = p_{(0)} + \Delta p_{(0)}$ and $C'_{(0)} = C_{(0)} + \Delta C_{(0)}$ denote allelic frequency and excess homozygosity in the next time step, and the $|_*$ notation indicates that all derivatives are evaluated at an equilibrium of the system (as given by eq. A29 and A30). An equilibrium point is considered to be stable when the leading eigenvalue of $\mathbf{J}^*_{(0)}$, $\lambda_{\mathbf{J}^*_{(0)}}$, has a real part strictly smaller than one in absolute value, i.e.,

$$|\lambda_{\mathbf{J}^*_{(0)}}| < 1. \tag{A32}$$

Checking for Condition (A32) for each of the three equilibria above, we can distinguish two cases. When heterozygotes self-fertilise at a higher rate than homozygotes ($d_\alpha > 0$), the equilibrium where both alleles are maintained at equal frequency is the only stable one (eq. A30), indicating that selection favours their coexistence. When $d_\alpha < 0$, on the other hand, this equilibrium is unstable and both $p^*_{(0)} = 0$ and $p^*_{(0)} = 1$ are stable, meaning that the population will either fix allele $\mathrm{A}$ or allele $\mathrm{a}$ depending on initial conditions. These results can be understood by realising that the average selfing rate associated with an allele becomes strongly frequency-dependent when the rate of heterozygotes differs from homozygotes, because the rarer allele is more likely to be found in heterozygous state. When $d_\alpha > 0$, the rarer allele enjoys a transmission advantage over the other, leading to negative frequency-dependence and the maintenance of polymorphism. Conversely, $d_\alpha < 0$ leads to heterozygote disadvantage and so positive frequency-dependence and the fixation of the most frequent allele. These results neglect the effect of mutation and fecundity selection against $\mathrm{a}$, but since these are small, they are only going to produce small deviations around the equilibria identified here.

#### A.2 Evolution of condition-dependent selfing

We now investigate the evolution of condition-dependent selfing. To do this, we assume that mutations at the modifier locus are rare, occur independently for each of the three traits and have weak, unbiased phenotypic effects. Under these assumptions, evolution at the modifier proceeds gradually, and evolutionary dynamics can be inferred from an invasion analysis (Metz et al., 1996; Geritz et al., 1998; Dercole and Rinaldi, 2008; Avila and Mullon, 2023). We describe our general approach and introduce necessary notations in Section A.2.1. We then solve special cases explicitly in Sections A.2.2-A.2.4, and present complementary numerical analyses in Section A.2.5.

##### A.2.1 Analytical method

We consider the invasion of a rare mutant allele m encoding a strategy $\boldsymbol{\alpha}_{\mathrm{m}} = (\alpha^{\mathrm{m}}_{\mathrm{AA}}, \alpha^{\mathrm{m}}_{\mathrm{Aa}}, \alpha^{\mathrm{m}}_{\mathrm{aa}})$ in a population otherwise fixed with allele M, which encodes a strategy $\boldsymbol{\alpha} = (\alpha_{\mathrm{AA}}, \alpha_{\mathrm{Aa}}, \alpha_{\mathrm{aa}})$. The dynamics of the mutant sub-population can be modelled by the recursion

$$\mathbf{n}_{t+1} = \mathbf{W}(\boldsymbol{\alpha}_{\mathrm{m}}, \boldsymbol{\alpha}) \cdot \mathbf{n}_t, \quad \text{(A33)}$$

where $\mathbf{n}_t = (n_{i,t})_{1\leqslant i\leqslant 8}$ is a vector that gives the number of mutant individuals in each of the eight possible genotypic classes in the sub-population at time $t$ (Table 1), and $\mathbf{W}(\boldsymbol{\alpha}_{\mathrm{m}}, \boldsymbol{\alpha})$ is an $8 \times 8$ matrix of which element $w_{ij}(\boldsymbol{\alpha}_{\mathrm{m}}, \boldsymbol{\alpha})$ gives the expected number of successfully established mutant offspring of class $i$ produced by a focal mutant of class $j$. We hereafter refer to $\mathbf{W}(\boldsymbol{\alpha}_{\mathrm{m}}, \boldsymbol{\alpha})$ as the invasion matrix.

| **Class** | **Genotype** |
|---|---|
| 1 | AM / Am |
| 2 | AM / am |
| 3 | aM / Am |
| 4 | aM / am |
| 5 | Am / Am |
| 6 | Am / am |
| 7 | am / Am |
| 8 | am / am |

**Table 1:** Class number and genotype as used throughout the analysis. For each genotype, the maternally inherited haplotype is given first, followed by the paternal haplotype after the slash symbol.

To facilitate writing the elements of the invasion matrix $\mathbf{W}(\boldsymbol{\alpha}_{\mathrm{m}}, \boldsymbol{\alpha})$, let us first denote as $F^*_x(\boldsymbol{\alpha})$ and $M^*_x(\boldsymbol{\alpha})$ the relative number of female and male gametes available for outcrossing carrying allele $x \in \{\mathrm{A}, \mathrm{a}\}$, respectively, produced by the resident population at its genetic and ecological equilibrium. These variables are given by

$$F_{\mathrm{A}}^*(\boldsymbol{\alpha}) = (1-\mu_{\mathrm{A}})\, q_{\mathrm{AA}}^* \frac{\varphi_{\mathrm{AA}}}{\overline{\varphi}^*}(1-\alpha_{\mathrm{AA}}) + q_{\mathrm{Aa}}^* \frac{1-\mu_{\mathrm{A}}+\mu_{\mathrm{a}}}{2} \frac{\varphi_{\mathrm{Aa}}}{\overline{\varphi}^*}(1-\alpha_{\mathrm{Aa}}) + \mu_{\mathrm{a}}\, q_{\mathrm{aa}}^* \frac{\varphi_{\mathrm{aa}}}{\overline{\varphi}^*}(1-\alpha_{\mathrm{aa}}),$$
$$F_{\mathrm{a}}^*(\boldsymbol{\alpha}) = \mu_{\mathrm{A}}\, q_{\mathrm{AA}}^* \frac{\varphi_{\mathrm{AA}}}{\overline{\varphi}^*}(1-\alpha_{\mathrm{AA}}) + q_{\mathrm{Aa}}^* \frac{1-\mu_{\mathrm{a}}+\mu_{\mathrm{A}}}{2} \frac{\varphi_{\mathrm{Aa}}}{\overline{\varphi}^*}(1-\alpha_{\mathrm{Aa}}) + (1-\mu_{\mathrm{a}})\, q_{\mathrm{aa}}^* \frac{\varphi_{\mathrm{aa}}}{\overline{\varphi}^*}(1-\alpha_{\mathrm{aa}}) \tag{A34a}$$

and

$$M_{\mathrm{A}}^*(\boldsymbol{\alpha}) = (1-\mu_{\mathrm{A}})\, q_{\mathrm{AA}}^* \frac{\varphi_{\mathrm{AA}}}{\overline{\varphi}^*} + q_{\mathrm{Aa}}^* \frac{1-\mu_{\mathrm{A}}+\mu_{\mathrm{a}}}{2} \frac{\varphi_{\mathrm{Aa}}}{\overline{\varphi}^*} + \mu_{\mathrm{a}}\, q_{\mathrm{aa}}^* \frac{\varphi_{\mathrm{aa}}}{\overline{\varphi}^*},$$
$$M_{\mathrm{a}}^*(\boldsymbol{\alpha}) = \mu_{\mathrm{A}}\, q_{\mathrm{AA}}^* \frac{\varphi_{\mathrm{AA}}}{\overline{\varphi}^*} + q_{\mathrm{Aa}}^* \frac{1-\mu_{\mathrm{a}}+\mu_{\mathrm{A}}}{2} \frac{\varphi_{\mathrm{Aa}}}{\overline{\varphi}^*} + (1-\mu_{\mathrm{a}})\, q_{\mathrm{aa}}^* \frac{\varphi_{\mathrm{aa}}}{\overline{\varphi}^*} \tag{A34b}$$

where $q_{x_1x_2}^*$ is the equilibrium frequency of genotype $x_1x_2$ (with $x_1,\ x_2 \in \{\mathrm{A},\mathrm{a}\}$) in the resident population and

$$\overline{\varphi}^* = \varphi_{\mathrm{AA}} q_{\mathrm{AA}}^* + \varphi_{\mathrm{Aa}} q_{\mathrm{Aa}}^* + \varphi_{\mathrm{aa}} q_{\mathrm{aa}}^* \tag{A35}$$

is the mean condition in the resident population. Further, we denote as $\alpha_i$ the selfing rate and $\varphi_i$ the condition of a mutant of class $i$, and by $G_{xy}^{(i)}$ (with $x \in \{\mathrm{A},\mathrm{a}\}$ and $y \in \{\mathrm{M},\mathrm{m}\}$) the proportion of gametes with haplotype $xy$ among those produced by a mutant of class $i$. The expressions for $G_{xy}^{(i)}$ variables are given in Table 2 (see associated caption for explanation).

**Invasion matrix.** Using the above notations, the number of successful mutant offspring of class 1 to 8 produced by a focal mutant of class $i$ are given by

$$\mathbf{w}_i(\boldsymbol{\alpha}_{\mathrm{m}}, \boldsymbol{\alpha}) = \begin{pmatrix} \dfrac{\varphi_i}{\overline{\varphi}^*}\left[2\alpha_i G^{(i)}_{\mathrm{AM}} G^{(i)}_{\mathrm{Am}} + (1-\alpha_i) M^*_{\mathrm{A}}(\boldsymbol{\alpha}) G^{(i)}_{\mathrm{Am}}\right] + F^*_{\mathrm{A}}(\boldsymbol{\alpha}) \dfrac{\varphi_i}{\overline{\varphi}^*} G^{(i)}_{\mathrm{Am}} \\ \dfrac{\varphi_i}{\overline{\varphi}^*}\left[2\alpha_i G^{(i)}_{\mathrm{AM}} G^{(i)}_{\mathrm{am}} + (1-\alpha_i) M^*_{\mathrm{A}}(\boldsymbol{\alpha}) G^{(i)}_{\mathrm{am}}\right] + F^*_{\mathrm{A}}(\boldsymbol{\alpha}) \dfrac{\varphi_i}{\overline{\varphi}^*} G^{(i)}_{\mathrm{am}} \\ \dfrac{\varphi_i}{\overline{\varphi}^*}\left[2\alpha_i G^{(i)}_{\mathrm{Am}} G^{(i)}_{\mathrm{aM}} + (1-\alpha_i) M^*_{\mathrm{a}}(\boldsymbol{\alpha}) G^{(i)}_{\mathrm{Am}}\right] + F^*_{\mathrm{a}}(\boldsymbol{\alpha}) \dfrac{\varphi_i}{\overline{\varphi}^*} G^{(i)}_{\mathrm{Am}} \\ \dfrac{\varphi_i}{\overline{\varphi}^*}\left[2\alpha_i G^{(i)}_{\mathrm{am}} G^{(i)}_{\mathrm{aM}} + (1-\alpha_i) M^*_{\mathrm{a}}(\boldsymbol{\alpha}) G^{(i)}_{\mathrm{am}}\right] + F^*_{\mathrm{a}}(\boldsymbol{\alpha}) \dfrac{\varphi_i}{\overline{\varphi}^*} G^{(i)}_{\mathrm{am}} \\ \dfrac{\varphi_i}{\overline{\varphi}^*} \alpha_i G^{(i)}_{\mathrm{Am}} G^{(i)}_{\mathrm{Am}} \\ \dfrac{\varphi_i}{\overline{\varphi}^*} \alpha_i G^{(i)}_{\mathrm{Am}} G^{(i)}_{\mathrm{am}} \\ \dfrac{\varphi_i}{\overline{\varphi}^*} \alpha_i G^{(i)}_{\mathrm{Am}} G^{(i)}_{\mathrm{am}} \\ \dfrac{\varphi_i}{\overline{\varphi}^*} \alpha_i G^{(i)}_{\mathrm{am}} G^{(i)}_{\mathrm{am}} \end{pmatrix}, \tag{A36}$$

which corresponds to the $i^{\text{th}}$ column of the invasion matrix $\mathbf{W}(\boldsymbol{\alpha}_{\mathrm{m}}, \boldsymbol{\alpha})$, i.e.,

$$\mathbf{W}(\boldsymbol{\alpha}_{\mathrm{m}}, \boldsymbol{\alpha}) = \Big(\mathbf{w}_i(\boldsymbol{\alpha}, \boldsymbol{\alpha}_{\mathrm{m}})\Big)_{1 \leqslant i \leqslant 8}. \tag{A37}$$

To understand how the elements in eq. (A36) were computed, let us focus on its first entry, which gives the number of successfully recruited $\mathrm{AM/Am}$ offspring produced by a mutant of class $i$ and is composed of two terms. The first term gives the number of successful $\mathrm{AM/Am}$ offspring produced by the mutant through its female function. It is proportional to the relative contribution of the mutant to the seed pool through its female function, which is given by its relative condition $\varphi_i/\overline{\varphi}^*$. The mutant self-fertilises a fraction $\alpha_i$ of its ovules, producing a proportion $2G^{(i)}_{\mathrm{AM}}G^{(i)}_{\mathrm{Am}}$ of $\mathrm{AM/Am}$ zygotes. The remaining ovules $(1-\alpha_i)$ are fertilised by resident individuals, in which case an $\mathrm{AM/Am}$ zygote can only be produced if the focal mutant contributes a $\mathrm{Am}$ haplotype (which it does with probability $G^{(i)}_{\mathrm{Am}}$) and receives resident pollen carrying haplotype $\mathrm{AM}$, which occurs with probability $M^*_{\mathrm{A}}(\boldsymbol{\alpha})$. The second term gives the number of successful $\mathrm{AM}/\mathrm{Am}$ offspring produced by the mutant through its male function, i.e. through the fertilisation of resident ovules. Similar to female function, a mutant of class $i$ can only produce $\mathrm{AM}/\mathrm{Am}$ offspring through outcrossing if it fertilises resident ovules carrying haplotype $\mathrm{AM}$. This term is therefore proportional to the contribution of AM ovules to the pool of resident ovules available for outcrossing, $F^{*}_{\mathrm{A}}(\boldsymbol{\alpha})$. The mutant fertilises part of these ovules in proportion to its relative condition $\varphi_i/\overline{\varphi}^{*}$ in which case $\mathrm{AM}/\mathrm{Am}$ offspring are formed when the mutant transmits haplotype $\mathrm{Am}$, which occurs with probability $G^{(i)}_{\mathrm{Am}}$.

| **Class** $i$ | $G^{(i)}_{\mathrm{AM}}$ | $G^{(i)}_{\mathrm{Am}}$ | $G^{(i)}_{\mathrm{aM}}$ | $G^{(i)}_{\mathrm{am}}$ |
|---|---|---|---|---|
| 1 | $\frac{1-\mu_{\mathrm{A}}}{2}$ | $\frac{1-\mu_{\mathrm{A}}}{2}$ | $\frac{\mu_{\mathrm{A}}}{2}$ | $\frac{\mu_{\mathrm{A}}}{2}$ |
| 2 | $\frac{(1-r)(1-\mu_{\mathrm{A}})+r\mu_{\mathrm{a}}}{2}$ | $\frac{r(1-\mu_{\mathrm{A}})+(1-r)\mu_{\mathrm{a}}}{2}$ | $\frac{r(1-\mu_{\mathrm{a}})+(1-r)\mu_{\mathrm{A}}}{2}$ | $\frac{(1-r)(1-\mu_{\mathrm{a}})+r\mu_{\mathrm{A}}}{2}$ |
| 3 | $\frac{r(1-\mu_{\mathrm{A}})+(1-r)\mu_{\mathrm{a}}}{2}$ | $\frac{(1-r)(1-\mu_{\mathrm{A}})+r\mu_{\mathrm{a}}}{2}$ | $\frac{(1-r)(1-\mu_{\mathrm{A}})+r\mu_{\mathrm{a}}}{2}$ | $\frac{r(1-\mu_{\mathrm{a}})+(1-r)\mu_{\mathrm{A}}}{2}$ |
| 4 | $\frac{\mu_{\mathrm{a}}}{2}$ | $\frac{\mu_{\mathrm{a}}}{2}$ | $\frac{1-\mu_{\mathrm{a}}}{2}$ | $\frac{1-\mu_{\mathrm{a}}}{2}$ |
| 5 | 0 | $1-\mu_{\mathrm{A}}$ | 0 | $\mu_{\mathrm{A}}$ |
| 6 | 0 | $\frac{1-\mu_{\mathrm{A}}+\mu_{\mathrm{a}}}{2}$ | 0 | $\frac{1-\mu_{\mathrm{a}}+\mu_{\mathrm{A}}}{2}$ |
| 7 | 0 | $\frac{1-\mu_{\mathrm{A}}+\mu_{\mathrm{a}}}{2}$ | 0 | $\frac{1-\mu_{\mathrm{a}}+\mu_{\mathrm{A}}}{2}$ |
| 8 | 0 | $\mu_{\mathrm{a}}$ | 0 | $1-\mu_{\mathrm{a}}$ |

**Table 2:** Proportion of gametes carrying genotype $xy$, $x \in \{\mathrm{A},\mathrm{a}\}$, $y \in \{\mathrm{M},\mathrm{m}\}$ among the gametes produced by a mutant individual of class $i \in \{1,...,8\}$. These expressions assume that recombination occurs first, followed by mutation at the condition locus. To understand how they were obtained, consider the gametes produced by an individual of class 2, i.e., with genotype $\mathrm{AM}/\mathrm{am}$. Following recombination, such an individual will produce haplotype AM, Am, aM and am in proportions $(1-r)/2$, $r/2$, $r/2$ and $(1-r)/2$, respectively. Alleles at the condition locus then each mutate with probability $\mu_x$, $x \in \{\mathrm{A},\mathrm{a}\}$, so that haplotype AM for instance is produced either from AM recombinant haplotypes that did not mutate, which occurs with probability $(1-r)(1-\mu_{\mathrm{A}})/2$; or from a aM that mutated, which occurs with probability $r\mu_{\mathrm{a}}/2$. Repeating this reasoning for each haplotype gives the proportions in the table.

**Selection gradient in a class-structured population.** The invasion fitness $\rho(\boldsymbol{\alpha}_{\mathrm{m}},\boldsymbol{\alpha})$ of a mutant $\boldsymbol{\alpha}_{\mathrm{m}}$ in a population fixed for $\boldsymbol{\alpha}$, from which the selection gradients on $\alpha_{\mathrm{AA}}$, $\alpha_{\mathrm{Aa}}$ and $\alpha_{\mathrm{aa}}$ can be computed, is given by the leading eigenvalue of the invasion matrix $\mathbf{W}(\boldsymbol{\alpha}_{\mathrm{m}},\boldsymbol{\alpha})$ (Avila and Mullon, 2023). Unfortunately, this eigenvalue cannot be computed explicitly and so cannot directly be used to characterise the evolution of condition-dependent selfing. Under the assumption that mutations have weak phenotypic effects, however, the selection gradients $\mathbf{S}(\boldsymbol{\alpha}) = (S_{\mathrm{AA}}(\boldsymbol{\alpha}), S_{\mathrm{Aa}}(\boldsymbol{\alpha}), S_{\mathrm{aa}}(\boldsymbol{\alpha}))$ on condition-dependent selfing rates can be obtained as

$$S_{x_1x_2}(\boldsymbol{\alpha}) = \mathbf{v}^{\circ}(\boldsymbol{\alpha}) \cdot \mathbf{D}_{x_1x_2}(\boldsymbol{\alpha}) \cdot \mathbf{q}^{\circ}(\boldsymbol{\alpha}), \tag{A38}$$

where $\mathbf{q}^{\circ}(\boldsymbol{\alpha}) = \left(q_i^{\circ}(\boldsymbol{\alpha})\right)_{1\leqslant i\leqslant 8}$ is the right eigenvector of $\mathbf{W}^{\circ}(\boldsymbol{\alpha}) = \mathbf{W}(\boldsymbol{\alpha}, \boldsymbol{\alpha})$, the invasion matrix under neutrality, normalised such that

$$\sum_{i=1}^{8} q_i^{\circ}(\boldsymbol{\alpha}) = 1,$$

which gives the asymptotic frequencies of classes in the population under neutrality; $\mathbf{D}_{x_1x_2}(\boldsymbol{\alpha})$ is an $8 \times 8$ matrix, the $(i, j)$-entry of which $d_{ij}^{x_1x_2}(\boldsymbol{\alpha})$ is given by the derivative of the corresponding entry of the invasion matrix $\mathbf{W}(\boldsymbol{\alpha}_{\mathrm{m}}, \boldsymbol{\alpha})$ with respect to $\alpha_{x_1x_2}^{\mathrm{m}}$ evaluated at the resident strategy, i.e.,

$$d_{ij}^{x_1x_2}(\boldsymbol{\alpha}) = \left.\frac{\partial w_{ij}(\boldsymbol{\alpha}_{\mathrm{m}}, \boldsymbol{\alpha})}{\partial \alpha_{x_1x_2}^{\mathrm{m}}}\right|_{\boldsymbol{\alpha}_{\mathrm{m}}=\boldsymbol{\alpha}}, \tag{A39}$$

which measures the effect of a change in trait $\alpha_{x_1x_2}$ on the number of successful offspring of class $i$ produced by a focal mutant of class $j$; and $\mathbf{v}^{\circ}(\boldsymbol{\alpha}) = \left(v_i^{\circ}(\boldsymbol{\alpha})\right)_{1\leqslant i\leqslant 8}$ is the left eigenvector of $\mathbf{W}^{\circ}(\boldsymbol{\alpha}) = \mathbf{W}(\boldsymbol{\alpha}, \boldsymbol{\alpha})$, normalised such that

$$\mathbf{v}^{\circ}(\boldsymbol{\alpha}) \cdot \mathbf{q}^{\circ}(\boldsymbol{\alpha}) = 1,$$

which gives the reproductive value of an individual of class $i$ under neutrality, that is its long-term contribution to population growth. Background for this decomposition can be found, e.g., in Taylor (1990) or Chap. 11 in Rousset (2004). See also Section 3 in Avila and Mullon (2023) for a review. In what follows, we use this decomposition to study the evolution of condition-dependent selfing.

**Weak variation in condition.** Similar to our calculations under fixed condition-dependent selfing (Section A.1), we assume that $s \sim \mathcal{O}(\epsilon)$ and $\mu_x \sim \mathcal{O}(\epsilon^2)$ in order to obtain analytical results. To incorporate these assumptions, our approach will again rely on Taylor-expanding relevant quantities using eq. (A8).

Applying this method to the selection gradient on $\alpha_{x_1x_2}$, we find that it can be expressed as

$$\begin{aligned} S_{x_1x_2}(\boldsymbol{\alpha}) = {} & \mathbf{v}^{\circ}_{(0)} \cdot \mathbf{D}^{x_1x_2}_{(0)} \cdot \mathbf{q}^{\circ}_{(0)} \\ & + \epsilon\left(\mathbf{v}^{\circ}_{(1)} \cdot \mathbf{D}^{x_1x_2}_{(0)} \cdot \mathbf{q}^{\circ}_{(0)} + \mathbf{v}^{\circ}_{(0)} \cdot \mathbf{D}^{x_1x_2}_{(1)} \cdot \mathbf{q}^{\circ}_{(0)} + \mathbf{v}^{\circ}_{(0)} \cdot \mathbf{D}^{x_1x_2}_{(0)} \cdot \mathbf{q}^{\circ}_{(1)}\right) \\ & + \frac{\epsilon^2}{2}\Bigg[\mathbf{v}^{\circ}_{(2)} \cdot \mathbf{D}^{x_1x_2}_{(0)} \cdot \mathbf{q}^{\circ}_{(0)} + \mathbf{v}^{\circ}_{(0)} \cdot \mathbf{D}^{x_1x_2}_{(2)} \cdot \mathbf{q}^{\circ}_{(0)} + \mathbf{v}^{\circ}_{(0)} \cdot \mathbf{D}^{x_1x_2}_{(0)} \cdot \mathbf{q}^{\circ}_{(2)} \\ & \qquad + 2\left(\mathbf{v}^{\circ}_{(1)} \cdot \mathbf{D}^{x_1x_2}_{(1)} \cdot \mathbf{q}^{\circ}_{(0)} + \mathbf{v}^{\circ}_{(1)} \cdot \mathbf{D}^{x_1x_2}_{(0)} \cdot \mathbf{q}^{\circ}_{(1)} + \mathbf{v}^{\circ}_{(0)} \cdot \mathbf{D}^{x_1x_2}_{(1)} \cdot \mathbf{q}^{\circ}_{(1)}\right)\Bigg] \\ & + \mathcal{O}\left(\epsilon^3\right) \end{aligned} \tag{A40}$$

for $x_1, x_2 \in \{\mathrm{A}, \mathrm{a}\}$, where $\mathbf{v}^{\circ}_{(k)}$, $\mathbf{D}^{x_1x_2}_{(k)}$ and $\mathbf{q}^{\circ}_{(k)}$, $k \in \{0, 1, ..., n\}$ are $k^{\text{th}}$ order perturbations of $\mathbf{v}^{\circ}$, the vector of individual reproductive values under neutrality; of $\mathbf{D}_{x_1x_2}(\boldsymbol{\alpha})$, the matrix of effects; and of $\mathbf{q}^{\circ}$, the vector of asymptotic class frequencies under neutrality, respectively.

Our aim will be to compute the perturbation terms that appear in eq. (A40) to increasing order in $\epsilon$, starting with order zero (first line) and stopping at the first non-zero term (the *leading* order).

##### A.2.2 Complete outcrossing

Computing the selection gradients on genotype-specific selfing rates for an arbitrary strategy $\boldsymbol{\alpha}$ is difficult. We therefore start by investigating special cases. Here, we compute a leading order approximation for the selection gradients on the selfing rates in a completely outcrossing population ($\alpha_{\mathrm{AA}} = \alpha_{\mathrm{Aa}} = \alpha_{\mathrm{aa}} = 0$). We first solve for the genetic equilibrium reached by an outcrossing resident population and then compute the selection gradients.

**Resident equilibrium.** To compute the genetic equilibrium in the resident population, we use the same method as in Appendix A.1.2, i.e., we assume that $s \sim \mathcal{O}(\epsilon)$ and $\mu_x \sim \mathcal{O}\left(\epsilon^2\right)$, and expand the allelic frequency $p(t)$ and excess in homozygotes $C(t)$, and their change between generations around $\epsilon$ with $\alpha_{\mathrm{AA}} = \alpha_{\mathrm{Aa}} = \alpha_{\mathrm{aa}} = 0$ (eqs. A10-A11).

We find that the change in allelic frequency and homozygosity are given by

$$\Delta p_{(0)} = 0 \quad \text{and} \quad \Delta C_{(0)} = -C_{(0)} \tag{A41}$$

to zero-th order in $\epsilon$. This indicates that in the absence of selection or mutation, the change frequency of allele $\mathrm{a}$ is expected to be zero, so that the population will retain the same allelic frequency indefinitely in the absence of drift, meanwhile the excess in homozygotes is expected to be zero in a strictly outcrossing population at equilibrium, as we have

$$\Delta C_{(0)} = 0 \quad \Leftrightarrow \quad C^{*}_{(0)} = 0. \tag{A42}$$

To first order in $\epsilon$, the changes in allelic frequency and excess in homozygotes are given by

$$\Delta p_{(1)} = -s\left[p_{(0)} + h\left(1 - 2p_{(0)}\right)\right] p_{(0)}\left(1 - p_{(0)}\right) \quad \text{and} \quad \Delta C_{(1)} = -C_{(1)}, \tag{A43}$$

which at equilibrium yields

$$p^{*}_{(0)} = 0 \quad \text{and} \quad C^{*}_{(1)} = 0. \tag{A44}$$

Eq. (A43) reveals that selection against allele $\mathrm{a}$ always lowers its frequency in the population and drives it to extinction in the absence of mutation – which does not feature in the change in allelic frequency to first order in $\epsilon$ because $\mu_x \sim \mathcal{O}\left(\epsilon^2\right)$ – so that the equilibrium frequency of allele $\mathrm{a}$ to leading order in $\epsilon$ is expected to be zero.

To second order in $\epsilon$, we have

$$\Delta p_{(2)} = 2\left(\mu_{\mathrm{A}} - shp_{(1)}\right) \quad \text{and} \quad \Delta C_{(2)} = -C_{(2)}, \tag{A45}$$

which gives

$$p^{*}_{(1)} = \frac{\mu_{\mathrm{A}}}{sh} \quad \text{and} \quad C^{*}_{(2)} = 0 \tag{A46}$$

at equilibrium. This corresponds to the classical result of Haldane (1937) for $\mu_{\mathrm{A}} \ll s$. The deleterious allele $\mathrm{a}$ is maintained at a balance between mutation and selection and the population is at Hardy-Weinberg equilibrium among newborns owing to random mating.

Finally, to third order in $\epsilon$,

$$\Delta p_{(3)} = -3\left(shp_{(2)} + 2(1-h)\frac{\mu_{\mathrm{A}}^2}{sh^2}\right), \tag{A47}$$

which yields

$$p^*_{(2)} = 2\left(1 - \frac{1}{h}\right)\left(\frac{\mu_{\mathrm{A}}}{sh}\right)^2. \tag{A48}$$

Thus, when selection is weak and mutation is rare,

$$p^* = \frac{\mu_{\mathrm{A}}}{sh}\left[1 + \left(1 - \frac{1}{h}\right)\frac{\mu_{\mathrm{A}}}{sh}\right] + \mathcal{O}\left(\epsilon^3\right) \quad \text{and} \quad C^* = 0 + \mathcal{O}\left(\epsilon^3\right) \tag{A49}$$

under complete outcrossing.

**Selection gradients to order zero.** The right and left eigenvectors of the invasion matrix under neutrality $\mathbf{W}^\circ(\boldsymbol{\alpha})$, $\mathbf{q}^\circ$ and $\mathbf{v}^\circ$, which give the asymptotic class frequencies and class-specific reproductive values in the mutant population under neutrality, are defined as the solutions of

$$\mathbf{W}^\circ(\boldsymbol{\alpha}) \cdot \mathbf{q}^\circ = \mathbf{q}^\circ \quad \text{and} \quad \mathbf{v}^\circ \cdot \mathbf{W}^\circ(\boldsymbol{\alpha}) = \mathbf{v}^\circ, \tag{A50}$$

which must hold to all orders in $\epsilon$. Thus, their zero-th order perturbations $\mathbf{q}^\circ_{(0)}$ and $\mathbf{v}^\circ_{(0)}$ must satisfy

$$\mathbf{W}^\circ_{(0)} \cdot \mathbf{q}^\circ_{(0)} = \mathbf{q}^\circ_{(0)} \quad \text{and} \quad \mathbf{v}^\circ_{(0)} \cdot \mathbf{W}^\circ_{(0)} = \mathbf{v}^\circ_{(0)}, \tag{A51}$$

with constraints $\mathbf{q}^\circ_{(0)} \cdot \mathbf{1} = 1$ and $\mathbf{v}^\circ_{(0)} \cdot \mathbf{q}^\circ_{(0)} = 1$, and where $\mathbf{W}^\circ_{(0)}$ is the zero-th order perturbation of $\mathbf{W}^\circ(\boldsymbol{\alpha})$. Solving eq. (A51) for $\mathbf{q}^\circ_{(0)}$ and $\mathbf{v}^\circ_{(0)}$, we obtain

$$\mathbf{q}^\circ_{(0)} = (1,\ 0,\ 0,\ 0,\ 0,\ 0,\ 0,\ 0) \quad \text{and} \quad \mathbf{v}^\circ_{(0)} = (1,\ 1,\ 1,\ 1,\ 2,\ 2,\ 2,\ 2). \tag{A52}$$

This indicates that to order zero in $\epsilon$, the mutant population under neutrality is expected to be fixed for allele A at the condition locus ($q^{\circ}_{1(0)} = 1$, which stems from the fact that $p^{*}_{(0)} = 0$ and homozygotes for the mutant allele at the modifier cannot be produced under complete outcrossing), and the reproductive value of an individual homozygous for the mutant allele is exactly twice that of an heterozygous individual, because it carries twice as many mutant copies.

The matrices of fitness effects to order zero, meanwhile, are given by

$$\mathbf{D}^{\mathrm{AA}}_{(0)} = \begin{pmatrix} 0 & 0 & 0 & 0 & -1 & 0 & 0 & 0 \\ 0 & 0 & 0 & 0 & 0 & 0 & 0 & 0 \\ 0 & 0 & 0 & 0 & 0 & 0 & 0 & 0 \\ 0 & 0 & 0 & 0 & 0 & 0 & 0 & 0 \\ \frac{1}{8} & 0 & 0 & 0 & 1 & 0 & 0 & 0 \\ 0 & 0 & 0 & 0 & 0 & 0 & 0 & 0 \\ 0 & 0 & 0 & 0 & 0 & 0 & 0 & 0 \\ 0 & 0 & 0 & 0 & 0 & 0 & 0 & 0 \end{pmatrix}, \tag{A53a}$$

$$
\mathbf{D}^{\mathrm{Aa}}_{(0)} = \begin{pmatrix}
0 & -\frac{r^2}{4} & -\frac{(1-r)^2}{4} & 0 & 0 & -\frac{1}{2} & -\frac{1}{2} & 0 \\
0 & -\frac{r(1-r)}{4} & -\frac{r(1-r)}{4} & 0 & 0 & -\frac{1}{2} & -\frac{1}{2} & 0 \\
0 & \frac{r^2}{4} & \frac{(1-r)^2}{4} & 0 & 0 & 0 & 0 & 0 \\
0 & \frac{r(1-r)}{4} & \frac{r(1-r)}{4} & 0 & 0 & 0 & 0 & 0 \\
0 & \frac{r^2}{8} & \frac{(1-r)^2}{8} & 0 & 0 & \frac{1}{4} & \frac{1}{4} & 0 \\
0 & \frac{r(1-r)}{8} & \frac{r(1-r)}{8} & 0 & 0 & \frac{1}{4} & \frac{1}{4} & 0 \\
0 & \frac{r(1-r)}{8} & \frac{r(1-r)}{8} & 0 & 0 & \frac{1}{4} & \frac{1}{4} & 0 \\
0 & \frac{(1-r)^2}{8} & \frac{r^2}{8} & 0 & 0 & \frac{1}{4} & \frac{1}{4} & 0
\end{pmatrix}, \tag{A53b}
$$

and

$$
\mathbf{D}^{\mathrm{aa}}_{(0)} = \begin{pmatrix}
0 & 0 & 0 & 0 & 0 & 0 & 0 & 0 \\
0 & 0 & 0 & -\frac{1}{4} & 0 & 0 & 0 & -1 \\
0 & 0 & 0 & 0 & 0 & 0 & 0 & 0 \\
0 & 0 & 0 & \frac{1}{4} & 0 & 0 & 0 & 0 \\
0 & 0 & 0 & 0 & 0 & 0 & 0 & 0 \\
0 & 0 & 0 & 0 & 0 & 0 & 0 & 0 \\
0 & 0 & 0 & 0 & 0 & 0 & 0 & 0 \\
0 & 0 & 0 & \frac{1}{8} & 0 & 0 & 0 & 1
\end{pmatrix}. \tag{A53c}
$$

Inserting eqs. (A52) and (A53) into eq. (A40), we have

$$S_{\mathrm{AA}}\left(\mathbf{0}\right) = \frac{1}{4} + \mathcal{O}\left(\epsilon\right) \tag{A54}$$

Selection on $\alpha_{\mathrm{AA}}$ therefore always favours an increase in the selfing rate of wild-type homozygotes in a completely outcrossing, and lower order terms need not be considered for this genotype. The gradients for heterozygotes and deleterious homozygotes, meanwhile, are given by

$$S_{\mathrm{Aa}}\left(\mathbf{0}\right) = S_{\mathrm{aa}}\left(\mathbf{0}\right) = 0 + \mathcal{O}\left(\epsilon\right), \tag{A55}$$

indicating that higher order terms must be computed.

**Selection gradients to order $\epsilon$.** To first order in $\epsilon$, eqs. (A50) become

$$\mathbf{W}^{\circ}_{(1)} \cdot \mathbf{q}^{\circ}_{(0)} + \mathbf{W}^{\circ}_{(0)} \cdot \mathbf{q}^{\circ}_{(1)} = \mathbf{q}^{\circ}_{(1)} \quad \text{and} \quad \mathbf{v}^{\circ}_{(0)} \cdot \mathbf{W}^{\circ}_{(1)} + \mathbf{v}^{\circ}_{(1)} \cdot \mathbf{W}^{\circ}_{(0)} = \mathbf{v}^{\circ}_{(1)}, \tag{A56}$$

and the renormalisation constraints imposed upon $\mathbf{q}^{\circ}$ and $\mathbf{v}^{\circ}$ give

$$\mathbf{q}^{\circ}_{(1)} \cdot \mathbf{1} = 0 \quad \text{and} \quad \mathbf{v}^{\circ}_{(1)} \cdot \mathbf{q}^{\circ}_{(0)} + \mathbf{v}^{\circ}_{(0)} \cdot \mathbf{q}^{\circ}_{(1)} = 0 \tag{A57}$$

for first order perturbations $\mathbf{q}^{\circ}_{(1)}$ and $\mathbf{v}^{\circ}_{(1)}$. Solving eqs. (A56) and (A57) for $\mathbf{q}^{\circ}_{(1)}$ and $\mathbf{v}^{\circ}_{(1)}$ yields

$$\mathbf{q}^{\circ}_{(1)} = \left(-2\frac{\mu_{\mathrm{A}}}{sh},\ \frac{\mu_{\mathrm{A}}}{sh},\ \frac{\mu_{\mathrm{A}}}{sh},\ 0,\ 0,\ 0,\ 0,\ 0\right), \tag{A58a}$$

and

$$\mathbf{v}^{\circ}_{(1)} = \left(0,\ -\frac{sh}{r},\ -2sh,\ -s\left(1+\frac{h}{r}\right),\ 0,\ -sh\frac{1+2r}{r},\ -sh\frac{1+2r}{r}, -2s\left(1+\frac{h}{r}\right)\right). \tag{A58b}$$

Meanwhile, the first-order perturbations of matrices $\mathbf{D}_{\mathrm{Aa}}(\boldsymbol{\alpha})$ and $\mathbf{D}_{\mathrm{aa}}(\boldsymbol{\alpha})$, $\mathbf{D}^{\mathrm{Aa}}_{(1)}$ and $\mathbf{D}^{\mathrm{aa}}_{(1)}$, are given by

$$\mathbf{D}^{\mathrm{Aa}}_{(1)} = \begin{pmatrix}
0 & \frac{r\left[r(sh)^2+\mu_{\mathrm{A}}\right]}{4sh} & \frac{(1-r)\left[(1-r)(sh)^2+\mu_{\mathrm{A}}\right]}{4sh} & 0 & 0 & \frac{(sh)^2+\mu_{\mathrm{A}}}{2sh} & \frac{(sh)^2+\mu_{\mathrm{A}}}{2sh} & 0 \\
0 & \frac{(1-r)\left(r(sh)^2+\mu_{\mathrm{A}}\right)}{4sh} & \frac{r\left[(1-r)(sh)^2+\mu_{\mathrm{A}}\right]}{4sh} & 0 & 0 & \frac{(sh)^2+\mu_{\mathrm{A}}}{2sh} & \frac{(sh)^2+\mu_{\mathrm{A}}}{2sh} & 0 \\
0 & -\frac{r\left(r(sh)^2+\mu_{\mathrm{A}}\right)}{4sh} & -\frac{(1-r)\left[(1-r)(sh)^2+\mu_{\mathrm{A}}\right]}{4sh} & 0 & 0 & -\frac{\mu_{\mathrm{A}}}{2sh} & -\frac{\mu_{\mathrm{A}}}{2sh} & 0 \\
0 & -\frac{(1-r)\left(r(sh)^2+\mu_{\mathrm{A}}\right)}{4sh} & -\frac{r\left[(1-r)(sh)^2+\mu_{\mathrm{A}}\right]}{4sh} & 0 & 0 & -\frac{\mu_{\mathrm{A}}}{2sh} & -\frac{\mu_{\mathrm{A}}}{2sh} & 0 \\
0 & -\frac{shr^2}{8} & -\frac{sh(1-r)^2}{8} & 0 & 0 & -\frac{sh}{4} & -\frac{sh}{4} & 0 \\
0 & -\frac{shr(1-r)}{8} & -\frac{shr(1-r)}{8} & 0 & 0 & -\frac{sh}{4} & -\frac{sh}{4} & 0 \\
0 & -\frac{shr(1-r)}{8} & -\frac{shr(1-r)}{8} & 0 & 0 & -\frac{sh}{4} & -\frac{sh}{4} & 0 \\
0 & -\frac{sh(1-r)^2}{8} & -\frac{shr^2}{8} & 0 & 0 & -\frac{sh}{4} & -\frac{sh}{4} & 0
\end{pmatrix}, \tag{A59a}$$

and

$$\mathbf{D}^{\mathrm{aa}}_{(1)} = \begin{pmatrix} 0 & 0 & 0 & 0 & 0 & 0 & 0 & 0 \\ 0 & 0 & 0 & \frac{1}{4}\left(s+\frac{\mu_{\mathrm{A}}}{sh}\right) & 0 & 0 & 0 & s+\frac{\mu_{\mathrm{A}}}{sh} \\ 0 & 0 & 0 & 0 & 0 & 0 & 0 & 0 \\ 0 & 0 & 0 & -\frac{1}{4}\left(s+\frac{\mu_{\mathrm{A}}}{sh}\right) & 0 & 0 & 0 & -\frac{\mu_{\mathrm{A}}}{sh} \\ 0 & 0 & 0 & 0 & 0 & 0 & 0 & 0 \\ 0 & 0 & 0 & 0 & 0 & 0 & 0 & 0 \\ 0 & 0 & 0 & 0 & 0 & 0 & 0 & 0 \\ 0 & 0 & 0 & -\frac{s}{8} & 0 & 0 & 0 & -s \end{pmatrix}. \tag{A59b}$$

Using eqs. (A58) and (A59), the selection gradient on $\alpha_{\mathrm{Aa}}$ becomes

$$S_{\mathrm{Aa}}(\mathbf{0}) = \frac{\mu_{\mathrm{A}}}{2sh} + \mathcal{O}\left(\epsilon^2\right) \tag{A60}$$

to first order in $\epsilon$, which shows that selection always favours an increase in the selfing rate of heterozygotes under complete outcrossing; whereas the selection gradient on $\alpha_{\mathrm{aa}}$ becomes

$$S_{\mathrm{aa}}(\mathbf{0}) = 0 + \mathcal{O}\left(\epsilon^2\right), \tag{A61}$$

indicating that higher order terms must be computed.

**Selection gradients to order $\epsilon^2$.** To compute the selection gradient on $\alpha_{\mathrm{aa}}$ to second order in $\epsilon$, we first remark that

$$\mathbf{D}^{\mathrm{aa}}_{(0)} \cdot \mathbf{q}^{\circ}_{(0)} = (0,\ 0,\ 0,\ 0,\ 0,\ 0,\ 0,\ 0), \tag{A62}$$

which implies that

$$\mathbf{v}^{\circ}_{(2)} \cdot \mathbf{D}^{\mathrm{aa}}_{(0)} \cdot \mathbf{q}^{\circ}_{(0)} = 0$$

irrespective of the elements of $\mathbf{v}^{\circ}_{(2)}$, so that we need not compute it. The second-order matrix of effects $\mathbf{D}^{\mathrm{aa}}_{(2)}$ is complicated, but can be found in the accompanying *Mathematica* notebook (available at `10.5281/zenodo.18523420`). Using this matrix, we have

$$\mathbf{v}^{\circ}_{(0)} \cdot \mathbf{D}^{\mathrm{aa}}_{(2)} \cdot \mathbf{q}^{\circ}_{(0)} = 0.$$

Using eqs. (A58- A59), we find that the selection gradient on $\alpha_{\mathrm{aa}}$ reduces to

$$S_{\mathrm{aa}}(\mathbf{0}) = \frac{\mathbf{v}^{\circ}_{(0)} \cdot \mathbf{D}^{\mathrm{aa}}_{(0)} \cdot \mathbf{q}^{\circ}_{(2)}}{2} + \mathcal{O}\left(\epsilon^3\right). \tag{A63}$$

Thus, all that is left to do is to compute the second-order perturbation $\mathbf{q}^{\circ}_{(2)}$. From eq. (A50), $\mathbf{q}^{\circ}_{(2)}$ is the solution of

$$\mathbf{W}^{\circ}_{(2)} \cdot \mathbf{q}^{\circ}_{(0)} + \mathbf{W}^{\circ}_{(0)} \cdot \mathbf{q}^{\circ}_{(2)} + 2\mathbf{W}^{\circ}_{(1)} \cdot \mathbf{q}^{\circ}_{(1)} = \mathbf{q}^{\circ}_{(2)}, \tag{A64}$$

subject to the constraint

$$\mathbf{q}^{\circ}_{(2)} \cdot \mathbf{1} = 0. \tag{A65}$$

Solving eq. (A64) for $\mathbf{q}^{\circ}_{(2)}$ yields

$$\begin{aligned}\mathbf{q}^{\circ}_{(2)} = \Bigg(&-2\left(\frac{\mu_{\mathrm{A}}}{sh}\right)^2\left[1-\frac{2}{h}+2\left(1-\frac{\mu_{\mathrm{a}}}{\mu_{\mathrm{A}}}\right)\right],\ -2\frac{h(\mu_{\mathrm{a}}-\mu_{\mathrm{A}})+\mu_{\mathrm{A}}}{\mu_{\mathrm{A}}h}\left(\frac{\mu_{\mathrm{A}}}{sh}\right)^2, \\ &-2\frac{h(\mu_{\mathrm{a}}-\mu_{\mathrm{A}})+\mu_{\mathrm{A}}}{\mu_{\mathrm{A}}h}\left(\frac{\mu_{\mathrm{A}}}{sh}\right)^2,\ 2\left(\frac{\mu_{\mathrm{A}}}{sh}\right)^2,\ 0,\ 0,\ 0,\ 0\Bigg),\end{aligned} \tag{A66}$$

and the selection gradient on $\alpha_{\mathrm{aa}}$ becomes

$$S_{\mathrm{aa}}(\mathbf{0}) = \frac{1}{4}\left(\frac{\mu_{\mathrm{A}}}{sh}\right)^2 + \mathcal{O}\left(\epsilon^3\right). \tag{A67}$$

**Summary.** In this section, we have shown that selection gradients on genotype-specific selfing rates under complete outcrossing are given by

$$\mathbf{s}(\mathbf{0}) = \left(\frac{1}{4} \times 1 + \mathcal{O}(\epsilon), \quad \frac{1}{4} \times 2\frac{\mu_{\mathrm{A}}}{sh} + \mathcal{O}\left(\epsilon^2\right), \quad \frac{1}{4} \times \left(\frac{\mu_{\mathrm{A}}}{sh}\right)^2 + \mathcal{O}\left(\epsilon^3\right)\right) \tag{A68}$$

to leading order (eqs. A54, A60 and A67). These can be shown to be proportional to the equilibrium frequency of the corresponding genotype at the condition locus multiplied by a factor $1/4$, i.e.,

$$S_{x_1 x_2}(\mathbf{0}) = \frac{q^*_{x_1 x_2}}{4}, \tag{A69}$$

$\mathrm{x}_1, \mathrm{x}_2 \in \{\mathrm{A}, \mathrm{a}\}$ to leading order.

##### A.2.3 Complete selfing

Here, we compute a leading order approximation for the selection gradients on the selfing rates in a fully selfing population ($\alpha_{\mathrm{AA}} = \alpha_{\mathrm{Aa}} = \alpha_{\mathrm{aa}} = 1$). Similar to the previous section, we first solve for the genetic equilibrium reached by the resident population and then compute the selection gradients.

**Resident equilibrium.** Using the same decomposition as above (eq. A10), we find that the change in frequency and in excess homozygosity at the condition locus are given by

$$\Delta p_{(0)} = 0 \quad \text{and} \quad \Delta C_{(0)} = \frac{1}{2}\left[p_{(0)}\left(1 - p_{(0)}\right) - C_{(0)}\right] \tag{A70}$$

to order zero in $\epsilon$. Similar to the complete outcrossing case, the change in frequency at the condition locus is null, because the effects of selection and mutation are both neglected to order zero. Meanwhile, we find that the equilibrium value of $C_{(0)}$ is simply given by

$$\Delta C_{(0)} = 0 \quad \Leftrightarrow \quad C^*_{(0)} = p^*_{(0)}\left(1 - p^*_{(0)}\right). \tag{A71}$$

To first order in $\epsilon$, selection effects appear and we obtain

$$\Delta p_{(1)} = -sp_{(0)}\left(1 - p_{(0)}\right) \quad \text{and} \quad \Delta C_{(1)} = \frac{1}{2}\left(p_{(1)} - C_{(1)}\right), \tag{A72}$$

which yields

$$p^*_{(0)} = 0 \quad \Rightarrow \quad C^*_{(0)} = 0, \tag{A73}$$

and

$$C^*_{(1)} = p^*_{(1)} \tag{A74}$$

at equilibrium. Down to order $\epsilon^2$, we then find that the change in frequency and excess homozygosity at the condition locus read

$$\Delta p_{(2)} = 2\left(\mu_{\mathrm{A}} - sp_{(1)}\right) \quad \text{and} \quad \Delta C_{(2)} = \frac{1}{2}\left[p_{(2)} - C_{(2)} - 2\mu_{\mathrm{A}}\left(2 + \frac{\mu_{\mathrm{A}}}{s^2}\right)\right], \tag{A75}$$

so that at equilibrium

$$p^*_{(1)} = \frac{\mu_{\mathrm{A}}}{s} \quad \Rightarrow \quad C^*_{(1)} = \frac{\mu_{\mathrm{A}}}{s}, \quad \text{and} \quad C^*_{(2)} = p_{(2)} - 2\mu_{\mathrm{A}}\left(2 + \frac{\mu_{\mathrm{A}}}{s^2}\right). \tag{A76}$$

Finally, the third order perturbation of $\Delta p$ is given by

$$\Delta p_{(3)} = -3sp_{(2)} + 6\mu_{\mathrm{A}}\left[2s(1-h) - \frac{\mu_{\mathrm{a}}}{s}\right], \tag{A77}$$

giving

$$p^*_{(2)} = 2\mu_{\mathrm{A}}\left[2(1-h) - \frac{\mu_{\mathrm{a}}}{s^2}\right] \quad \text{and} \quad C^*_{(2)} = -2\mu_{\mathrm{A}}\left(2h + \frac{\mu_{\mathrm{A}} + \mu_{\mathrm{a}}}{s^2}\right) \tag{A78}$$

at equilibrium. Thus, when selection is weak and mutations are rare,

$$p^* = \frac{\mu_{\mathrm{A}}}{s}\left(1 - \frac{\mu_{\mathrm{a}}}{s}\right) + 2(1-h)\mu_{\mathrm{A}} + \mathcal{O}\left(\epsilon^3\right) \quad \text{and} \quad C^* = \frac{\mu_{\mathrm{A}}}{s}\left(1 - \frac{\mu_{\mathrm{A}} + \mu_{\mathrm{a}}}{s}\right) - 2h\mu_{\mathrm{A}} + \mathcal{O}\left(\epsilon^3\right) \tag{A79}$$

under complete selfing.

**Class reproductive values and the basis to condition dependence.** Before turning out attention to selection gradients to order zero, let us first compute the zero-th order perturbations of asymptotic class frequencies $\mathbf{q}^{\circ}$ and reproductive values $\mathbf{v}^{\circ}$ under neutrality, $\mathbf{v}^{\circ}_{(0)}$ and $\mathbf{q}^{\circ}_{(0)}$, which we do by solving eq. (A51) with $\alpha_{\mathrm{AA}} = \alpha_{\mathrm{Aa}} = \alpha_{\mathrm{aa}} = 1$. This yields

$$\mathbf{q}^{\circ}_{(0)} = \big(0,\ 0,\ 0,\ 0,\ 1,\ 0,\ 0,\ 0\big) \tag{A80a}$$

and

$$\mathbf{v}^{\circ}_{(0)} = \left(\frac{1}{2},\ \frac{r}{1+2r},\ \frac{1}{2(1+2r)},\ 0,\ 1,\ \frac{1}{2},\ \frac{1}{2},\ 0\right). \tag{A80b}$$

Eq. (A80a) shows that a fully selfing resident population would be fixed for allele $\mathrm{A}$ to leading order, and any neutral mutant appearing at the selfing modifier would be exclusively found in homozygous state within the mutant sub-population.

It is interesting to compare the expression for $\mathbf{v}^{\circ}_{(0)}$ obtained here (eq. A80b) with the one we found in the complete outcrossing case (eq. A52). Under complete outcrossing, we found that all genotypic classes had a positive reproductive value to zero-th order, with classes that are homozygous for the mutant allele at the modifier having a reproductive value exactly twice that of heterozygous classes, because they transmit twice as many mutant copies under complete outcrossing. In contrast, eq. (A80) reveals that the zero-th order reproductive values of deleterious homozygote classes (classes 4 and 8) is zero under complete selfing. This is because in the absence of outcrossing, heterozygotes at the condition locus are very rare at equilibrium (indeed they only occur due to recurrent mutation) and homozygotes only produce descendants with the same genotype as them at the condition locus, so that wild-type and deleterious homozygotes effectively form independent, competing lineages. In such a setup, the contribution of deleterious homozygotes to the long-term fate of the population will be zero to leading order, because they suffer from a fecundity disadvantage and so will always be outcompeted by wild-type homozygotes.

More broadly, this means that any mutant allele found in linkage with the deleterious allele will asymptotically be lost unless it can recombine onto a wild-type background before being "trapped" in a deleterious homozygote. This is illustrated by the reproductive values of classes heterozygous at the condition locus.

Those classes that are heterozygous at the condition locus but homozygote for the mutant at the selfing modifier (classes 6 and 7) have a reproductive value exactly half that of a double homozygote carrying the wild-type (class 5), because half of their mutant copies will inevitably be associated with the deleterious allele and so make no asymptotic contribution to the future population. Meanwhile, the reproductive values of classes that are heterozygous at both loci (classes 2 and 3) depend on the recombination rate. Class 2, which corresponds to genotype $\mathrm{AM/am}$, carries a mutant allele at the modifier in linkage with the deleterious allele. Its long-term contribution to the mutant population is thus zero unless recombination allows it to produce $\mathrm{Am}$ haplotypes. As a result, its reproductive value is zero when $r = 0$ and increases with $r$. Conversely, class 3 individuals, which carry genotype $\mathrm{Am/aM}$, see their reproductive value maximised for $r = 0$ and decrease with $r$ because their mutant copy is already on the wild-type background. This deleterious background trap generates selection to escape it, which forms the basis to the evolution of condition-dependent self-fertilisation, as we will see below.

**Selection gradients to order zero.** We now compute selection gradients to order zero. The matrices of effects are identical to the complete outcrossing case to order zero, so using in eq. (A53), we find

$$S_{\mathrm{AA}}\left(\mathbf{1}_3\right) = \frac{1}{2} + \mathcal{O}\left(\epsilon\right), \tag{A81}$$

meaning that selection will always favour wild-type homozygotes to remain fully selfing when the rest of the population is fully selfing as well, and

$$S_{\mathrm{Aa}}\left(\mathbf{1}_3\right) = S_{\mathrm{aa}}\left(\mathbf{1}_3\right) = 0 + \mathcal{O}\left(\epsilon\right), \tag{A82}$$

so that higher order terms must be computed for other genotypes.

**Selection gradients to order $\epsilon$.** First of all, we note that

$$\mathbf{D}^{\mathrm{aa}}_{(0)} \cdot \mathbf{q}^{\circ}_{(0)} = \mathbf{D}^{\mathrm{Aa}}_{(0)} \cdot \mathbf{q}^{\circ}_{(0)} = \mathbf{0}, \tag{A83}$$

which implies that we need not compute the first order perturbation of reproductive values $\mathbf{v}^{\circ}_{(1)}$ for now. To obtain the first order perturbation of class frequencies $\mathbf{q}^{\circ}_{(1)}$, we solve eqs. (A56-A57) with $\alpha_{\mathrm{AA}} = \alpha_{\mathrm{Aa}} = \alpha_{\mathrm{aa}} = 1$, which yields

$$\mathbf{q}^{\circ}_{(1)} = \left(0, \quad 0, \quad 0, \quad 0, \quad -\frac{\mu_{\mathrm{A}}}{s}, \quad 0, \quad 0, \quad \frac{\mu_{\mathrm{A}}}{s}\right). \tag{A84}$$

Meanwhile, the matrices of effects differ from the complete outcrossing case to first order in $\epsilon$ and are given by

$$\mathbf{D}^{\mathrm{Aa}}_{(1)} = \begin{pmatrix} 0 & \frac{r\left(rs^2h+\mu_{\mathrm{A}}\right)}{4s} & \frac{(1-r)\left[(1-r)s^2h+\mu_{\mathrm{A}}\right]}{4s} & 0 & 0 & \frac{s^2h+\mu_{\mathrm{A}}}{2s} & \frac{s^2h+\mu_{\mathrm{A}}}{2s} & 0 \\ 0 & \frac{(1-r)\left(rs^2h+\mu_{\mathrm{A}}\right)}{4s} & \frac{r\left[(1-r)s^2h+\mu_{\mathrm{A}}\right]}{4s} & 0 & 0 & \frac{s^2h+\mu_{\mathrm{A}}}{2s} & \frac{s^2h+\mu_{\mathrm{A}}}{2s} & 0 \\ 0 & -\frac{r\left(rs^2h+\mu_{\mathrm{A}}\right)}{4s} & -\frac{(1-r)\left[(1-r)s^2h+\mu_{\mathrm{A}}\right]}{4s} & 0 & 0 & -\frac{\mu_{\mathrm{A}}}{2s} & -\frac{\mu_{\mathrm{A}}}{2s} & 0 \\ 0 & -\frac{(1-r)\left(rs^2h+\mu_{\mathrm{A}}\right)}{4s} & -\frac{r\left[(1-r)s^2h+\mu_{\mathrm{A}}\right]}{4s} & 0 & 0 & -\frac{\mu_{\mathrm{A}}}{2s} & -\frac{\mu_{\mathrm{A}}}{2s} & 0 \\ 0 & -\frac{shr^2}{8} & -\frac{sh(1-r)^2}{8} & 0 & 0 & -\frac{sh}{4} & -\frac{sh}{4} & 0 \\ 0 & -\frac{shr(1-r)}{8} & -\frac{shr(1-r)}{8} & 0 & 0 & -\frac{sh}{4} & -\frac{sh}{4} & 0 \\ 0 & -\frac{shr(1-r)}{8} & -\frac{shr(1-r)}{8} & 0 & 0 & -\frac{sh}{4} & -\frac{sh}{4} & 0 \\ 0 & -\frac{sh(1-r)^2}{8} & -\frac{shr^2}{8} & 0 & 0 & -\frac{sh}{4} & -\frac{sh}{4} & 0 \end{pmatrix}, \tag{A85a}$$

and

$$\mathbf{D}^{\mathrm{aa}}_{(1)} = \begin{pmatrix} 0 & 0 & 0 & 0 & 0 & 0 & 0 & 0 \\ 0 & 0 & 0 & \frac{1}{4}\left(s+\frac{\mu_{\mathrm{A}}}{s}\right) & 0 & 0 & 0 & s+\frac{\mu_{\mathrm{A}}}{s} \\ 0 & 0 & 0 & 0 & 0 & 0 & 0 & 0 \\ 0 & 0 & 0 & -\frac{1}{4}\left(s+\frac{\mu_{\mathrm{A}}}{s}\right) & 0 & 0 & 0 & -\frac{\mu_{\mathrm{A}}}{s} \\ 0 & 0 & 0 & 0 & 0 & 0 & 0 & 0 \\ 0 & 0 & 0 & 0 & 0 & 0 & 0 & 0 \\ 0 & 0 & 0 & 0 & 0 & 0 & 0 & 0 \\ 0 & 0 & 0 & -\frac{s}{8} & 0 & 0 & 0 & -s \end{pmatrix}. \tag{A85b}$$

Using eqs. (A83-A85), the selection gradients on $\alpha_{\mathrm{Aa}}$ and $\alpha_{\mathrm{aa}}$ to first order in $\epsilon$ are therefore given by

$$S_{\mathrm{Aa}}\left(\mathbf{1}_3\right) = 0 + \mathcal{O}\left(\epsilon^2\right) \quad \text{and} \quad S_{\mathrm{aa}}\left(\mathbf{1}_3\right) = -\frac{r}{1+2r}\frac{\mu_{\mathrm{A}}}{s} + \mathcal{O}\left(\epsilon^2\right) \tag{A86}$$

Eq. (A86) reveals that the selection gradient on $\alpha_{\mathrm{Aa}}$ is zero to first order in $\epsilon$, so that higher order terms must be computed. This is because the frequency of heterozygotes is of the order of the mutation rate under complete selfing, which is of order $\epsilon^2$. As for the selection gradient on $\alpha_{\mathrm{aa}}$, eq. (A86) shows that it is negative to leading order, indicating that selection favours deleterious homozygotes to reduce their selfing rate when the rest of the population is fully selfing. To understand this result, one must think from the point of view of an allele at the selfing modifier, keeping in mind that alleles linked to the deleterious allele at the condition locus are asymptotically lost. For such an allele, complete selfing in deleterious homozygotes 'traps' it onto a deleterious background and so dooms it to extinction, whereas outcrossing gives it a chance at escaping the trap by mating with wild-type individuals and recombining. Accordingly, the gradient is zero when there is no recombination ($r = 0$) as outcrossing then no longer constitutes a viable escape route, and increases with $r$.

**Selection gradients to order $\epsilon^2$.** To compute second order terms in $\epsilon$ (eq. A40), we first note that

$$\mathbf{D}^{\text{Aa}}_{(0)} \cdot \mathbf{q}^{\circ}_{(0)} = \mathbf{D}^{\text{Aa}}_{(1)} \cdot \mathbf{q}^{\circ}_{(0)} = \mathbf{D}^{\text{Aa}}_{(2)} \cdot \mathbf{q}^{\circ}_{(0)} = \mathbf{0}, \tag{A87}$$

as we show in the accompanying *Mathematica* notebook, so that we need not compute first and second order perturbations of $\mathbf{v}^{\circ}$, $\mathbf{v}^{\circ}_{(1)}$ and $\mathbf{v}^{\circ}_{(2)}$. We next compute the second order perturbation of $\mathbf{q}^{\circ}$, $\mathbf{q}^{\circ}_{(2)}$ by solving eqs. (A64-A65), which yields

$$\mathbf{q}^{\circ}_{(2)} = \left(0,\quad 0,\quad 0,\quad 0,\quad 2\mu_{\text{A}}\left(\frac{\mu_{\text{A}}}{s^2} - 2(2-h)\right),\quad 4\mu_{\text{A}},\quad 4\mu_{\text{A}},\quad -2\mu_{\text{A}}\left(\frac{\mu_{\text{A}}}{s^2} + 2h\right)\right) \tag{A88}$$

so that the selection gradient on $\alpha_{\text{Aa}}$ is given by

$$S_{\text{Aa}}\left(\mathbf{1}_3\right) = \frac{\mu_{\text{A}}}{1+2r} + \mathcal{O}\left(\epsilon^3\right) \tag{A89}$$

to leading order, demonstrating that selection favours complete selfing in heterozygotes.

##### A.2.4 Pre-existing positive condition dependence ($\alpha_{\text{aa}} < \alpha_{\text{Aa}} < \alpha_{\text{AA}}$)

Section A.2.3 shows that selection favours a decrease in the selfing rate in deleterious homozygotes as an escape from the trap that constitutes linkage to the deleterious allele under complete selfing. But how much should the selfing rate of these homozygotes be reduced, and does this reduction eventually drive a change in the direction of selection on the selfing rate of other genotypes?

In this section, we consider the evolution of condition-dependent selfing under the looser assumption that $\alpha_{\text{aa}} < \alpha_{\text{Aa}} < \alpha_{\text{AA}}$, which is a situation likely to be observed. Indeed, section A.2.2 demonstrates that selection favours an increase in all three genotype-specific selfing rates from complete outcrossing, but the strength of selection is proportional to the frequency of the three genotypes at the condition locus, and so

$$S_{\text{aa}}\left(\mathbf{0}\right) \ll S_{\text{Aa}}\left(\mathbf{0}\right) \ll S_{\text{AA}}\left(\mathbf{0}\right).$$

This indicates that alleles increasing $\alpha_{\text{AA}}$ are much more likely to fix than alleles increasing $\alpha_{\text{Aa}}$, which are themselves much more likely to fix than alleles increasing $\alpha_{\mathrm{aa}}$. Assuming the mutational input is identical and independent for all three traits, it follows that the selfing rate of wild-type homozygotes should increase much faster – in expectation – than that of heterozygotes, which in turn should increase much faster than that of deleterious homozygotes. That is, in a large population with rare mutations of small effect, we are likely to see

$$\alpha_{\mathrm{aa}} < \alpha_{\mathrm{Aa}} < \alpha_{\mathrm{AA}}, \tag{A90}$$

at least initially.

**Selection on the selfing rate of wild-type homozygotes, $\alpha_{\mathrm{AA}}$.** To order zero in $\epsilon$, the change in allelic frequency $\Delta p_{(0)}$ and in homozygosity $\Delta C_{(0)}$ at the condition locus depend on the $\boldsymbol{\alpha}$ in a complicated way, so we refrain from giving them here (but they are available in the *Mathematica* notebook accompanying this Appendix; `10.5281/zenodo.18523420`). Solving for equilibrium (eq. A7), we find that

$$p^*_{(0)} = C^*_{(0)} = 0, \tag{A91}$$

so long as eq. (A90) holds. This is because, even in the absence of mutation or fecundity selection, allele $\mathrm{A}$ enjoys a transmission advantage relative to allele $\mathrm{a}$ here, as it is found in genotypes self-fertilising at a higher rate on average, and so displaces allele $\mathrm{a}$ competitively. Using eq. (A91), we may compute the zero-th order perturbation of $\mathbf{q}^\circ$ and $\mathbf{v}^\circ$, $\mathbf{q}^\circ_{(0)}$ and $\mathbf{v}^\circ_{(0)}$, as the solutions of

$$\mathbf{W}^\circ_{(0)} \cdot \mathbf{q}^\circ_{(0)} = \mathbf{q}^\circ_{(0)} \quad \text{and} \quad \mathbf{v}^\circ_{(0)} \cdot \mathbf{W}^\circ_{(0)} = \mathbf{v}^\circ_{(0)}, \tag{A92}$$

with constraints $\mathbf{q}^\circ_{(0)} \cdot \mathbf{1}_3 = 1$ and $\mathbf{v}^\circ_{(0)} \cdot \mathbf{q}^\circ_{(0)} = 1$. These solutions are given in the accompanying *Mathematica* notebook. Using the matrix of effects $\mathbf{D}^{\mathrm{AA}}_{(0)}$ (which is identical to eq. (A53a)), we find that the selection gradient on $\alpha_{\mathrm{AA}}$ reduces to

$$S_{\mathrm{AA}}\left(()\,\boldsymbol{\alpha}\right) = \frac{1}{2(2-\alpha_{\mathrm{AA}})}, \tag{A93}$$

which depends only on $\alpha_{\mathrm{AA}}$ and is strictly positive. This shows that wild-type homozygotes will evolve a selfing rate $\alpha_{\mathrm{AA}} \to 1$ irrespective of the selfing rates at other genotypes (provided that eq. A90 holds). We next compute selection gradients on $\alpha_{\mathrm{Aa}}$ and $\alpha_{\mathrm{aa}}$ under the assumption that $\alpha_{\mathrm{AA}} = 1$.

**Selection on the selfing rate of heterozygotes, $\alpha_{\mathrm{Aa}}$.** Using the same approach as in sections A.2.2 and A.2.3 (eqs. A10-A49; see accompanying *Mathematica* notebook for the detailed approach), we find that the equilibrium allelic frequency and homozygosity in the resident population are given by

$$p^* = \frac{2\mu_{\mathrm{A}}\left[2(1-\alpha_{\mathrm{aa}}) + \alpha_{\mathrm{Aa}}\right]}{(2-\alpha_{\mathrm{Aa}})(1-\alpha_{\mathrm{aa}})} + \mathcal{O}\left(\epsilon^3\right) \quad \text{and} \quad C^* = \frac{2\mu_{\mathrm{A}}\alpha_{\mathrm{Aa}}}{(2-\alpha_{\mathrm{Aa}})(1-\alpha_{\mathrm{aa}})} + \mathcal{O}\left(\epsilon^3\right) \tag{A94}$$

to second order in $\epsilon$. These highlight the effect of the transmission advantage on genetic dynamics at the condition locus. The higher $\alpha_{\mathrm{aa}}$ and $\alpha_{\mathrm{Aa}}$, the lower the transmission advantage enjoyed by allele A over allele a, and the equilibrium frequency of allele a increases. In the absence of selfing in heterozygotes ($\alpha_{\mathrm{Aa}} = 0$), there is no excess homozygosity in the population ($C^* = 0$) to leading order, because there is then no way for a deleterious allele a arising through mutation to be transmitted in homozygous state.

After computing perturbations of $\mathbf{q}^\circ$ and $\mathbf{v}^\circ$ using the same approach as in sections A.2.2 and A.2.3 (see *Mathematica* notebook), we find that the selection gradient on the selfing rate of heterozygotes $\alpha_{\mathrm{Aa}}$ reduces to

$$S_{\mathrm{Aa}}(\boldsymbol{\alpha}) = \frac{\mu_{\mathrm{A}}}{2(2-\alpha_{\mathrm{Aa}})^2}\left[3 + \frac{2r(1-r)\alpha_{\mathrm{Aa}}}{2-\alpha_{\mathrm{Aa}}\big(1-2r(1-r)\big)}\right] + \mathcal{O}\left(\epsilon^3\right), \tag{A95}$$

which is strictly positive and does not depend on $\alpha_{\mathrm{aa}}$. Thus, selection favours the evolution of complete selfing in heterozygotes ($\alpha_{\mathrm{Aa}} \to 1$), similar to wild-type homozygotes.

**Selection on the selfing rate of deleterious homozygotes, $\alpha_{\mathrm{aa}}$.** To obtain the selection gradient on $\alpha_{\mathrm{aa}}$, we may now assume that $\alpha_{\mathrm{Aa}} = 1$. Terms of order $\epsilon^3$ must be computed, as all lower order terms are zero. However, we show in the accompanying *Mathematica* notebook that

$$\mathbf{v}^\circ_{(0)} \cdot \mathbf{D}^{\mathrm{aa}}_{(0)} = \mathbf{D}^{\mathrm{aa}}_{(0)} \cdot \mathbf{q}^\circ_{(0)} = \mathbf{D}^{\mathrm{aa}}_{(3)} \cdot \mathbf{q}^\circ_{(0)} = (0,\ 0,\ 0,\ 0,\ 0,\ 0,\ 0,\ 0), \tag{A96}$$

which implies that all terms involving third order perturbations will be zero. In fact, we find that the selection gradient on $\alpha_{\mathrm{aa}}$ simplifies to

$$\begin{aligned} S_{\mathrm{aa}}(\boldsymbol{\alpha}) &= \frac{\mathbf{v}_{(1)}^{\circ}\cdot\mathbf{D}_{(0)}^{\mathrm{aa}}\cdot\mathbf{q}_{(2)}^{\circ}}{2}+\mathcal{O}\left(\epsilon^{4}\right) \\ &= -\mu_{\mathrm{A}}\frac{s}{(1-\alpha_{\mathrm{aa}})^{2}}\frac{4r}{1+4r}\left[1-\frac{r(1-r)(1-\alpha_{\mathrm{aa}})}{[1+2r(1-r)](2-\alpha_{\mathrm{aa}})}\right]+\mathcal{O}\left(\epsilon^{4}\right) \end{aligned} \tag{A97}$$

which is strictly negative. Thus, although selection initially favours an increase in the selfing rate for all genotypes, deleterious homozygotes will always be selected towards a selfing rate of zero ($\alpha_{\mathrm{aa}}\to 0$) once the selfing rate of other genotypes is sufficiently high. In other words, selection favours the evolution of condition-dependent selfing, where high-condition individuals are completely self-fertilising and low-condition individuals are fully outcrossing.

##### A.2.5 Numerical analyses

To complement our analytical results, we investigated the evolution of condition-dependent selfing for an arbitrary initial relationship between $\alpha_{\mathrm{AA}}$, $\alpha_{\mathrm{Aa}}$ and $\alpha_{\mathrm{aa}}$ with numerical analyses in *Mathematica*. We used two different approaches, which we describe below.

**Iteration of evolutionary dynamics.** In the first approach, we assume that the population is initially fixed for an arbitrary selfing strategy $\boldsymbol{\alpha}_0=(\alpha_{\mathrm{AA}}^0,\alpha_{\mathrm{Aa}}^0,\alpha_{\mathrm{aa}}^0)$ and study how evolutionary dynamics proceed from there as follows. At any time step $n$, we start by determining the genetic equilibrium reached by a resident population expressing strategy $\boldsymbol{\alpha}_n=(\alpha_{\mathrm{AA}}^n,\alpha_{\mathrm{Aa}}^n,\alpha_{\mathrm{aa}}^n)$, which we do by iterating the dynamics of genotypic frequencies $\mathbf{q}_t=(q_{\mathrm{AA}}(t),q_{\mathrm{Aa}}(t),q_{\mathrm{aa}}(t))$ given by eq. A3 until an equilibrium is reached, i.e.,

$$\mathbf{1}_3\cdot(\mathbf{q}_{t+1}-\mathbf{q}_t)^2<\epsilon_{\mathrm{q}} \tag{A98}$$

where $\epsilon_{\mathrm{q}}>0$ is a small tolerance threshold. We then compute the selection gradient on the three traits at this equilibrium, $\mathrm{s}(\boldsymbol{\alpha}_n)=\left(S_{\mathrm{AA}}(\boldsymbol{\alpha}_n),S_{\mathrm{Aa}}(\boldsymbol{\alpha}_n),S_{\mathrm{aa}}(\boldsymbol{\alpha}_n)\right)$ using eq. (A38). The trait values expressed

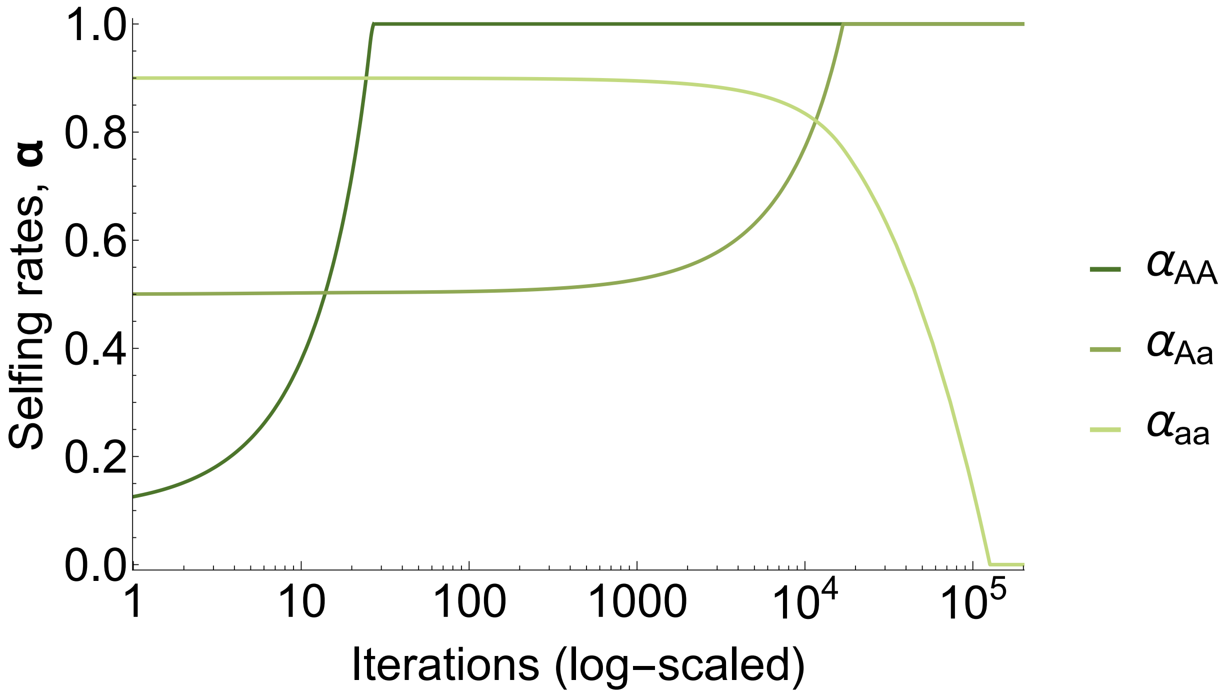

**Figure S1:** Evolution of condition-dependent selfing in the two locus model as a function of time (algorithm iterations) from initial negative condition dependence, $\boldsymbol{\alpha} = (0.1, 0.5, 0.9)$. Parameters used: $\mu_{\mathrm{A}} = \mu_{\mathrm{a}} = 10^{-3}$, $s = 0.05$, $h = 0.25$, $r = 0.5$.

in the population in the next time step are then given by

$$\boldsymbol{\alpha}_{n+1} = \boldsymbol{\alpha}_n + \Delta_\alpha \, \mathbf{s}(\boldsymbol{\alpha}_n), \tag{A99}$$

where $\Delta_\alpha > 0$ is a small phenotypic deviation and elements of $\boldsymbol{\alpha}_{n+1}$ are clipped to remain in the $[0, 1]$-interval. This algorithm is repeated until the change in trait between two time steps is sufficiently small, i.e.,

$$\mathbf{1}_3 \cdot (\boldsymbol{\alpha}_{n+1} - \boldsymbol{\alpha}_n)^2 < \epsilon_\alpha, \tag{A100}$$

where $\epsilon_\alpha > 0$ is a small tolerance threshold. Figure S1 illustrates the dynamics obtained using this approach.

**Gradient interpolation.** The approach given above can be greedy in computation time, as it is only valid for small phenotypic deviations and the selection gradients can be weak in some regions, leading to very small phenotypic changes from one iteration to the next. One way to circumvent this issue is to calculate the gradient for evenly spaced points in the $[0, 1]^3$ cube in which selfing strategies $\boldsymbol{\alpha}$ =

$(\alpha_{\mathrm{AA}}, \alpha_{\mathrm{Aa}}, \alpha_{\mathrm{aa}})$ are defined. For each point, we compute the equilibrium genotypic values in the resident population using eq. (A3) and then compute the selection gradient using eq. (A38), as above. The output of these calculations is stored to a file (see *Mathematica* notebook for technical details), and can be used to interpolate the selection gradient on $\boldsymbol{\alpha}$ in the entire phenotypic space. Given this interpolated gradient, we can then apply standard gradient descent methods to find the equilibrium strategy $\boldsymbol{\alpha}^* = (\alpha^*_{\mathrm{AA}}, \alpha^*_{\mathrm{Aa}}, \alpha^*_{\mathrm{aa}})$ favoured by selection.

**Evolution of condition dependence and the effect of back-mutation.** We used numerical analyses to investigate the evolution of condition-dependent selfing over a broad range of parameter values and found that a form of positive condition dependence where wild-type homozygotes and heterozygotes self-fertilise fully while deleterious homozygotes are purely outcrossing, i.e.,

$$\boldsymbol{\alpha}^* = (\alpha^*_{\mathrm{AA}}, \alpha^*_{\mathrm{Aa}}, \alpha^*_{\mathrm{aa}}) = (1,\ 1,\ 0) \tag{A101}$$

was favoured in almost all cases. However, we found that low recombination rates can sometimes prevent the evolution of condition dependence and lead to a pure selfing strategy instead ($\boldsymbol{\alpha}^* = \mathbf{1}_3$) because the benefits of outcrossing are low in this case, sufficiently so that the benefits reaped from self-fertilising when wild-type alleles are produced through back-mutation are larger. This is illustrated by the fact that selection favours the evolution of condition dependence for all $r \neq 0$ when there is no back-mutation ($\mu_{\mathrm{a}} = 0$), as can be seen from Fig. S2.

##### A.2.6 Fixed inbreeding depression component

In this section, we investigate the effect of a fixed inbreeding depression component on the evolution of condition-dependent selfing. We assume that this fixed inbreeding depression component acts at the seed stage: the population follows the same life cycle as before, expect that self-fertilised seeds become viable and join the competition for recruitment with probability $1 - \delta$, whereas the outcrossed to so with probability one. The parameter $\delta \in [0, 1]$ thus measures the strength of inbreeding depression.

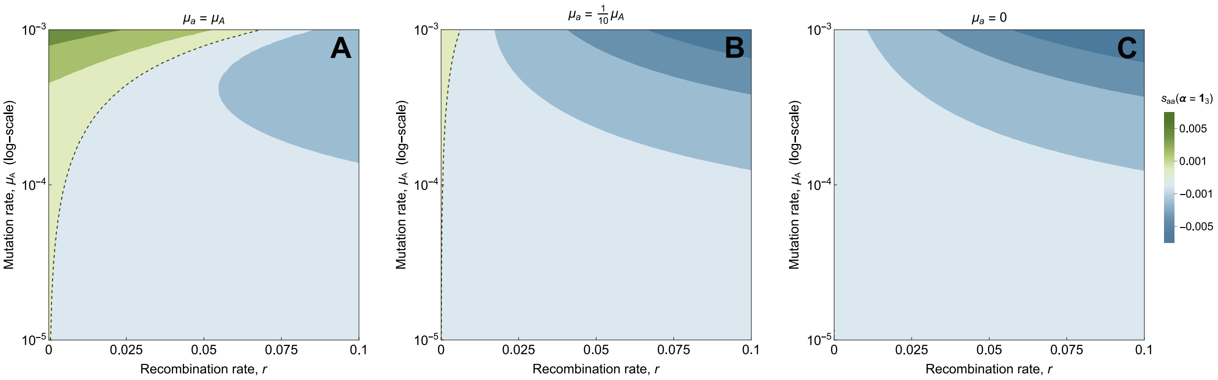

**Figure S2:** Effect of back-mutation on the evolution of condition-dependent selfing under low recombination rates. Plots give the value of the selection gradient on the selfing rate of deleterious homozygotes in a fully selfing population. Shades of blue indicate negative values, where condition dependence is favoured; shades of green indicate green values, where it is not. The dashed line indicates where the gradient changes sign. condition dependence is favoured for most of the parameter space, except when back-mutation is frequent and recombination very low. Parameters used: $s = 0.01$, $h = 0.25$.

With fixed inbreeding depression, the number of seeds produced by an individual depends directly on its selfing rate, which may vary depending on its genotype at both loci. The mean number of seeds produced by an individual in the resident population at equilibrium thus becomes

$$\overline{F}(\boldsymbol{\alpha}) = q^*_{\mathrm{AA}}\varphi_{\mathrm{AA}}\left(1 - \alpha_{\mathrm{AA}}\delta\right) + q^*_{\mathrm{Aa}}\varphi_{\mathrm{Aa}}\left(1 - \alpha_{\mathrm{Aa}}\delta\right) + q^*_{\mathrm{aa}}\varphi_{\mathrm{aa}}\left(1 - \alpha_{\mathrm{aa}}\delta\right), \tag{A102a}$$

and the relative number of ovules carrying allele A and a and available for outcrossing produced by residents, $F^*_{\mathrm{A}}(\boldsymbol{\alpha})$ and $F^*_{\mathrm{a}}(\boldsymbol{\alpha})$, now read

$$F^*_{\mathrm{A}}(\boldsymbol{\alpha}) = \frac{1}{\overline{F}(\boldsymbol{\alpha})}\left[(1-\mu_{\mathrm{A}})\, q^*_{\mathrm{AA}}\varphi_{\mathrm{AA}}(1-\alpha_{\mathrm{AA}}) + q^*_{\mathrm{Aa}}\frac{1-\mu_{\mathrm{A}}+\mu_{\mathrm{a}}}{2}\varphi_{\mathrm{Aa}}(1-\alpha_{\mathrm{Aa}}) + \mu_{\mathrm{a}}\, q^*_{\mathrm{aa}}\varphi_{\mathrm{aa}}(1-\alpha_{\mathrm{aa}})\right] \tag{A102b}$$

and

$$F^*_{\mathrm{a}}(\boldsymbol{\alpha}) = \frac{1}{\overline{F}(\boldsymbol{\alpha})}\left[\mu_{\mathrm{A}}\, q^*_{\mathrm{AA}}\varphi_{\mathrm{AA}}(1-\alpha_{\mathrm{AA}}) + q^*_{\mathrm{Aa}}\frac{1-\mu_{\mathrm{a}}+\mu_{\mathrm{A}}}{2}\varphi_{\mathrm{Aa}}(1-\alpha_{\mathrm{Aa}}) + (1-\mu_{\mathrm{a}})\, q^*_{\mathrm{aa}}\varphi_{\mathrm{aa}}(1-\alpha_{\mathrm{aa}})\right], \tag{A102c}$$

respectively. Using these notations, the number of offspring of class 1 to 8 produced by a focal mutant of class $i$ is given by

$$
\mathbf{w}_i(\boldsymbol{\alpha}, \boldsymbol{\alpha}_{\mathrm{m}}) = \begin{pmatrix}
\frac{\varphi_i}{\overline{F}(\boldsymbol{\alpha})}\left[2\alpha_i(1-\delta)G^{(i)}_{\mathrm{AM}}G^{(i)}_{\mathrm{Am}} + (1-\alpha_i)M^*_{\mathrm{A}}(\boldsymbol{\alpha})G^{(i)}_{\mathrm{Am}}\right] + F^*_{\mathrm{A}}(\boldsymbol{\alpha})\frac{\varphi_i}{\overline{\varphi}^*}G^{(i)}_{\mathrm{Am}} \\
\frac{\varphi_i}{\overline{F}(\boldsymbol{\alpha})}\left[2\alpha_i(1-\delta)G^{(i)}_{\mathrm{AM}}G^{(i)}_{\mathrm{am}} + (1-\alpha_i)M^*_{\mathrm{A}}(\boldsymbol{\alpha})G^{(i)}_{\mathrm{am}}\right] + F^*_{\mathrm{A}}(\boldsymbol{\alpha})\frac{\varphi_i}{\overline{\varphi}^*}G^{(i)}_{\mathrm{am}} \\
\frac{\varphi_i}{\overline{F}(\boldsymbol{\alpha})}\left[2\alpha_i(1-\delta)G^{(i)}_{\mathrm{Am}}G^{(i)}_{\mathrm{aM}} + (1-\alpha_i)M^*_{\mathrm{a}}(\boldsymbol{\alpha})G^{(i)}_{\mathrm{Am}}\right] + F^*_{\mathrm{a}}(\boldsymbol{\alpha})\frac{\varphi_i}{\overline{\varphi}^*}G^{(i)}_{\mathrm{Am}} \\
\frac{\varphi_i}{\overline{F}(\boldsymbol{\alpha})}\left[2\alpha_i(1-\delta)G^{(i)}_{\mathrm{am}}G^{(i)}_{\mathrm{aM}} + (1-\alpha_i)M^*_{\mathrm{a}}(\boldsymbol{\alpha})G^{(i)}_{\mathrm{am}}\right] + F^*_{\mathrm{a}}(\boldsymbol{\alpha})\frac{\varphi_i}{\overline{\varphi}^*}G^{(i)}_{\mathrm{am}} \\
\frac{\varphi_i}{\overline{F}(\boldsymbol{\alpha})}\alpha_i(1-\delta)G^{(i)}_{\mathrm{Am}}G^{(i)}_{\mathrm{Am}} \\
\frac{\varphi_i}{\overline{F}(\boldsymbol{\alpha})}\alpha_i(1-\delta)G^{(i)}_{\mathrm{Am}}G^{(i)}_{\mathrm{am}} \\
\frac{\varphi_i}{\overline{F}(\boldsymbol{\alpha})}\alpha_i(1-\delta)G^{(i)}_{\mathrm{Am}}G^{(i)}_{\mathrm{am}} \\
\frac{\varphi_i}{\overline{F}(\boldsymbol{\alpha})}\alpha_i(1-\delta)G^{(i)}_{\mathrm{am}}G^{(i)}_{\mathrm{am}}
\end{pmatrix}. \quad \text{(A103)}
$$

We used eq. (A103) to analyse the extended model numerically following the same procedure as detailed in section A.2.5 (see accompanying *Mathematica* notebook for details). Overall, we found that inbreeding depression lower than a half ($\delta < 1/2$) had little effect on the results, with the complete selfing favoured in wild-type homozygotes and heterozygotes and complete outcrossing favoured in deleterious homozygotes for most of the parameter space. However, we found that levels of inbreeding depression close to a half could lead heterozygotes to evolve complete outcrossing instead ($\boldsymbol{\alpha}^* = (1, 0, 0)$), as shown in Fig. S3 (see main text for interpretation).

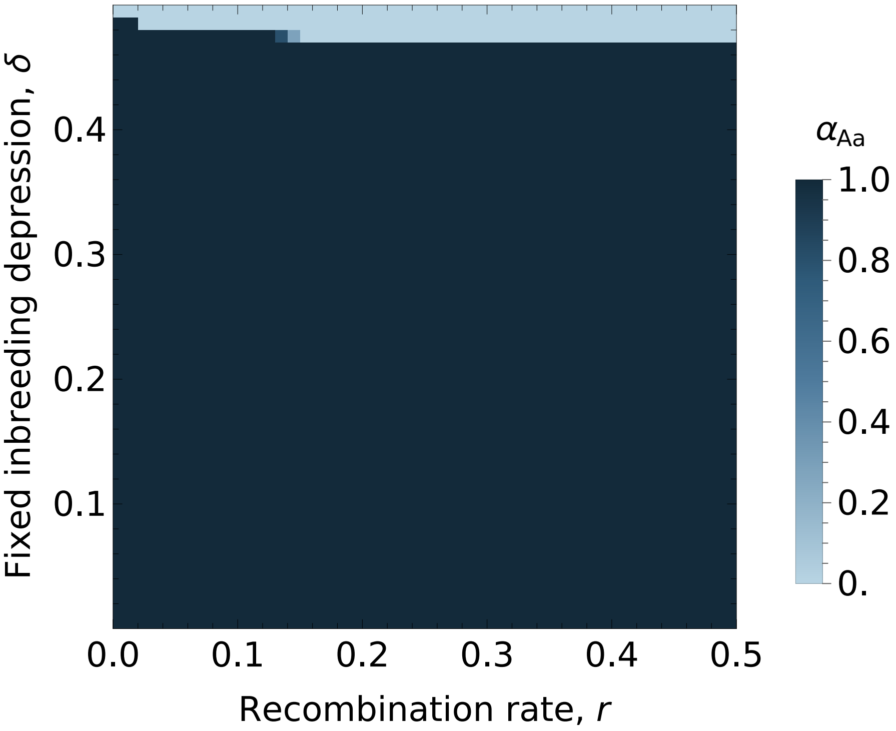

**Figure S3:** Effect of fixed inbreeding depression ($\delta$) and recombination ($r$) on the selfing rate of heterozygotes at the condition locus at evolutionary equilibrium.

### Appendix B

### Multilocus simulations

In this Appendix, we describe the simulation program and its extensions. The description given in this section focuses on the biological processes included in the simulation program. The program is coded in C++20 and available at `10.5281/zenodo.20510607`. We also provide additional results for parameter values not shown in the main.

#### B.1 The baseline program

The program simulates a population of $N$ individuals, where each of them is characterised by its diploid genotype at $L$ condition loci and at $n_{\mathrm{g}}$ network loci involved in a gene regulatory network that determines their selfing rate.

**Condition.** Condition loci are assumed to be diallelic, with a wild-type allele $\mathrm{A}_\ell$ and a deleterious allele $\mathrm{a}_\ell$ at each locus $l \in \{1, \dots, L\}$. These loci are assumed to be unlinked meaning that they all recombine with probability $1/2$ during meiosis, and each allele mutates with probability $\mu_{A_\ell} = \mu_{a_\ell} = \mu$ during gametogenesis, in which case they change state (i.e., the wild-type becomes a deleterious allele or the deleterious allele becomes wild-type). The deleterious allele at each locus reduces its bearer's condition by a proportion $s$ when homozygous and expresses proportionally to its dominance coefficient $h$ in heterozygotes. Condition loci act multiplicatively, so that the condition of an individual $i$ heterozygous at $n_{\mathrm{het}}^{(i)}$ loci and homozygous for the deleterious allele at $n_{\mathrm{hom}}^{(i)}$ loci is given by

$$\varphi_i = (1 - sh)^{n_{\mathrm{het}}^{(i)}} \times (1 - s)^{n_{\mathrm{hom}}^{(i)}}. \tag{B1}$$

**Selfing rate.** The gene regulatory network is composed of $n_{\mathrm{g}}$ loci and is inspired from the Wagner model (Wagner, 1994). Each locus $k \in \{1, ..., n_{\mathrm{g}}\}$ contains a protein-coding gene that is expressed as the individual develops into an adult and contributes to determining its selfing rate. The level of expression of each gene at developmental time step $t \in \{0, \dots, T\}$ in individual $i$, which we denote as $Z_k^{(i)}(t) \in \mathbb{R}$, is influenced by the combined action of individual condition $\varphi_i$, which acts as an external cue to the network, and regulatory interactions within the network (including self-regulation). Specifically, we assume that each locus additionally carries $n_{\mathrm{g}}$ cis-regulatory sequences on which the gene products of the $n_{\mathrm{g}}$ genes involved in the network may bind to up- or down-regulate the gene expression at locus $k$. Each locus $k$ is thus characterised by five attributes, an input weight $u_k^{(i)} \in \mathbb{R}$, which measures how much the expression of locus $k$ is directly influenced by condition; a slope $a_k^{(i)} \in \mathbb{R}$ and a bias $b_k^{(i)} \in \mathbb{R}$, which determine the shape of the response of locus $k$ to condition and regulation; a vector of regulatory weights $\mathbf{y}_k^{(i)} \in \mathbb{R}^{n_{\mathrm{g}}}$, the $m^{\mathrm{th}}$ element of which $y_{k,m}^{(i)}$ gives the regulatory effect of locus $m$ on the expression of locus $k$; and an output weight $v_k^{(i)}$ which measures the final contribution of locus $k$ to the phenotype. Alleles at each locus are additive, so that the attributes of locus $k$ in a diploid individual carrying alleles $k_1$ and $k_2$ at this locus are given by the average of the two alleles, i.e.,

$$x_k^{(i)} = \frac{x_{k_1} + x_{k_2}}{2}$$

for $x \in \{u, a, b, \mathbf{y}, v\}$. Each allele mutates with probability $\mu_{\mathrm{g}}$ during meiosis, in which case a normally distributed value with mean zero and standard deviation $\sigma_{\mathrm{g}}$ is added to each of its attributes.

At the beginning of development ($t = 0$), that is before any regulatory interaction, the level of expression of locus $k$ is determined by its sensitivity to individual condition, i.e.,

$$Z_k^{(i)}(0) = u_k^{(i)} \varphi_i. \tag{B2}$$

Then, given expression levels $\mathbf{Z}_i(t) = \left( Z_k^{(i)}(t) \right)_{1 \leqslant k \leqslant n_{\mathrm{g}}}$ at developmental time $t$, the expression level of locus $k$ at $t+1$ is given by

$$Z_k^{(i)}(t+1) = \frac{2}{1+\exp\left[-a_k^{(i)}\left(\mathbf{Z}_i(t)\cdot\mathbf{y}_k^{(i)} + u_k^{(i)}\varphi_i - b_k^{(i)}\right)\right]} - 1. \tag{B3}$$

The recursion given in eq. (B3) is iterated until expression levels reach a steady state $\mathbf{S}_i^* = \left( S_{k*}^{(i)} \right)_{1\leqslant k\leqslant n_\mathrm{g}}$, defined as

$$\mathbf{Z}_i^* = \mathbf{Z}_i(t) = \mathbf{Z}_i(t+1) \quad \text{such that} \quad \frac{1}{n_\mathrm{g}}\sum_{k=1}^{n_\mathrm{g}}\left(Z_k^{(i)}(t+1) - Z_k^{(i)}(t)\right)^2 < \epsilon_\mathrm{g}, \tag{B4}$$

where $0 < \epsilon_\mathrm{g} \ll 1$ is a small tolerance threshold, or the maximal development time $T$ is reached. Those individuals that fail to converge to steady expression levels as defined by eq. (B4) within $T$ iterations are assumed to be sterile owing to developmental instability (Wagner, 1994). Otherwise, the selfing rate expressed by the individual is given by

$$\alpha_i = \begin{cases} 0 \quad \text{if} \quad \mathbf{v}_i\cdot\mathbf{Z}_i^* < 0, \\ \mathbf{v}_i\cdot\mathbf{Z}_i^* \quad \text{if} \quad 0 \leqslant \mathbf{v}_i\cdot\mathbf{Z}_i^* \leqslant 1, \\ 1 \quad \text{if} \quad \mathbf{v}_i\cdot\mathbf{Z}_i^* > 1, \end{cases} \tag{B5}$$

where $\mathbf{v}_i = \left(v_k^{(i)}\right)_{1\leqslant k\leqslant n_\mathrm{g}}$ collects the output weights of the $n_\mathrm{g}$ loci involved in the network. Allowing the dot product $\mathbf{v}_i\cdot\mathbf{S}_i^*$ to vacate the $[0,1]$-interval in which the selfing rate is defined prevents the occurrence of directional mutational bias due to boundary effects.

**Initialisation.** The population is initially fixed for the wild-type allele at all $L$ condition loci and output weights $v_0$ at the $n_\mathrm{g}$ network loci. In this study, we set $v_0 = 0$ in all simulations to ensure that populations were initially purely outcrossing (see main text). To assess the sensitivity of our results to the initial state of the network, the input and regulatory weights, and the slope and bias of each trait locus were initialised by sampling values in Gaussian distributions with mean zero and standard deviation of $2\sigma_\mathrm{g}$. This led to initial networks with low sensitivity to condition and weak interactions among loci.

**Detailed life-cycle.** Each generation proceeds as follows. We first determine the condition $\varphi_i$ of each individual $i \in \{1, \dots, N\}$ using eq. (B1), and its selfing rate $\alpha_i$ by iterating its gene regulatory network as described above. If the network does not stabilise within $T$ developmental time steps, the condition of the individual is flagged with a selfing rate $\alpha_\bullet = -1$, which serves to identify it as sterile. From these, we calculate the female and male fecundities of individual $i$, $f_♀(\varphi_i, \alpha_i)$ and $f_♂(\varphi_i, \alpha_i)$, as

$$f_♀(\varphi_i, \alpha_i) = f_♂(\varphi_i, \alpha_i) = \begin{cases} \varphi_i & \text{if} \quad \alpha_i \geqslant 0, \\ 0 & \text{otherwise.} \end{cases} \tag{B6}$$

Fecundities through both sex functions are equal here, but we distinguish between the two functions now in anticipation of extensions introduced in the next section.

To create the next generation, we generate $N$ diploid offspring as follows. We first sample a maternal parent with replacement from the current generation. Each individual $i$ has a probability of being sampled proportional to its female fecundity, $f_♀(\varphi_i, \alpha_i)/\sum_j f_♀(\varphi_j, \alpha_j)$. This maternal parent then self-fertilises with probability $\alpha_i$, in which case the paternal parent is taken to be the same individual. Otherwise, a paternal parent $k \neq i$ is sampled from the current generation with a probability proportional to male fecundity, i.e., individual $k \neq i$ has a probability $f_♂(\varphi_k, \alpha_k)/\sum_{j\neq i} f_♂(\varphi_j, \alpha_j)$ to be sampled.

Each parents transmit one haplotype to the offspring. Recombination occurs freely between all loci, meaning that alleles on the transmitted haplotype are equally likely to come from each of the two chromosomes of the parent, independently for each of the $L$ condition loci and $n_{\text{g}}$ network loci. Mutation occurs with probability $\mu$ for each allele at condition loci, in which case the allele switches state (i.e., mutating wild-type alleles become deleterious, and mutating deleterious alleles revert to wild-type). Meanwhile, mutation occurs with probability $\mu_{\text{g}}$ for each allele at regulatory loci, in which case a normally distributed value with mean zero and standard deviation $\sigma_{\text{g}}$ is added to each of its attributes, i.e., network loci evolve under a continuum-of-alleles model (Kimura, 1965).

**Measurements and population back-up.** We let simulations run for $t_{\text{max}}$ generations. Every $t_{\text{mes}}$ generations, we record the following information.

First of all, we save the condition $\varphi_i$, selfing rate $\alpha_i$ and the number of generations of selfing since the last outcrossing event along an individual's genealogy of randomly sampled $n_{\text{mes}}$ individuals in the population. If enabled, we also record the GRN attributes of those individuals (this generates large output files and so should be used advisedly). In the extension that includes environmental fluctuations described in section B.2.1, we also record the environment they find themselves in.

Second, we record mean $\overline{\varphi}$ and variance $\sigma^2_\varphi$ in condition in the population, the magnitude of inbreeding depression $\delta$, which we compute as

$$\delta = \frac{\overline{\varphi_{\text{out}}} - \overline{\varphi_{\text{self}}}}{\overline{\varphi_{\text{out}}}}, \tag{B7}$$

where $\overline{\varphi_{\text{self}}}$ and $\overline{\varphi_{\text{out}}}$ are the mean condition of $N$ selfed and $N$ outcrossed individuals generated from randomly chosen parents, respectively; the mean $\overline{\alpha}$ and variance $\sigma^2_\alpha$ in selfing rate in the population, the number of deleterious alleles per haploid genome at the condition loci $n_{\text{mut}}$, and the mean excess in homozygote $F$ at condition loci relative to the Hardy-Weinberg expectation.

Third, we record the distribution of selfing rates and condition in the population with bins of size $0.01$ and $0.001$, respectively. Finally, every $t_{\text{save}}$ generations, we back-up the entire population into a single file, which can then be used to relaunch simulations from the latest back-up (or indeed any back-up with the same number of individuals and loci).

#### B.2 Extensions

We extended our baseline simulation program to include two additional biological mechanisms, namely environmental fluctuations and pollen discounting. We describe these extensions in turn below.

##### B.2.1 Environmental fluctuations

To include environmental fluctuations, we assume that each patch is characterised by an environmental value, $\varepsilon$, which we take to be a normally distributed random variable

$$\varepsilon \sim \mathcal{N}(0,1), \tag{B8}$$

that changes every generation. This value could represent any environmental variable relevant to plant development, such as exposure to sunlight or water availability. We assume that an environmental value $\varepsilon = 0$ is optimal and so maximises condition, while values deviating from zero result in lower condition. Specifically, the condition of an individual $i$ heterozygous for $n_{\text{het}}^{(i)}$ deleterious alleles and homozygous for $n_{\text{hom}}^{(i)}$, developing in a patch with value $\varepsilon_i$, is given by

$$\varphi_i = \exp\left(-\Delta_{\text{E}}\frac{\varepsilon_i^2}{2}\right)\left(1-s\right)^{n_{\text{hom}}^{(i)}}\left(1-sh\right)^{n_{\text{het}}^{(i)}} \tag{B9}$$

Eq. (B9) is composed of three terms. The first term describes the effect that the environment has on individual condition. This environmental component is maximised for $\varepsilon_i = 0$, i.e., when the individual develops in optimal environmental conditions, and decreases as $\varepsilon_i$ moves away from zero. Parameter $\Delta_{\text{E}}$ controls how fast condition decreases as the environment deviates from the optimum. The next two terms correspond to the effect of deleterious mutations in homozygous and heterozygous state on condition, as before.

##### B.2.2 Pollen discounting

To include pollen discounting in our model, that is, the reduction in pollen export that can accompany increased selfing, we modify our male fecundity function $f_{♂}(\varphi_i, \alpha_i)$, such that it is now given by

$$f_{♂}(\varphi_i, \alpha_i) = \begin{cases} \varphi_i\left(1-\kappa{\alpha_i}^{\gamma}\right) & \text{if} \quad \alpha_i \geqslant 0, \\ 0 \quad \text{otherwise.} \end{cases} \tag{B10}$$

where $\kappa \in [0,1]$ and $\gamma > 0$ respectively give the intensity and shape of the trade-off between selfing and pollen export. This functional shape is inspired from Johnston (1998), who showed that non-linear pollen discounting effects can maintain mixed mating. A linear relationship is obtained with $\gamma = 1$, as commonly assumed in the mating system literature (e.g., Abu Awad and Roze (2020)); and becomes non-linear as soon as $\gamma \neq 1$. The absence of pollen discounting is obtained for $\kappa = 0$, in which case we recover the baseline model.

#### B.3 Supplementary figures for environmental fluctuations

This section shows figures for all the simulations we ran for our environmental fluctuations extension.

### No environmental fluctuations, $\Delta_\mathrm{E} = 0$

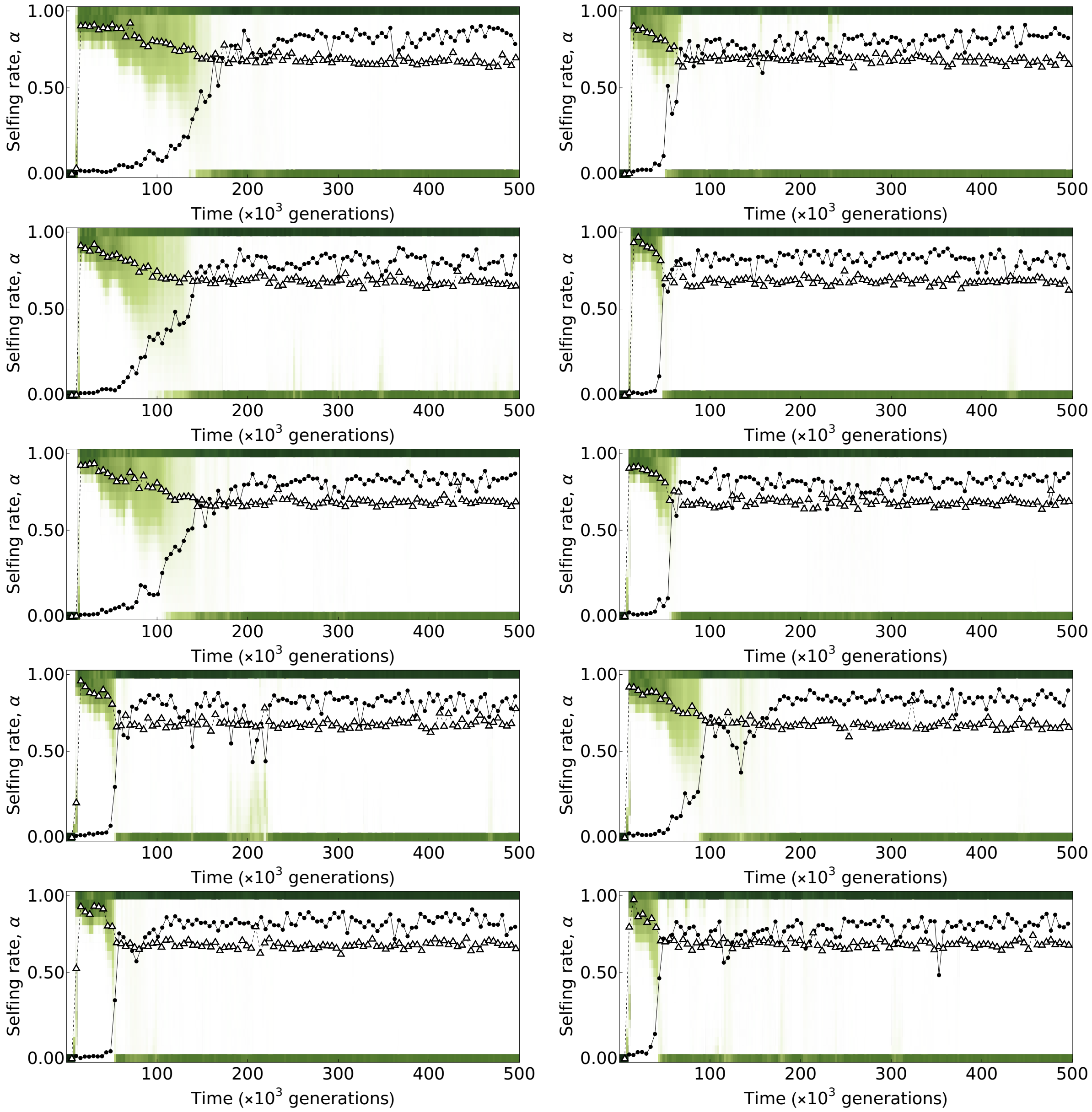

**Figure S4:** Simulation trajectories in the absence of environmental fluctuations ($\Delta_\mathrm{E} = 0$). As in the main text, green tiles indicate the proportion of individuals with a selfing rate in the corresponding range, and white triangles and black points respectively indicate mean and variance in the selfing rate. All simulations evolve condition-dependent selfing. Parameters used: $N = 5 \times 10^3$, $L = 10^3$, $\mu = 5 \times 10^{-4}$, $s = 0.05$, $h = 0.25$, $\mu_\mathrm{g} = 5 \times 10^{-3}$, $\sigma_\mathrm{g} = 0.05$.

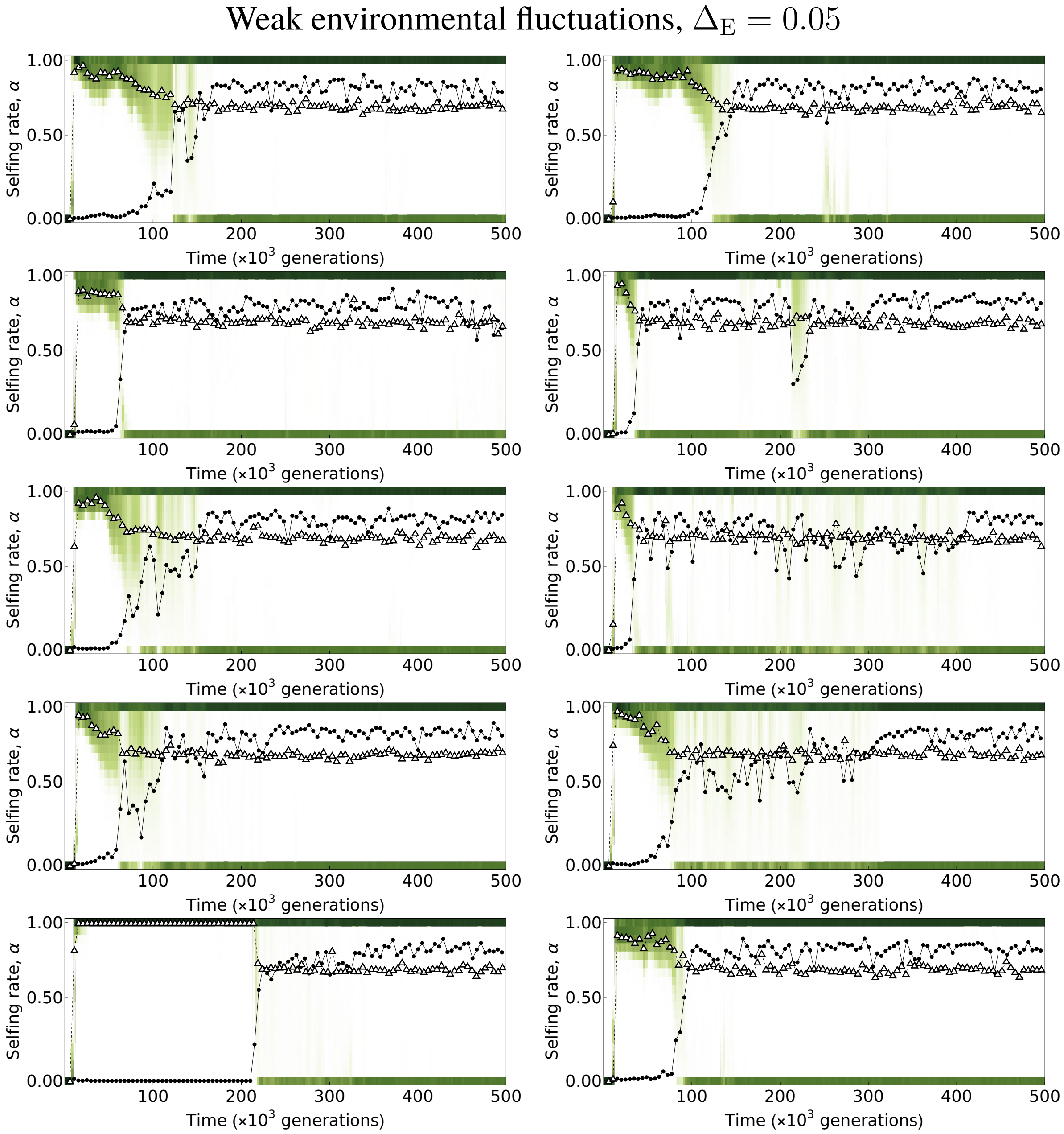

**Figure S5:** Simulation trajectories with weak environmental fluctuations ($\Delta_E = 0.05$). As in the main text, green tiles indicate the proportion of individuals with a selfing rate in the corresponding range, and white triangles and black points respectively indicate mean and variance in the selfing rate. All simulations evolve condition-dependent selfing. Parameters used: $N = 5 \times 10^3$, $L = 10^3$, $\mu = 5 \times 10^{-4}$, $s = 0.05$, $h = 0.25$, $\mu_g = 5 \times 10^{-3}$, $\sigma_g = 0.05$.

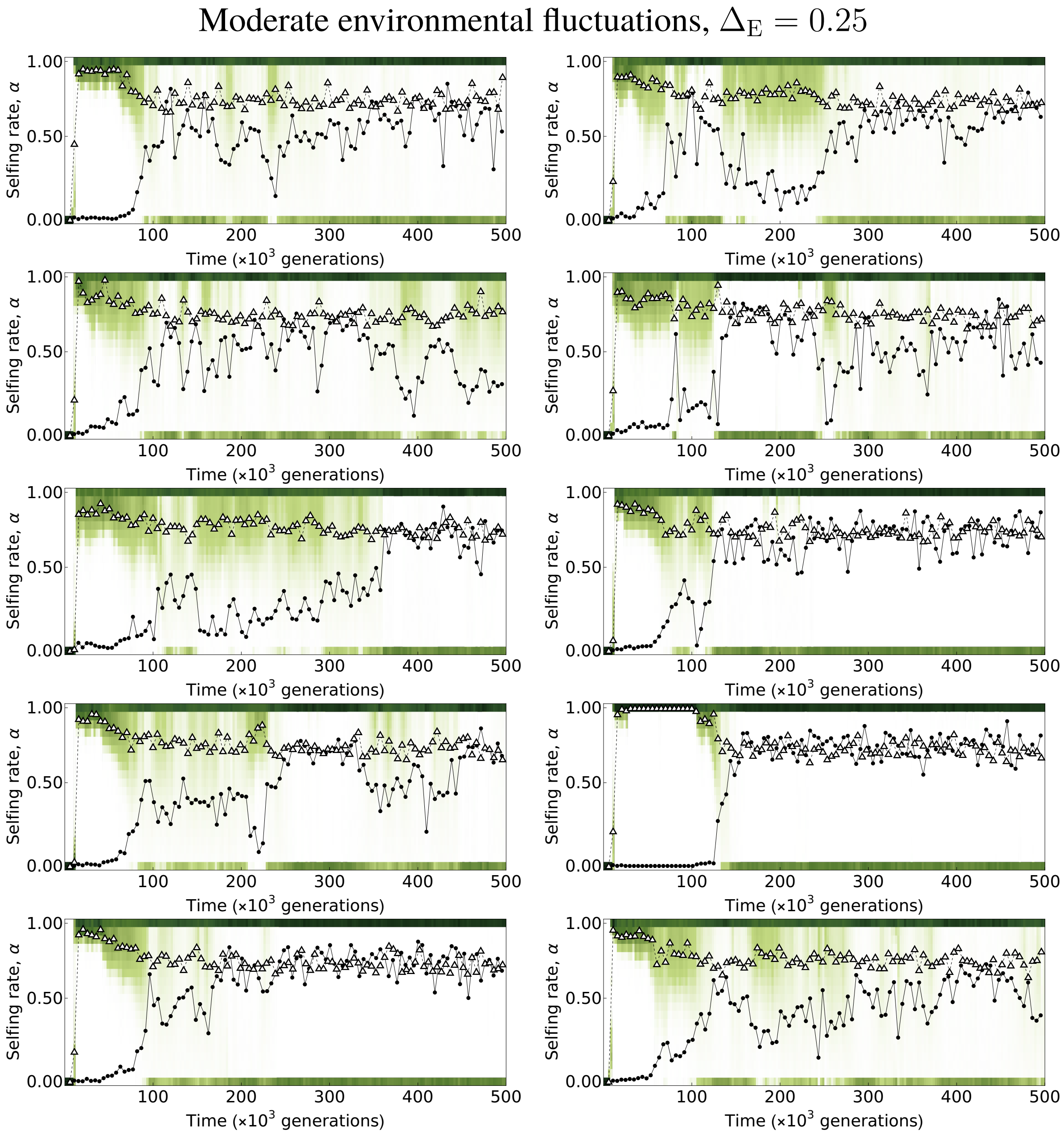

**Figure S6:** Simulation trajectories with moderate environmental fluctuations ($\Delta_E = 0.25$). As in the main text, green tiles indicate the proportion of individuals with a selfing rate in the corresponding range, and white triangles and black points respectively indicate mean and variance in the selfing rate. All simulations evolve condition-dependent selfing. Parameters used: $N = 5 \times 10^3$, $L = 10^3$, $\mu = 5 \times 10^{-4}$, $s = 0.05$, $h = 0.25$, $\mu_g = 5 \times 10^{-3}$, $\sigma_g = 0.05$.

Strong environmental fluctuations, $\Delta_E = 0.5$

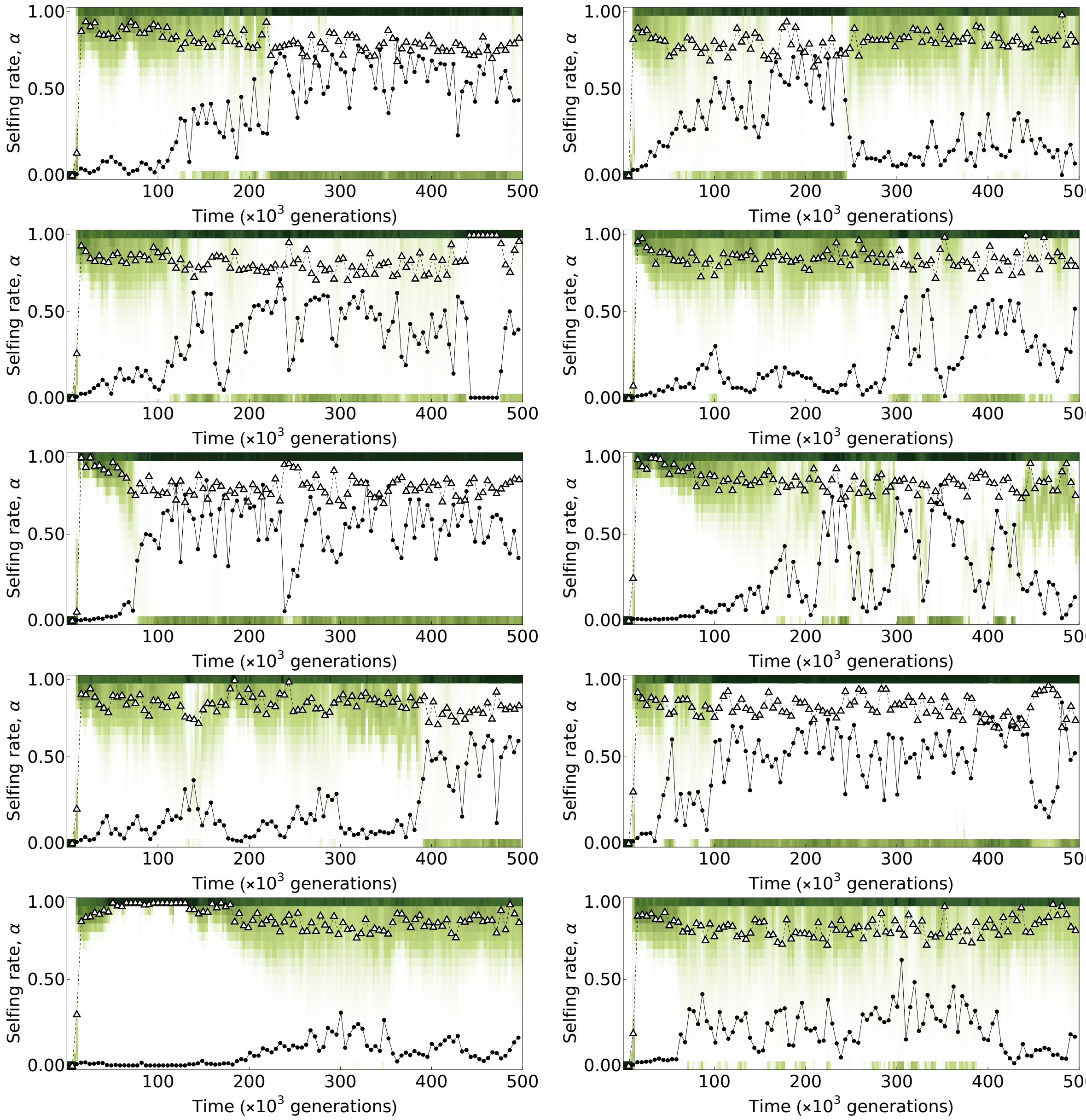

**Figure S7:** Simulation trajectories with strong environmental fluctuations ($\Delta_E = 0.5$). As in the main text, green tiles indicate the proportion of individuals with a selfing rate in the corresponding range, and white triangles and black points respectively indicate mean and variance in the selfing rate. All simulations evolve condition-dependent selfing. Parameters used: $N = 5 \times 10^3$, $L = 10^3$, $\mu = 5 \times 10^{-4}$, $s = 0.05$, $h = 0.25$, $\mu_g = 5 \times 10^{-3}$, $\sigma_g = 0.05$.

### Very strong environmental fluctuations, $\Delta_E = 1$

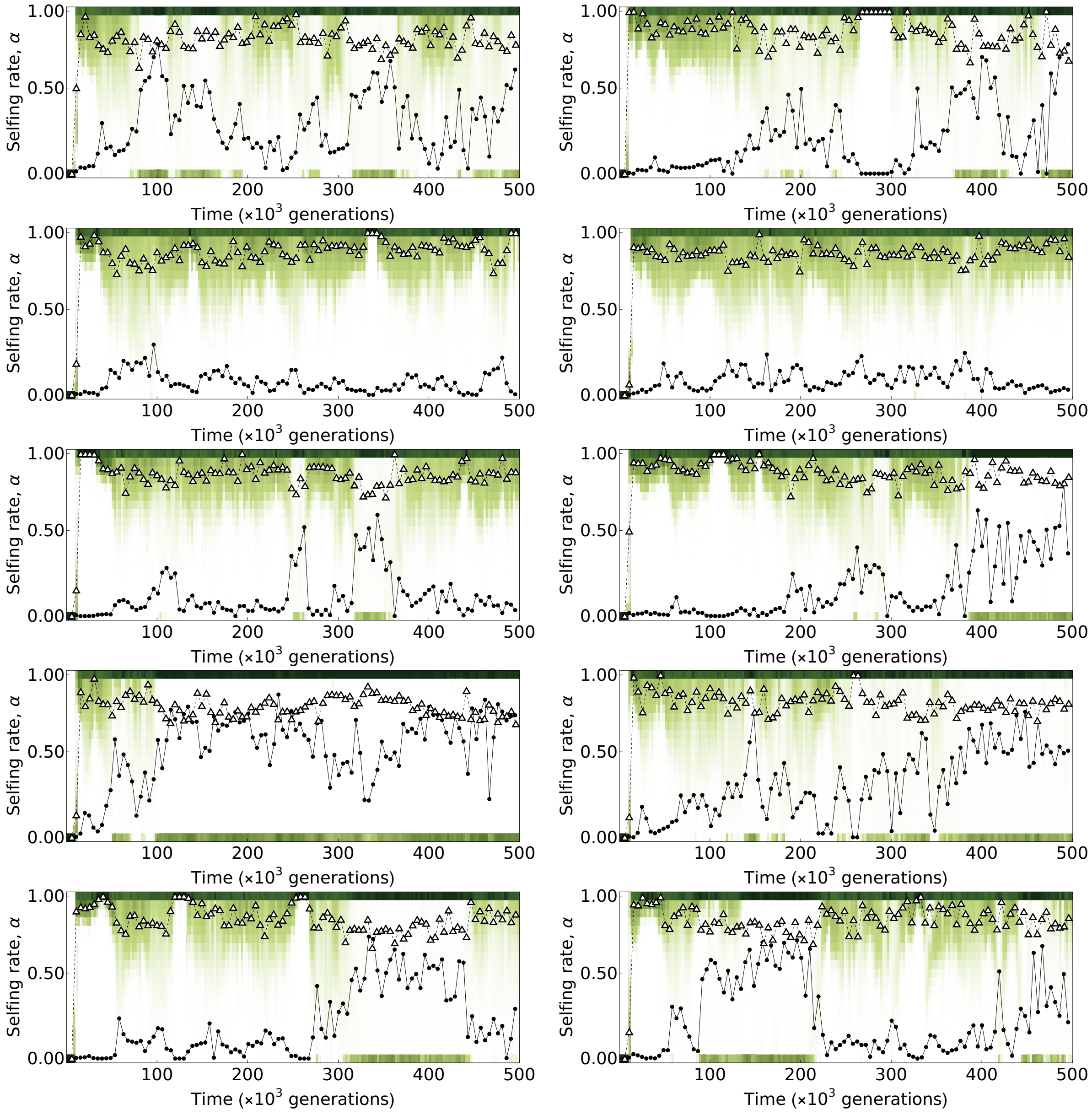

**Figure S8:** Simulation trajectories with very strong environmental fluctuations ($\Delta_E = 1$). As in the main text, green tiles indicate the proportion of individuals with a selfing rate in the corresponding range, and white triangles and black points respectively indicate mean and variance in the selfing rate. All simulations evolve condition-dependent selfing. Parameters used: $N = 5 \times 10^3$, $L = 10^3$, $\mu = 5 \times 10^{-4}$, $s = 0.05$, $h = 0.25$, $\mu_g = 5 \times 10^{-3}$, $\sigma_g = 0.05$.
